# Massively Parallel Selection of NanoCluster Beacons

**DOI:** 10.1101/2021.12.04.471212

**Authors:** Yu-An Kuo, Cheulhee Jung, Yu-An Chen, Hung-Che Kuo, Oliver S. Zhao, Trung D. Nguyen, James R. Rybarski, Soonwoo Hong, Yuan-I Chen, Dennis C. Wylie, John A. Hawkins, Jada N. Walker, Samuel W. Shields, Jennifer S. Brodbelt, Jeffrey T. Petty, Ilya J. Finkelstein, Hsin-Chih Yeh

## Abstract

NanoCluster Beacons (NCBs) are multicolor silver nanocluster probes whose fluorescence can be activated or tuned by a proximal DNA strand called the activator. While a single-nucleotide difference in a pair of activators can lead to drastically different activation outcomes, termed the polar opposite twins (POTs), it is difficult to discover new POT-NCBs using the conventional low-throughput characterization approaches. Here we report a high-throughput selection method that takes advantage of repurposed next-generation-sequencing (NGS) chips to screen the activation fluorescence of ∼40,000 activator sequences. We find the nucleobases at positions 7-12 of the 18-nucleotide-long activator are critical to creating bright NCBs and positions 4-6 and 2-4 are hotspots to generate yellow and red POTs, respectively. Based on these findings, we propose a “zipper bag model” that explains how these hotspots lead to the creation of distinct silver cluster chromophores and contribute to the difference in chromophore chemical yields. Combining high-throughput screening with machine learning algorithms, we establish a pipeline to rationally design bright and multicolor NCBs.

---

Activatable and multicolor fluorescent probes are indispensable tools in analytical chemistry and quantitative biology as they enable sensitive detection of analytes and diagnostic imaging of biomarkers in complex environments^1,2^. Whereas activatable probes have greatly simplified the assays by eliminating the need to remove unbound probes, the development of new activatable probes is largely constrained by the scarce activation mechanisms (e.g., FRET), the limited activation colors (e.g., existing FRET pairs) and the poor enhancement ratios (e.g., 10-to 60-fold for a typical molecular beacon)^3^. NanoCluster Beacons (NCBs)^4^ are a unique class of activatable probes as they provide a palette of activation colors from the same dark origin^5^ (not via FRET) and achieve fluorescence enhancement ratios as high as 1,500^6^ to 2,400-fold^7^. The core of NCB is a few-atom silver nanocluster^8-13^ (e.g., Ag_8_, Ag_10_ or Ag_16_) whose fluorescence can be tuned by its surrounding nucleobases^9,10,14-19^. To create an NCB, a dark AgNC is first synthesized in a C-rich DNA host (termed the NC probe), and a G-rich overhang (termed the activator) is brought into close proximity of the AgNC (via target-probe binding, **Supplementary Fig. S1**) to activate its fluorescence (**Fig. 1a-b**)^4-6,14,17^. Being a low-cost probe that can be easily prepared in a single-pot reaction at room temperature^8^, NCBs have been applied to the detection of nucleic acids^14,17,20^, proteins^21^, small molecules^22^, enzyme activities^16^ and cancer cells^23^.

**Fig. 1.**
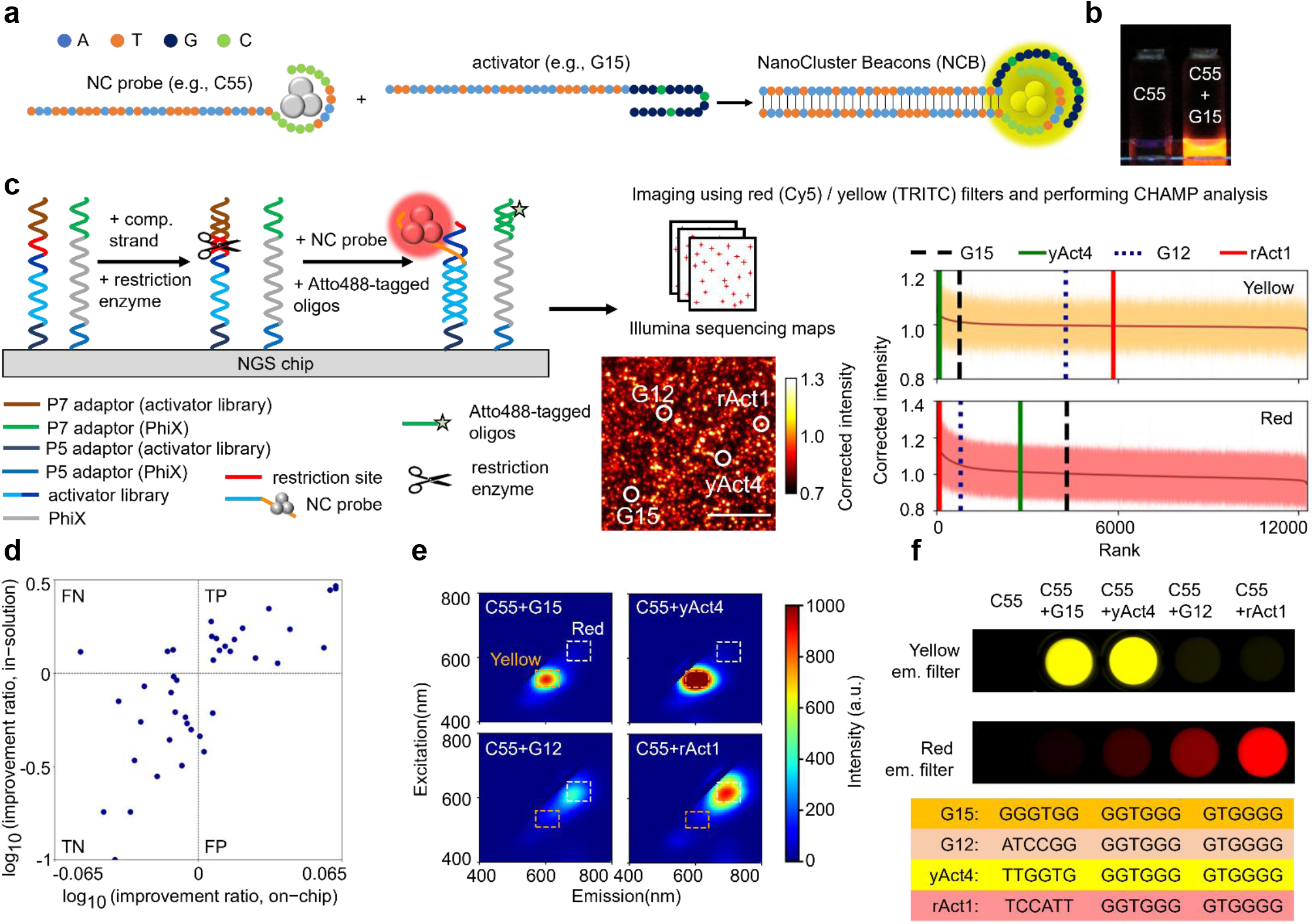
Massively parallel selection of NanoCluster Beacons (NCBs) using *MiSeq* chips. **a** The interactions between a silver nanocluster (AgNC, left) and a proximal guanine-rich activator (middle) activate the fluorescence of AgNC by hundreds to thousands fold, creating an activated NCB (right). Here a common C55 nanocluster (NC) probe is used for NCB selection and optimization. G15 is the canonical activator for yellow NCB. **b** C55 NC probe before and after activation by G15 activator, under UV excitation (365 nm). **c** Workflow of our high-throughput NCB selection on a next-generation sequencing chip (NGS; *MiSeq*, Illumina). After sequencing a library of activators (> 12,000) on the *MiSeq* chip, unwanted sequence above the activator was cleaved by a restriction enzyme. The Atto488-tagged fiducial marker probes and the C55 probes were then injected into the chip to hybridize with the PhiX markers and the library, and imaged sequentially under an epi-fluorescence microscope. A custom bioinformatics and imaging processing pipeline was employed to identify activator sequence behind each activated NCB spot. After ranking all activators based on their median activation brightness, we could clearly differentiate strong activators from weak ones in the yellow (Ex/Em: 535/50, 605/70 nm) and red (Ex/Em: 620/60, 700/75 nm) emission channels. Here G15 and G12 were the standards (the known best) for the yellow and red NCB comparisons, respectively. Both G15 and G12 ranking ∼800 among the yellow and red NCBs in this library (library_1). **d** Twenty top-ranked and twenty bottom-ranked activators were further investigated in test tubes using traditional florometry. The *MiSeq* results were 85% accurate in both true positive (TP) and true negative (TN) selections. Definition of the improvement ratio can be found in the methods. **e** 2D spectra of the four representative NCBs in the yellow (orange dashed box) and red (white dashed box) emission channels. Through florometry characterization, we found yAct4 2.03-fold brighter than G15 and rAct1 2.94-fold brighter than G12. Intensities were calculated based on a volumetric integral shown in **Supplementary Fig. S2. f**. Plate-reader images acquired using yellow (top) and red (bottom) excitation/emission filter sets, and the sequences of the four representative bright NCBs.

Whereas new applications of NCBs are emerging across a broad range of disciplines, it is unclear what sequence features of the activators ultimately control the enhancement ratio and activation color of an NCB. To answer this fundamental question and to unleash the power of NCB in biosensing, we design and study “polar opposite twin” NCBs (hereafter, denoted as POT-NCBs). Polar opposite twins are similar in appearance, but with very different personalities. In NCBs, polar opposite twins refer to a pair of NCBs that differ only by a single nucleotide in their activators, but have drastically distinct activation intensities or colors. Whereas POTs hold the key to understanding the NCB activation processes, there is no effective way to rapidly scan the vast activator sequence space and identify the most extreme POT-NCBs.

Here we repurpose the next-generation sequencing (NGS) chips for high-throughput screening of fluorescent nanomaterials. In a single experiment, more than 10^4^ activator mutations can be evaluated based on their capabilities in fluorescence activation of a common NC probe (C55 in **Fig. 1a**). Although the fluorescence properties of tens to hundreds of silver nanocluster species templated in short DNA strands can be studied in DNA microarrays^24^ and robotic plates^15^, less than 3,000 DNA hosts have been investigated as of today using these methods. While NGS chips are repurposed for studying protein-nucleic acid interactions^25-29^, they have never been used for study, selection, and optimization of fluorescent nanomaterials. By screening more than 40,000 activator sequences on three Illumina *MiSeq* chips, we not only discover new NCBs that are brighter than the known best (G15 and G12) but also identify the positions of nucleobases that are key to stabilizing bright AgNC chromophores (termed the critical zone). In the search for the most extreme POTs, the chip platform helps pinpoint the single-nucleotide substitution hotspots for generating yellow and red POT-NCBs, reaching 31-fold and 9-fold differences in the enhancement ratios, respectively (563 vs. 18 for a pair of yellow POTs and 285 vs. 32 for a pair of red POTs). Based on the findings of the critical zone for hosting bright chromophores (positions 7-12) and the hotspots for generating POTs (positions 4-6 for yellow POTs and positions 2-4 for red POTs), we propose a “zipper bag model” that explains how POT hotspots lead to the creation of distinct AgNC chromophores and contribute to the difference in chromophore chemical yields. In addition, with proper selection of the sequence features, we build machine learning models that can rationally design yellow and red NCBs. NCBs designed using these tools are 8.5 times and twice more likely to be bright yellow and red, respectively, as compared to the ones with random activator sequences. From our models, we create a new yellow NCB (with activator GTGTTGGGTGGTCGGGGG) that is twice as bright as the yellow standard G15 NCB, and a new red NCB (with activator ATCCCTCGGGGAGGGGGC) that is 1.3-fold brighter than the red standard G12 NCB. Our high-throughput screening and machine-learning-based design pipeline is not only accelerating the discovery of new NCBs for diverse applications, but also providing insights into the chemical yield and the emitter brightness controlled by the sequence features.

## Results

### High-throughput selection of red and yellow NCB on NGS chips

Our activator libraries were designed by systematically randomizing the canonical 18-nt-long activator G15^4,5,14,17^ (<u>GGGTGG GGTGGG GTGGGG</u>, **Fig. 2a** and **Supplementary Table S1**). Together with the fiducial markers (PhiX), the library sequences were immobilized, bridge amplified and sequenced on each of the Illumina *MiSeq* chips. As the sequencing-needed barcodes and adapters (i.e., SP2/barcode/P7 adapter, **Supplementary Table S1**) could suppress or alter the NCB fluorescence (**Supplementary Fig. S6-S8**), they were removed using a restriction enzyme, leaving behind 20-nt-long activators (the library) that were only 2 nt (CG dinucleotides) longer than the activators used in the traditional low-throughput test-tube selections^4,5,14,17^ (**Fig. 1c** and **Supplementary Table S1**). After enzymatic cleavage, the quality of the library was checked by staining the library with an Atto647N-labeled probe, before using the library for NCB selections (**Supplementary Fig. S6**).

**Fig. 2.**
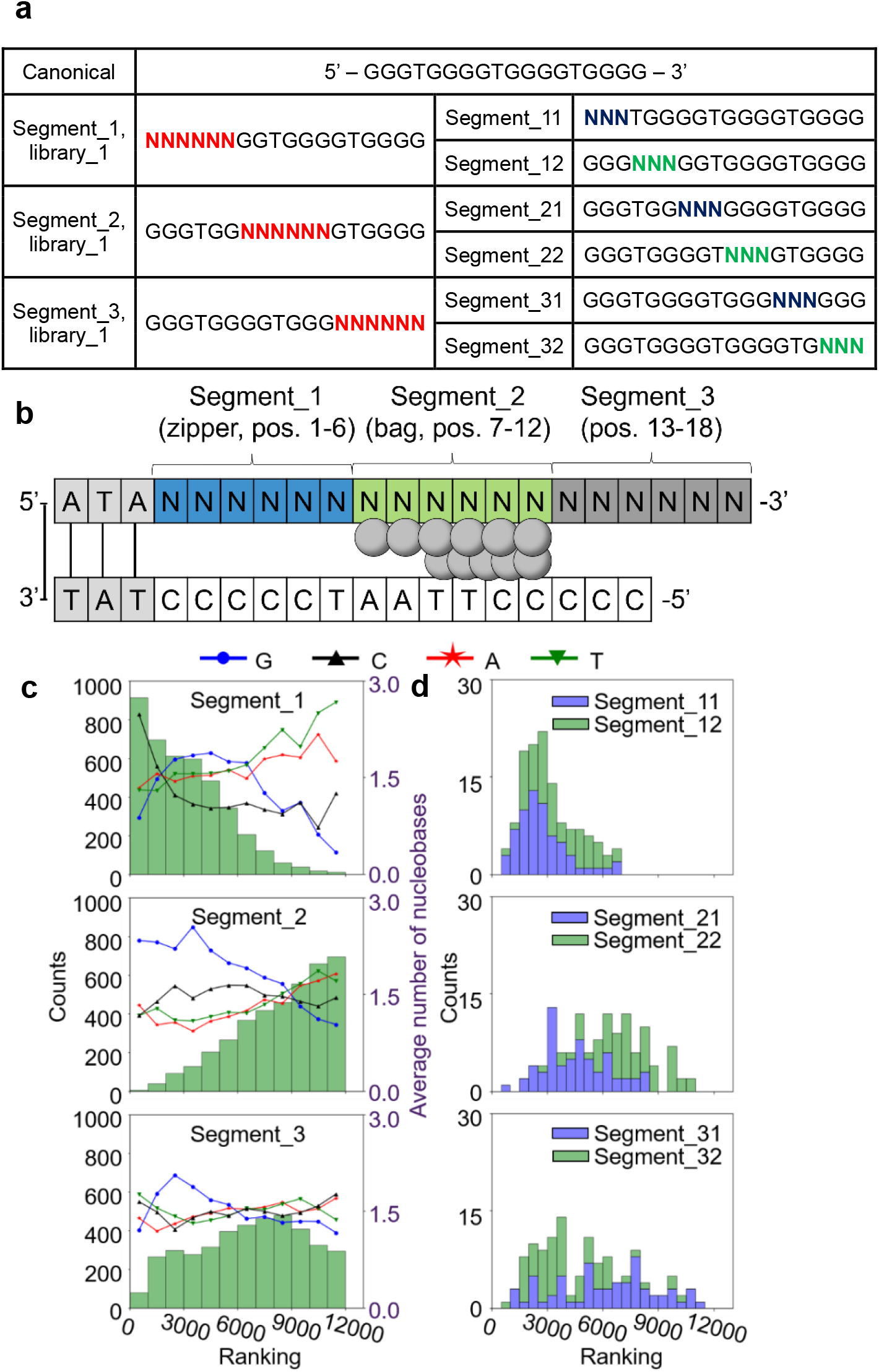
Influence of activator mutations on red NCB brightness. **a** In library_1, the 18-nt-long canonical activator G15 was divided into three 6-nt-long segments and each segment was separately randomized, creating a library with total 12,286 activators. **b** Schematic of NCB construct and definition of nucleobase positions in the activator. **c** Histograms of the brightness rankings corresponding to the mutated segments and the average numbers of the 4 nucleobases in the mutated segments. The library_1 results clearly indicated that, to make a bright NCB, segment_2 (the middle 6 nucleobases, positions 7-12) prefers the canonical G-rich sequence, as randomizing segment_2 (while keeping segment_1 and _3 canonical) leads to many low-ranking NCBs in both emission channels. Each histogram contained 4,096 activators. **d** The 6-segment interrogation further demonstrated that segment_22 (positions 10-12) is more important than segment_21 (positions 7-9) in creating bright red NCBs.

We first aimed to discover new activators that light up the common C55 NC probe^5^ (<u>CCCCC</u>TTAAT<u>CCCCC</u>, which hosts a dark AgNC) more intensely than G15 in yellow emission (within 570-640 nm). We also searched for activators that give distinct emission colors (e.g., red emission within 663-738 nm). Once the complementary AT sequences on the C55 probe and the activators hybridized, AgNC emission developed^5,6^. In the traditional test-tube experiments, the NC probe-activator mixtures were heated to 90-95 °C for a minute and gradually cooled down to room temperature to allow for hybridzation^4,5^. On the *MiSeq* chip, a constant temperature of 40 °C was maintained to enable hybridization (**Supplementary Fig. S9; *Methods***). This relatively low hybridization temperature extended the life of the chip, allowing us to go through at least 20 rounds of activation experiments on a single chip (hybridized with C55 probes, washed and imaged, and then removed C55 probes using alkaline solution) without showing any degradation, thus providing highly reproducible selection results (Spearman’s ρ= 0.93 ± 0.005 for the red NCBs and 0.88 ± 0.024 for the yellow NCBs, **Supplementary Fig. S10-S11**).

After injecting the common C55 NC probes into the *MiSeq* chip, the entire chip was scanned using a wide-field fluorescence microscope equipped with a metal halide illuminator, an sCMOS camera and an xyz translation stage. The activated NCBs were sequentially imaged using the filter cubes designed for conventional red emitters (e.g., Cy5, Ex/Em: 620/60, 700/75 nm) and yellow emitters (e.g., TRITC, Ex/Em: 535/50, 605/70 nm). These two filter cubes were selected due to their popularity in fluorescence imaging. A custom bioinformatics and imaging processing pipeline^28^ was employed to identify activator sequence behind each activated NCB spot (**Fig. 1c**). After ranking the activators based on their median activation brightness (each activator had 457 ± 308 polonies on a *MiSeq* chip), we could clearly distinguish strong activators from weak ones in the two emission channels (**Fig. 1c** right).

Compared to other high-throughput screening methods that relied on fluorescence from single molecule for characaterization^30^, photobleaching was not a severe issue in our approach, as each activator polony contained tens to hundreds of activated NCBs. Besides, by employing an auto-scan algorithm and shutter control (***Methods***), excitation dose to each polony was precisely regulated, avoiding any uneven photobleaching and ensuring consistent imaging conditions. By acquiring a fluorescence image every 200 ms, intensity time traces of polonies were obtained, which could be fitted with a single-exponential decay. After one second of strong illumination (∼10 W/cm^2^), polony intensity decreased by ∼20% at most (**Supplementary Fig. S13**). Based on the chip screening results, twenty top-ranked and twenty bottom-ranked activators were selected for further investigation in test tubes by traditional fluorometry. Using G12 activator (ATCCGGGGTGGGGTGGGG) as the standard for red NCB comparison, the *MiSeq* chip screening results were 85% accurate in both true positive and true negative selections (brighter or darker than G12 in chip screening and confirmed in fluorometry, **Fig. 1d, Supplementary Fig. S22-S23**, and **Supplementary Table S2**). When comparing the chip screening results with the test-tube results, a Pearson correlation coefficient of 0.50 was obtained (**Fig. 1d**). We noticed the intensity differences found on *MiSeq* chips were substantially smaller than those found in test tubes (e.g., 1.20-fold red-emission difference was found between rAct1 and G12 NCBs on a *MiSeq* chip, but it became 2.94-fold in test tubes). The underestimation was attributed to the relatively higher fluorescence background on *MiSeq* chips and the variations in polony numbers among the library sequences (for instance, ∼240 activators had less than 30 polonies). In a separate experiment, we selected ten activators there were found brighter than G15 (the standard for yellow NCB comparison) in *MiSeq* chips. All ten activators were still brighter than G15 in test tubes. In particular, we found yAct4 (TTGGTGGGTGGGGTGGGG) 2.03-fold brighter than G15 in activating C55 in the yellow channel (enhancement ratios were 1,125 vs. 553, **Supplementary Fig. S24** and **Supplementary Table S3**) and rAct1 (TCCATTGGTGGGGTGGGG) 2.94-fold brighter than G12 in the red channel (enhancement ratios were 1,292 vs. 439, **Fig. 1e, Supplementary Fig. S22** and **Supplementary Table S2**). We emphasize that the small-scale investigations carried out in test tubes, such as single-nucleotide substitutions from G15 at each nucleobase position (totally 3×18 = 54 variants), would not lead to any activators that are significantly brighter than G15 (**Supplementary Fig. S18**).

### Identification of critical nucleobases in stabilizing bright AgNC chromophores

In our first library design (library_1, **Supplementary Table S1**), the 18-nt-long canonical G15 activator was divided into three 6-nt-long segments, and each segment was separately randomized to create 3×4^6^-2 = 12,286 activator mutations (two were G15 duplicates in 3×4^6^ combinations, **Supplementary Table S1**). The chip screening results on library_1 clearly indicated that segment_2 prefers to be conserved (GGTGGG) in order to maintain NCB brightness (**Fig. 2**). In contrast, randomizing segment_1 still produced many bright red NCBs, especially when segment_1 became C-rich. The effect of segment_3 was diverse, suggesting an indirect activation role. The 6-segment interrogation further revealed that segment_22 (positions 10-12) is more important than segment_21 (positions 7-9) in creating bright red NCBs (**Fig. 2c** and **Supplementary Fig. S14**). Independent investigations using library_2 and library_3 (∼28,000 frame-shifted activators, **Supplementary Table S1**) also confirmed that the nucleobases in positions 10-12 are critical in C55 activation (**Supplementary Fig. S14**). These results indicated that bright AgNC chromophores are most likely “clamped” by the two strands at positions 7-12 (**Fig. 2b**), possibly forming silver-mediated pairs between the two strands^31,32^.

Drawing from the **Fig. 2** results, one possible design rule for creating bright red NCBs on C55 could be having an activator with a C-rich segment_1, a GC-rich segment_21, a G-rich segment_22, and a TC-rich segment_3. Nevertheless, the activator CCCCCCGCGGGGTTTCCC (termed G5) actually had a low red enhancement ratio (39, as compared to 439 for G12; **Supplementary Fig. S29** and **Supplementary Table S8**). This result clearly indicated that segments do not work alone – cooperativities among the segments determine the activation color and intensity of an NCB. Whereas previous investigations showed that more guanines in the activator generally leads to brighter red emission^4^, our large-scale investigations revealed a different design rule – brighter red NCBs can be achieved with fewer numbers of guanines (e.g., 10G_5 in **Supplementary Fig. S25** and **Supplementary Table S4**). As the results from our high-throughput screening could not be easily transformed into simple design rules, we trained machine learning algorithms on the big data and used them to rationally design bright NCBs (see ***Discussion***).

### Discovery of polar opposite twins using NGS chips

Taking advantage of the chip screening platform, we searched for POT-NCBs that have the most extreme color or intensity differences (**Fig. 3a-c**). Although NCBs were previously used for single-nucleotide polymorphism detection^5,14^, only tens of activators were tested, providing little information on the rules to design POTs. In contrast, library_1 alone contained more than 110,000 pairs of twin NCBs, where the top 2,000 pairs were readily candidates for POTs. Upon examining these 2,000 pairs of twin NCBs (**Fig. 3d**), it was clear that the nucleobases in positions 4-6 are critical for creating yellow POTs (e.g., bright (x-axis)→dark (y-axis) conversion by G/T→C/A substitution at position 5 and G→C substitution at positions 4 and 6), while the positions 2-4 are critical for creating red POTs (bright→dark conversion by C→ATG substitution). Fifteen top POT pairs were further investigated in test tubes. The most extreme yellow and red POTs had 31-fold (yPOT5-yPOT6 with G→C substitution at position 5) and 9-fold (rPOT5-rPOT6 with C→T substitution at position 4) differences in their enhancement ratios, respectively (**Fig. 3e**). For the ease of comparison, we termed the difference in the enhancement ratios the “POT difference ratio” (**Supplementary Fig. S30-S31** and **Supplementary Table S10-S11**), where the pairs with the largest POT difference ratios are the most extreme POTs.

**Fig. 3.**
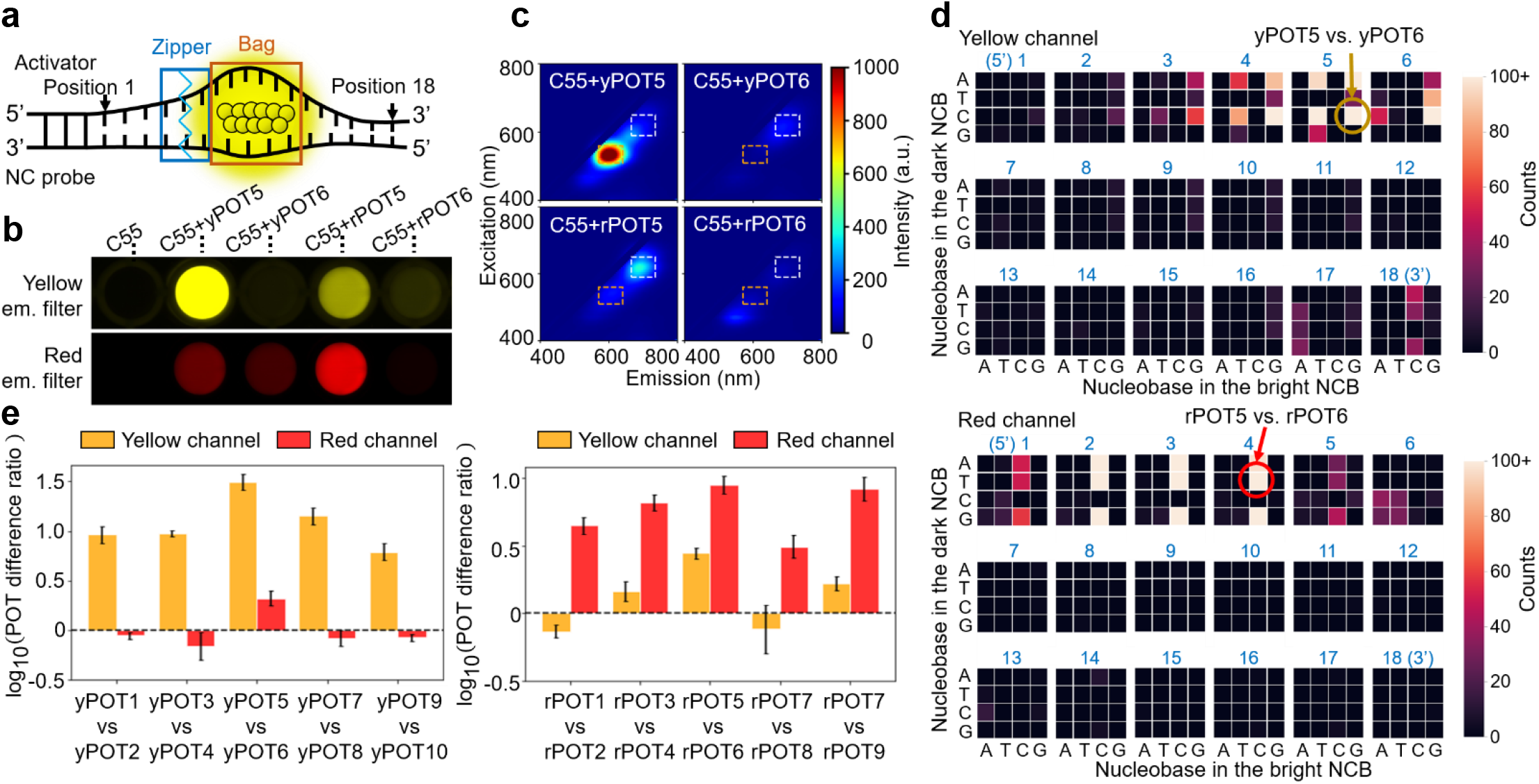
*S*ubstitution hotspots to generate polar opposite twins (POTs) revealed by *MiSeq* chip selection. **a** Schematic of the zipper bag model. The blue box represents the “zipper” location (e.g., positions 4-6 for yellow POTs) and the orange box represents the “bag” location (i.e., the critical zone at positions 7-12). When the zipper does not seal well, the bag is leaky, thus leading to a low chemical yield and dimmer NCB. **b** Plate-reader images acquired using yellow (top) and red (bottom) excitation/emission filter sets and the sequences of the representative POTs. Large differences in fluorescence enhancement ratios were seen in these twin NCBs (C55+yPOT5 vs. C55+yPOT6 for yellow channel, and C55+rPOT5 vs. C55+rPOT6 for red channel), making them POTs. **c** 2D spectra of the representative POTs in the yellow (orange dashed box) and red (white dashed box) emission channels. **d** Heat maps of the top 2,000 twin NCB pairs in library_1. Here the x-axis and the y-axis represent bright to dark conversion in these twin NCBs. These heat maps clearly indicated that the nucleobases in positions 4-6 are critical for creating yellow POTs, while the positions 2-4 are critical for creating red POTs. **e** The POT difference ratios of five representative yellow (left) and red (right) pairs of POTs in the two emission channels. Through fluorometry characterization, the yPOT5-yPOT6 pair and the rPOT5-rPOT6 pair were identified as the most extreme yellow and red POTs, respectively, reaching POT difference ratios as high as 31 and 9. Definition of the POT difference ratio can be found in the methods. Error bars: mean ± s.d. in logarithmic scale, with 3 repeats for each pair of POTs.

The POT difference ratio reflected the sample brightness difference at the ensemble level, which is equal to the product of “chromophore chemical yield ratio” and “single-emitter brightness ratio”. Using fluorescence correlation spectroscopy (FCS)^33,34^, we found the chromophore chemical yield of rPOT5 NCB 5.54-fold higher than that of rPOT6 NCB (20% vs. 3.6%), and the single-emitter brightness of rPOT5 1.64-fold higher than that of rPOT6 (5.67 kHz vs. 3.45 kHz, **Supplementary Fig. S16**). The product of the 5.54× chromophore chemical yield ratio and the 1.64× single-emitter brightness ratio was indeed the 9× POT difference ratio measured by fluorometry. Similarly, the chromophore chemical yield of yPOT5 NCB was 16.33-fold higher than that of yPOT6 NCB (25.8% vs. 1.6%) and the single-emitter brightness of yPOT5 NCB was 2.17-fold higher than that of yPOT6 NCB (7.04 kHz vs. 3.24 kHz, **Supplementary Fig. S17**). The product of the 16.33× chromophore chemical yield ratio and the 2.17× single-emitter brightness ratio (35×) was close to the 31× POT difference ratio measured at the ensemble level. Our investigation of POTs led to two important findings. First, the red AgNCs in rPOT5 and rPOT6 were actually different species as their excitation peak wavelengths (610 vs. 605 nm, **Supplementary Fig. S30a**), absorption spectra (a clear peak at 610 nm in rPOT5 NCBs spectrum but no clear peak in rPOT6 NCBs spectrum, **Supplementary Fig. S19d**), and single-emitter brightness were all different. Second, the chemical yield of AgNC chromophores could be significantly altered by substituting single nucleobases at positions (2-6 in **Fig. 3d**) outside the critical zone (7-12 in **Fig. 2**). Based on these findings, we proposed a zipper bag model that explains the mechanism behind POT formation (**Fig. 3a**).

## Discussion

### Investigation of the zipper bag model

In our zipper bag model, the bag is the critical zone (positions 7-12) that holds the AgNC chromophore while the zipper is the POT hotspot that seals the bag. A subtle change in the zipper can alter the sealing condition of the zipper bag, which perturbs the short-range ligand environment around the AgNC inside the bag and possibly changes its binding footprint with the bag (**Fig. 3a**). We have previously shown that by slightly shifting the position of the activator with respect to the NC probe, a new ligand environment can be created around the AgNC that alters its emission spectrum^14,17^, and we believe such a nucleobase-AgNC interaction is within a short range (≤ 1 nm)^14^. When studying AgNC structures using 193 nm activated-electron photodetachment mass spectrometry (a-EPD MS), we have previously found two Ag^10^ clusters can be completely distinct chromophores due to very different binding footprints in their DNA hosts^35^. The earlier structural studies by extended X-ray absorption fine structure (EXAFS) spectra complemented the a-EPD MS footprint results^11,36^. Here by using FCS, we further demonstrated that a change in zipper may result in not only a distinct AgNC chromophore in the bag but also a different chemical yield of the chromophore, thus providing a basis for POT formation.

In our model, the zipper may not necessarily be formed by the Watson-Crick (WC) basepairs – it can also be formed by the silver-mediated pairs (e.g., the C-Ag^+^-C pair)^31,32^. Upon close examination, we believe the yellow POT zippers at positions 4-6 are caused by a WC pair GC or a wobble pair GT (where G is on the activator and C or T is on the NC probe), as disrupting the GC or GT pair at these positions often creates dim yellow NCB samples (**Supplementary Fig. S31b**). As aforementioned, these dim yellow samples attribute to less emissive AgNC chromophores and lower chemical yield of the chromophores. In contrast, the red POT zippers at positions 2-4 may be formed by a silver-mediated pair C-Ag^+^-C. Disrupting the C-Ag^+^-C pair at these positions darkens red NCB samples (**Supplementary Fig. S30b**). One recent report showed evidence that silver-mediated heteroduplexes (e.g., C_11_-Ag^+^_N_-T_11_) could be less stable than their homoduplex counterparts (e.g., C_11_-Ag^+^_N_-C_11_)^31^, while another report showed a Ag^+^-mediated interaction at a place away from the AgNC core^37^, both supporting the hypothesis behind our model.

Why are the zipper locations different for the yPOT5-yPOT6 pair (at position 5) and the rPOT5-rPOT6 pair (at position 4)? One possibility is red AgNC chromophores have higher silver stoichiometries and larger footprints in the bag, pushing the zipper locations further away from the bag (positions 7-12). Using electrospray ionization mass spectrometry (ESI-MS), Gwinn’s group has previously shown a general trend for yellow chromophores having a smaller core (Ag_10_-Ag_11_) while red chromophores having a larger core (Ag_14_-Ag_16_)^10^. According to their rod-shaped model^9^, red chromophores are expected to have larger footprints in their DNA hosts. However, in Gwinn*’*s experiments, their AgNC chromophores most likely stabilized inside dimers of 10-mers, which create ligand environments that are different from our activator/NC probe systems. To investigate the silver stoichiometries of the two major chromophores in our POT experiments, we purified yPOT5, yPOT6, rPOT5 and rPOT6 NCB samples using 20% native polyacrylamide gel electrophoresis (native PAGE) (**Supplementary Fig. S20; *Methods***) and analyzed the purified samples by ESI-MS (**Supplementary Fig. S21**). Upon close examination of the gel shift results, yPOT5 NCB clearly had a higher mobility as compared to rPOT5 NCB, possibly due to a more condensed form of the chromophore-DNA complex. Similar results were also observed for the yellow standard G15 NCB, which ran faster than the red standard G12 NCB in gel. We also noticed that the brighter NCBs (yPOT5 and rPOT5) had slightly higher mobility than their dim counterparts (yPOT6 and rPOT6), which could infer that the activators of the dim chromophores are “wrapped” loosely around the C55 NC probe and hence slightly larger complexes were formed.

Since our NCB constructs were much larger (45-nt long for the NC probe strand and 48-nt long for the activator strand) than the constructs used in previous studies involving ESI-MS analysis^10,38,39^ (10-to 26-nt-long), extensive cationic metal adduction during the ESI process prevented us from deciphering the exact silver stoichiometries in purified NCB samples. To circumvent this issue, an ion-pairing reagent, octylamine, was added to the ESI samples to suppress salt adduct formation^40-43^. Although the number of adducts was reduced, the addition of octylamine destabilized the duplex NCB structures. ESI-MS of purified yPOT5 NCB and rPOT5 NCB samples containing octylamine showed clear peaks of single-stranded species (i.e., C55 NC probe and activator) with various silver stoichiometries (**Supplementary Fig. S21**). Whereas it was still difficult to conclude the exact silver stoichiometries in the original duplexes, introducing octylamine to the MS spray solution led to well-resolved single-stranded species with highly reproducible numbers of silver atoms on them. The C55 NC probes from yPOT5 NCB and rPOT5 NCB carried 0-4 and 0-3 silver atoms, while the activators yPOT5 and rPOT5 carried 0-7 and 0-6 silver atoms, respectively. These results indicated that the original silver stoichiometry for the intact yPOT5 NCB may be larger than that of the intact rPOT5 NCB. Although our results are contradictory to previous investigations that suggest red AgNC chromophores have a larger core^10^, we predict that the red chromophores featured in this study have a larger footprint in the activator/C55 bag, owing to different AgNC shapes and DNA conformations^9,12,13,35,44^. In future studies, we aim to pinpoint the binding sites of AgNC chromophores within the activator/NC probe duplexes by employing activated-electron photodetachment mass spectrometry (a-EPD MS), a structural characterization technique and tandem MS^n^ method that was previously shown to reveal the binding sites of AgNCs in shorter single-stranded DNA hosts (up to 28-nt long)^35^. Alternatively, X-ray crystallography of DNA-templated AgNCs can reveal not only the binding sites but also the binding geometries of surrounding bases to AgNCs^12,13,37,44^, provided that NCBs can be crystallized.

### Predictive design of bright NCBs using chip screening results and machine learning models

Since the results from our high-throughput screening (**Fig. 2**) cannot be easily transformed into simple design rules, we take advantage of machine learning algorithms to classify NCBs and uncover sequence features that give bright NCBs. Machine learning approaches have previously helped identify sequence features in DNA hosts that preferentially stabilize bright AgNCs, establishing the first statistical model for rational design of AgNCs with desired colors^15,45^. However, such a model was built upon the emission properties of ∼2,000 AgNCs sandwiched between two identical strands (8-mers to 16-mers)^46^. Our AgNCs were different, as each of them was stabilized within an 18-nt long activator and a 15-nt long C55. Although both robotic-well-plate studies^46^ (2,000 short strands) and NGS screening (40,000 NCBs in this report) covered only a small fraction of the overall ligand composition space of 10-mers and 18-mers, respectively, statistical models could be built based on these small fractions of data. This was due to the fact that a majority of the ligand composition space was only occupied by dark AgNCs. Previous selection and our screening here focused heavily on the sequences that had a high chance to produce bright AgNCs, thus giving a higher “effective” fraction of search space.

To connect activator sequences to NCB brightness, we adopted machine learning algorithms to recognize sequence features (which consist of motifs and motif locations) in the bright activators. Following the approaches proposed by Copp and Gwinn^15,45,46^, we labeled the top 30% NCBs (3,600) as “bright” class and the bottom 30% as “dark” class. 339 and 567 features from the bright and dark classes in the yellow channel (denoted as bright yellow and dark yellow features) and 402 and 1,164 features from the bright and dark classes in the red channel (denoted as bright red and dark red features) were separately identified by MERCI^47^. To decrease the chance of overfitting, we further narrowed down to a set of the most discriminative features with 61 bright yellow, 121 dark yellow, 103 bright red and 112 dark red features using Weka^48^ (**Supplementary Table S13-S14**; ***Methods***). A feature vector was employed to describe the location of each activator in the high-dimensional space for classification model development (**Fig. 4a**).

**Fig. 4.**
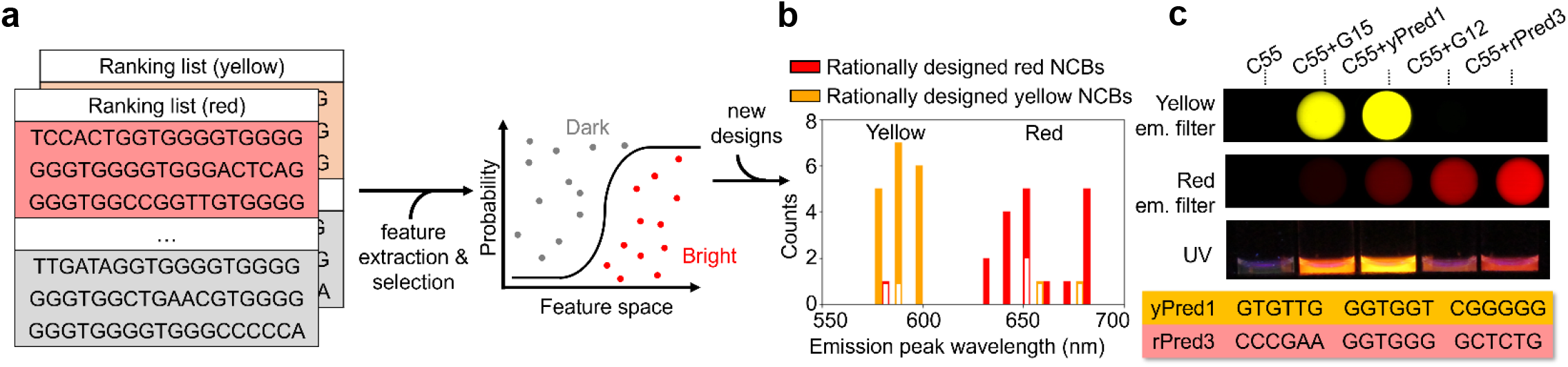
Predictive design of bright yellow and red NCBs based on machine learning results. **a** From the chip selection results, we labeled the top 30% NCBs as ‘bright’ class and the bottom 30% as ‘dark’ class. The sequence features of these selected NCBs were then extracted by MERCI and selected by Weka. The resulting feature vectors thus defined the location of individual activators in the high-dimensional space. Several machine learning models were tested for activator classification, among which the logistic regression had the best performance. **b** Forty new activators were rationally designed and tested by fluorometry. Three out of the 20 rationally designed red NCBs actually showed either low emission or yellow emission (85% test-tube validation accuracy), while three of the 20 rationally designed yellow NCBs showed low emission of red emission (also 85% test-tube validation accuracy). Empty boxes represent the failed designs with low emission. **c** Plate-reader images acquired using yellow (top) and red (bottom) excitation/emission filter sets, and the activator sequences of the two successfully predicted NCBs and one failed prediction.

A number of models were established for classifying the chip screening results, based on algorithms such as logistic regression (LR), linear discriminant analysis (LDA), decision tree (DT), AdaBoost (ADA), and support vector machines (SVM) (**Supplementary Table S12**). To evaluate the model performance, we defined the accuracy of the model (Acc) as (T_B_ + T_D_)/(T_B_ + F_B_ + T_D_ + F_D_), where T_B_ is the number of true predictions that the model makes for bright activators, T_D_ and F_D_ are the numbers of true and false dark predictions, and F_B_ is the number of false bright predictions. In other words, Acc represented the fraction of test sequences that the model correctly identifies as “bright” or “dark” activators. We found the model built on LR has the best performance, achieving an average accuracy of 0.89 and 0.87 in the yellow and red emission classification, respectively. In our model development process, the categorized dataset was divided into a training set (80% of the selected sequences) and a test set (20% of the selected sequences). The selection process iterated 5 times while rotating the training set and the test set, resulting in a 5-fold cross-validation (CV) that guarantees the model consistency (**Supplementary Fig. S34**). As expected, the propensity of an activator to be “bright” is not only determined by having “bright” motifs within the activator but also by positioning these motifs at proper locations.

Separately, based on the most discriminative features identified by Weka^48^, 1,000 bright yellow and 1,000 bright red activator candidates were rationally designed *in silico* (**Supplementary Fig. S35; *Methods***). In the high-dimensional space, we employed the minimal “edit distance”^46,49^ to identify the “closest” activator sequence in the library dataset for each of the rationally designed candidates. When the closest library sequence was not among the top 200 bright activators screened, the candidate was discarded. Besides, when the candidate sequence had less than 3 or more than 5 single-base mutations, they were also discarded. After going through these candidate refining steps, we were down to 100 bright yellow and 54 bright red candidates. Among these candidates, the LR model classified 85 and 41 of them as bright yellow and red activators, respectively. From these most promising candidates, 20 were randomly selected, synthesized, hybridized with the C55 probes in test tubes, and measured by a fluorometers.

Compared to the random activator sequences (**Supplementary Fig. S28** and **Supplementary Table S8**), our design and classification pipeline generated new activators that were 8.5 times (85% vs. 10%) and twice (85% vs. 40%) more likely to be bright yellow and red activators, respectively (**Supplementary Fig. S26-S27** and **Supplementary Table S6-S7**). Besides, the average enhancement ratio of rationally designed yellow and red activators were 22 times and twice higher than that of random sequences, respectively (**Supplementary Table S8**). Moreover, while all random sequences gave more or less red emission, our pipeline successfully produced NCBs with yellow emission (**Fig. 4b**). In particular, among all bright candidates tested, we identified a new yellow activator (yPred1: GTGTTGGGTGGTCGGGGG, with only 12 guanines) and a new red activator (rPred3: ATCCCTCGGGGAGGGGGC, with only 9 guanines) that were 2.13-fold and 1.30-fold brighter than the gold standards G15 and G12 in activating the C55 NC probe, respectively (**Fig. 4c, Supplementary Table S6-S7**).

### Summary

We have performed high-throughput screening on more than 40,000 activators using repurposed NGS chips. Not only did we discover new NCBs (yAct4 and rAct1) that are 2-3 times brighter than the known best (G15 and G12, **Fig. 1**), but we also identified a critical zone in the activator (positions 7-12) that stabilizes bright AgNC chromophores (**Fig. 2**). In the search for the most extreme POT-NCBs, the chip platform helped identify a red pair (rPOT5-rPOT6) and a yellow pair (yPOT5-yPOT6) with POT difference ratios as high as 9 and 31, respectively (**Fig. 3**). By probing the NCBs at the near single-molecule level, we confirmed the observed brightness difference at the ensemble level is attributed to the differences in the single-emitter brightness and the chromophore chemical yield (**Supplementary Fig. S16-S17**). Based on the findings of the critical zone (positions 7-12) and the POT hotspots (positions 2-4 for the red POTs and positions 4-6 for the yellow POTs), we proposed a zipper bag model that explains how POT hotspots lead to the creation of distinct AgNC chromophores and contribute to the difference in chromophore chemical yields (**Fig. 3**). As the results from high-throughput screening could not be easily converted into simple rules for designing bright NCBs, we employed machine learning algorithms to classify the screening results and used the trained model to rationally design multicolor NCBs. Forty new NCBs were generated, clearly showing two designated color bands (**Fig. 4**). We also found brighter NCBs could be achieved with fewer numbers of guanine bases in the activators. Our chip screening platform can facilitate the development of new chemical sensors based on DNA-templated AgNCs^50^ or be used to study other metal nanoclusters templated in DNA^51-53^. To our knowledge, this article is the first report that NGS chips are repurposed for high-throughput screening of fluorescent nanomaterials. Our high-throughput screening and machine-learning-based design pipeline is not only accelerating the discovery of new NCBs for diverse applications, but also providing insights into the chemical yield and the emitter brightness controlled by the sequence features. We anticipate new NCB-based sensors and new fluorescence barcodes will soon be developed based on the design strategy that we layout in this article.

## Methods

### NC probe preparation

Sodium phosphate dibasic anhydrous (Na_2_HPO_4_; F.W. 141.96), sodium phosphate monobasic monohydrate Na_2_HPO_4_·H_2_O; F.W. 137.99) and sodium borohydride (NaBH_4_) were purchased from Fisher Scientific, whereas silver nitrate (AgNO_3_) was acquired from Sigma-Aldrich. All oligonucleotides were purchased from Integrated DNA Technologies (IDT) and were purified by desalting. Deionized (DI) water (18 MΩ·cm) was used for all solution preparations.

In a typical preparation, a 15 µM (final concentration) NC probe solution was prepared by adding 12.5 µl of 1.2 mM NC probe (C55, **Supplementary Table S1**) to 940 µl of 20 mM sodium phosphate buffer (pH 6.6). The solution was vortexed for 2 s, and 45 µl of 4 mM silver nitrate solution was added to it. Again, the mixture was vortexed for 2 s. The solution was allowed to sit in the dark for 10 min at room temperature. For silver cluster formation, 7 µl of freshly prepared 13.2 mM NaBH_4_ solution was added to the reaction, resulting in a pale-yellow mixture, which was then stored in the dark overnight. The resulting NC probe solution had the [NC probe]: [Ag^+^]: [NaBH_4_] molar ratio of 1:12:6.

Before *MiSeq* chip experiment, a 0.5 ml centrifugal filter (cat. no. UFC503024, MilliporeSigma) was employed to remove excess silver ions. Purification protocol followed the manufacturer guidelines. The filtered solution was then diluted to 500 nM (DNA concentration, which was verified using the NanoDrop 2000 UV-Vis spectrophotometer, Thermo Scientific) before being injected into *MiSeq* chip.

### NCB preparation and in-solution validation

To activate NCB, 1.5 µl of 1.2 mM activator solution was added to a 120 µl aliquot of the previously prepared 15 µM C55 probe solution. The mixture was vortexed, centrifuged, and immersed in a hot water bath (90-95 °C) for 1 min, followed by gradually cooling down to room temperature for 1 hr. The activated NCB had the [NC probe]: [activator] molar ratio of 1:1. The fluorescence measurements started exactly at 1 hr after the addition of activator.

We quantified NCB fluorescence using a fluorometer (FluoroMax-4, Horiba) and a 100 µl quartz cuvette (16.100F-Q-10/Z15, Starna Cells). Both the excitation and emission wavelength scan ranges were set to be from 400 nm to 800 nm using 5 nm slit size, 5 nm increment step, 0.1 s integration time. Two control samples, a 20 mM sodium phosphate pH 6.6 buffer only sample and an NC probe only sample (with AgNCs but no activators) were also measured. The acquired spectra were saved as csv files and processed using a Python script.

### NCB fluorescence enhancement ratio calculation

We followed a similar definition described in ref. 5 to calculate the ensemble enhancement ratio of NCB after activation. However, in ref. 5, 1D spectra based on 580 nm excitation were acquired and area integrated intensities were calculated over 595 to 740 nm emission range. Here we collected 2D spectra of samples and calculated volumetric integrated intensities over the red (Ex/Em: 620/60, 700/75 nm) and the yellow (Ex/Em: 535/50, 605/70 nm) excitation/emission “windows” (**Supplementary Fig. S2**). From there we calculated the enhancement ratio:

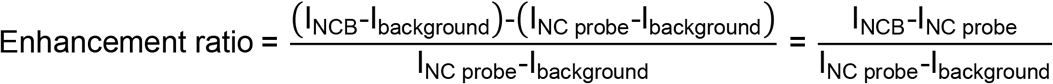

where I_NCB_ stands for the volumetric integrated intensity of NCB in red or yellow window, I_NC probe_ represents the volumetric integrated intensity of dark AgNC on the C55 probe, and I_background_ is the volumetric integrated intensity of the sodium phosphate buffer. The improvement ratio is simply the ratio of the enhancement ratio of an activator to that of the standard activator (G12 or G15). Similarly, the POT difference ratio is simply the ratio of the enhancement ratios of the twins:

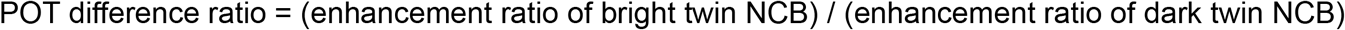

### NCB fluorescence visualization

Color photos of inactivated (NC probe only) and activated (the duplex) NCBs were acquired using a digital camera (PowerShot SX 500 IS, Canon) on a Syngene gel imager (with 365 nm excitation) (**Fig. 1b**). NCB fluorescence were also visualized using a gel imaging scanner (Typhoon 9500, GE Healthcare Life Sciences). For the Typhoon 9500 experiments, 240 µl of NCB sample was placed in a single well on the 96 multi-well plate. The fluorescence was acquired using the built-in Cy3 channel (EX: 532 nm, EM: 575 nm long pass) or the Cy5 channel (EX: 635 nm, EM: 665 nm long pass) while the PMT gain was set to 400 and the pixel size was 10 µm. The imaging results were saved as tiff files and changed to 16-bit false colors (yellow for Cy3-channel imaging and red for Cy5-channel imaging) using ImageJ.

### Activator library design

All activator strands contained a universal, 30-nt-long TA-rich hybridization segment followed by an 18-nt-long variable region (the activator) and an 8-nt-long restriction site (**Supplementary Table S1**). Additional adapters at 5’-end (P5 and SP1) and 3’-end (P7 and SP2) were designed by Illumina for sequencing purpose. To identify our library sequences, a 6-nt-long barcode was also needed and that was added to the 3’-end. The barcode for the canonical activator G15 was different from that of any other activators in order to monitor the sequencing yield. Three different libraries were established based on shifted frames, giving totally 40,068 unique activator sequences (**Supplementary Table S1**).

### NGS library preparation

A standard PCR process was performed using Q5 high-fidelity DNA polymerase from NEB (cat. no. M0491S). All PCR primers were purchased from IDT (**Supplementary Table S1**). The PCR procedure and the thermal cycler (Eppendorf, Mastercycler^®^ nexus) settings followed the protocol provided by NEB. The library sequences and the canonical activator were amplified separately, reaching a final concentration greater than 5 ng/µl for each tube. After PCR amplification, the concentration of DNA library was verified using a NanoDrop 2000 UV-Vis spectrophotometer (Thermo Scientific). Together with the fiducial markers (PhiX), the library sequences were immobilized and bridge amplified on an Illumina *MiSeq* chip, followed by sequencing using a 2×300 paired end reagent kit (v3, Illumina). For the 3-segment interrogation (**Fig. 2**), we targeted to have 1.2 million reads for the mutations in each segment. 10,000 additional reads were expected for the canonical activator G15. The fiducial markers, PhiX, were counted for 15%∼20% of the overall coverage. The actual numbers of reads varied from batch to batch, but were within 5-13 million reads.

### *MiSeq* chip preparation

After sequencing, *MiSeq* chips were kept at 4 oC in storage buffer (1X TBE buffer, cat. no. AM9865, Invitrogen). Before hybridizing with NC probes, all DNA strands covalently affixed to the *MiSeq* chip surface were denatured with 20 µl 0.1N NaOH solution for 5 min and then rinsed with 20 µl 1X TBE buffer 3 times to remove excess NaOH. This rinsing step removed untethered DNA strands containing residual fluorescent dyes from sequencing. Before the NCB screening experiment, chip was rinsed with working buffer (150 µl, 200 mM sodium phosphate buffer pH 6.6) three times. To cleave the unwanted sequence beyond the activator sequence, a 32-nt-long strand complementary to the restriction site (RE strand in **Supplementary Table S1**) was introduced to the chip and the chip was annealed at 40 °C for 40 min. After annealing, 1X TBE buffer was used to rinse the chip, followed by MauBI (cat. no. ER2081, Thermo Fisher Scientific) restriction enzyme digestion. The reaction buffer was prepared following the manufacturer’s protocol and 20 µl enzyme solution was kept in the *MiSeq* chip at 40 °C for 40 min. After digestion, the chip was washed with 20 μl 0.1N NaOH solution for 5 min and rinsed with 20 µl 1X TBE buffer for three times. The PhiX sequences were labeled using an Atto488-tagged probe (500 nM and 20 µl, **Supplementary Table S1**) for fiducial marker imaging. To optimize the annealing conditions, we evaluated the intensities of NCBs that had gone through different temperature treatments (40 °C for 40 min, room temperature for 40 min, and 90 °C for 10 min, **Supplementary Fig. S9**). Holding the chip at 40 °C for 40 min not only gave an excellent annealing result but also extended the chip life to up to 20 runs of NCB activation experiments. After testing different concentrations of C55 probes for the NCB screening experiment (**Supplementary Fig. S12**), we chose to use 20 µl of 500 nM C55 probe solution for all our chip experiments. The chip was imaged at room temperature with microscope settings stated below. After each experiment, the chip was washed with 20 µl 0.1N NaOH solution for 5 minutes, followed by rinsing with 20 µl 1X TBE buffer for three times and storing at 4 °C.

### Fluorescence microscopy and image acquisition

An open-source software, Micro-Manager^54^, was used to control an sCMOS camera (ORCA-Flash 4.0, Hamamatsu), an xyz translation stage (ProScan III, Prior Scientific), and an auto-shutter (Lambda SC, Shutter Instrument) on an Olympus IX71 fluorescence microscope for all our *MiSeq* screening experiments. A metal-halite illuminator (Lumen 200, Prior) and a 60× water-immersion objective (UPLSAPO60XW, Olympus) were used in the IX71 system. We developed a MATLAB script to generate the position list of each field of view for automatic acquisition. On each *MiSeq* chip, we acquired 60 images (FOV: 220 × 220 µm^2^) per row for 3 rows on both floor and ceiling (totally 360 images), covering a total surface area of 5.81 mm^2^. It is worthwhile to note that polonies in some regions of the chip were not registered in the Illumina sequencing files. To bypass most of these unregistered regions, we shifted the imaging starting position by 380 µm in the vertical direction and 1,611 µm in the horizontal direction with respect to the reference point at the bottom left corner (**Supplementary Fig. S3c**). We first recorded fiducial marker images (Atto488, FOV: 220 × 220 µm^2^, 1 second exposure time, green channel, Ex/Em: 480/40, 535/50 nm, cat. no. 51006, Chroma), and then recorded NCB images in both red (Ex/Em: 620/60 nm, 700/75 nm, cat. no. 49006, Chroma) and yellow channels (Ex/Em: 535/50 nm, 605/70 nm, cat. no. 49004, Chroma) under the same imaging settings.

### Flat-field correction

We implemented flat-field correction to eliminate the variation of fluorescence background across the field of view (FOV, **Supplementary Fig. S3a-b**). A Gaussian-blur filter was applied to generate the flat-field reference image for each FOV. We found that a Gaussian-blur filter with sigma equal to 50 best fit our purpose. The corrected imaged used for the following analysis were computed as follows:

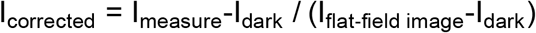

where I_measure_ is the recorded fluorescence images, I_flat-field image_ is the flat-field reference image generated by the Gaussian-blur filter, and I_dark_ is the dark image recorded using 1 second exposure time while illuminator is turned off.

### Fluorescence correlation spectroscopy (FCS) measurements

FCS measurements were carried out using a confocal system (Alba v5, ISS) built around a Nikon microscope body (Eclipse TE2000-U, Nikon). A super-continuum laser (SuperK EVO EU-4, NTK Photonics) and a 60× water-immersion objective (CFI Plan Apochromat VC, Nikon) were used in the FCS experiments. To validate the NCB chemical yield measurement results, an Alexa647N-labeled ssDNA probe was used to generate the concentration calibration curve shown in **Supplementary Fig. S16-S17**. All FCS measurements were carried out using 200 µl samples in 8-well chamber slides (Nunc Lab-Tek, Thermo Fisher Scientific). The laser beam was focused 25 µm into the sample for all FCS measurements in this work.

### FCS analysis

Autocorrelation curves were fitted using the software package provided by ISS, giving estimates on the average number of emitters in the detection volume (*N*) and the average translational diffusion time constant (τ). Single-emitter brightness (SEB) was computed based on:

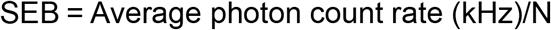

The “activated” NC probe concentration was derived from the calibration curve established by the Alexa647N probe (**Supplementary Fig. S16-S17**). NCB chemical yield was then computed as:

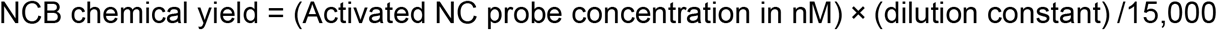

where 15,000 nM represents the DNA concentration in the original reaction (i.e., 15 µM).

### Absorption measurements

Absorption spectra of NCBs were measured using Cary 5000 UV-Vis-NIR spectrometer from Agilent. 500 µl of 15 µM NCB solution was prepared following aforementioned protocol and was injected into a 700 µl Micro Fluorescence Cuvette from Thorlabs. The dual-beam mode was used with baseline/zero correction. All absorption spectra were measured from 300 nm to 800 nm with slit size of 2 nm (**Supplementary Fig. S19**). The acquired data were processed and analyzed using Python scripts.

### Image alignment algorithm

A custom bioinformatics and imaging processing pipeline named CHAMP (<u>C</u>hip-<u>H</u>ybridized <u>A</u>ssociated <u>M</u>apping <u>P</u>latform) was developed by Finkelstein’s group and the detailed algorithm description can be found in ref. 28. CHAMP helped decipher the activator sequence behind each activated NCB spot (termed the NCB-CHAMP selection method, **Fig. 1c** and **Supplementary Fig. S4-S5**). In brief, mapping the alignment markers was done at four stages. First, a rough alignment was carried out using Fourier-based cross correlation, followed by a precision alignment using least-squares constellation mapping between FASTQ and *de novo* extracted NCB spots. We built up the consensus sequences and their corresponding information (e.g., lane number, tile number, and x-y coordinates) at all reported positions in the FASTQ file using the ***map*** command. Second, the ***init*** command was executed to record the metadata of imaging settings (e.g., rotation and scaling). Third, the ***h5*** command was applied to generate a single hdf5 file containing all 512×512 PhiX fiducial marker images. Fourth, the ***align*** command transformed the processed sequence information into pseudo-images and performed precise alignment. The output files were saved individually by image positions. The content included x, y coordinates of each sequence and the corresponding sequence ID. To analyze our NCB images, we developed an additional function named ***ncb***, which corrected the uneven illumination using flat-field correction. A bootstrap method was then performed to derive the median intensity of each activator in order to rank the NCB brightness (**Supplementary Fig. S36**).

### Feature extraction/selection and machine learning model establishment/validation

The feature extraction was performed using MERCI^47^. For all extraction processes, both positive and negative thresholds were set to 5%, the maximal motif length was set to 6 bases, and the maximal number of wildcard nucleotide (A, T, G, C, or nothing) was set to 1 base. For example, in 5-fold cross-validation, the threshold was set to be 144 (5% of 2,880 sequences for each class). Separately, to extract “bright” features, the entire bright and dark classes were used (3,600 sequences for each class) and 180 was set as the threshold. The dark feature extraction was performed by simply swapping the bright and dark classes with the same parameter settings. The extracted motifs were then processed with Python scripts to include the position information. 339 bright yellow, 567 dark yellow features, 402 bright red and 1,164 dark red features were separately identified.

To decrease the chance of overfitting, we further narrowed down to a set of the most discriminative features with 61 bright yellow, 121 dark yellow, 103 bright red and 112 dark red features using Weka^48^ – a process we termed feature selection. The attribute evaluator was set to “CfsSubsetEval”^55^ and the search method was set to “GreedyStepwise”^48^. CFS scored a feature subset based on high correlation of features with predictive classes and low inter-correlation of features^56^. The “greedy” algorithm started with an empty set and iteratively added the feature that maximizes the gain in the CFS score. Feature selection process stopped when any additional feature decreased the CFS score. All other parameters were set to default values (**Supplementary Fig. S34**).

After feature extraction and selection, classification models were established based on various ML algorithms including logistic regression (LR), linear discriminant analysis (LDA), decision tree (DT), AdaBoost (ADA), and support vector machines (SVM), using the scikit-learn package in Python. 5-fold cross-validation was performed to evaluate the model performance. The best model (i.e., LR model) was employed to rationally design bright and multicolor NCBs (**Supplementary Fig. S34**).

### *In-silico* design of bright NCBs

To rationally design red and yellow NCBs, we again divided the 18-nt-long activator into 3 segments. Based on the most discriminative features identified by Weka, we sampled the distribution of these features in each segment and generated a list of common motifs with their corresponding positions. Please note that the features selected from each segment could slightly go beyond the range of that segment. To construct a red NCB candidate, we assigned 3 features to the blank 18-nt template, starting with feature_1 insertion into segment_1. As feature_1 could go beyond segment_1, feature_2 might have an overlap with feature_1 when being inserted into segment_2. In that situation, the design algorithm would replace feature_2 with another feature to ensure no overlap. However, if any two features shared identical bases at their overlapping site, they were considered as “compatible” and could be inserted into the same template. For example, as shown in **Supplementary Fig. S35**, feature C_CTG (positions 1-5) and feature GGG_GC (positions 5-10) shared a guanine base at the overlapping site (position 5). Consequently, they were compatible and were used in constructing a bright NCB candidate. The same procedure was repeated until a compatible feature for segment_3 was found. Once all three features were inserted into the template, the remaining blank positions were filled up based on the composition popularity (at the same positions) from the bright class sequences. The edit distance^46,49^ of the new candidate was then assessed. We only selected new candidates with edit distance between 3 to 5 from the top 200 bright activators screened on chip for test-tube investigation (**Supplementary Table S6** and **S7**).

### NCB mobility evaluation in native PAGE gels

30 µM of dark C55 probes were prepared following the previously stated protocol. The same molar ratio of [NC probe]: [Ag^+^]: [NaBH_4_] = 1:12:6 was used while doubling the amount of each chemical. 30 µM NCB sample was prepared by adding 3 µl of 1.2 mM activator to the 120 µl, 30 µM C55 probe solution, followed by the aforementioned hybridization and buffer exchange protocol.

Hand-cast native polyacrylamide gels were prepared by mixing 3.75 ml 40% acrylamide solution (cat. no. HC2040, Invitrogen), 0.75 ml TBE 1X buffer (Invitrogen), 3 ml DI water, 75 µl 10% w/w ammonium persulfate solution (cat. no. HC2005, Invitrogen), and 7.5 µl TEMED (cat. no. 45-000-226, Fisher Scientific), targeting gel percentage at 20%. Gel solution was then poured onto the 1.0 mm empty gel cassette (cat. no. NC2010, Life Technologies). The cured gel was pre-run at 60V, 5 mA for 30 min in 1X TBE buffer before loading any samples. The DNA ladder was prepared by mixing 4 µl of 100 µM 20/100 ladder (cat. no. 51-05-15-02, Integrated DNA Technology) with 11 µl DI water, 4 µl 5X loading dye (cat. no. LC6678, Invitrogen), and 1 μl SYBR gold nucleic acid gel stain (cat. no. S11494, Invitrogen), while NCB samples were prepared by mixing the 16 µl, 30 µM NCB solution with 4 µl 5X loading dye. After the gel was pre-run for 30 min, each lane was loaded with 20 μl NCB samples or ladder and the gel was run at 50V, 5 mA for 800 min. All gels were imaged on Syngene gel imager (with 365 nm excitation) to evaluate the mobility. As no SYBR gold dye was added to the NCB samples, the gel bands of NCBs showed their nature fluorescence colors (**Supplementary Fig. S20**).

### NCB elution from native PAGE gels

NCB gel bands were extracted using a gel band cutter and collected in a 1.5 ml Eppendorf tube, followed by smashing the gel into pieces with a plastic stick. For elution, 450 µl of 20 mM SPB pH6.6 solution was added to each tube. The tube was shaken for 1 hr and stored at room temperature overnight. The suspension was then filtered using a micro-centrifugal filter (cat. no. F2517-5, Thermo Scientific) at 8,000 g for 20 min, followed by buffer exchange using the 0.5 ml centrifugal filter described above (cat. no. UFC503024, MilliporeSigma) and 10 mM ammonium acetate buffer (cat. no. AM9070G, Invitrogen).

### Sample preparation for native mass spectrometry

Two gel-purified NCB samples (yPOT5 and rPOT5) were desalted and buffer exchanged into 10 mM ammonium acetate using a spin column (Micro Bio-Spin™ P-6 Gel Columns, Bio-Rad). Octylamine was added to aliquots of NCB solution at a concentration of 0.1% (v/v), to reduce the extensive metal cationic adduction that is commonly seen for ESI-MS analysis of oligonucleotides > 20 nt^40-43^.

### Native mass spectrometry by direct infusion

3-5 µl of 5 µM purified yPOT5 and rPOT5 NCB solutions were loaded into Au/Pd-coated nanospray borosilicate static tips (prepared in-house) for nano electrospray ionization (nESI). All direct infusion experiments were performed on a Thermo Scientific Q Exactive HF-X Hybrid Quadrupole-Orbitrap Mass Spectrometer. A spray voltage of 0.65-0.8 kV and heated capillary temperature of 150 °C were used to ionize and desolvate the NCB complexes, facilitating their transmission into the gas-phase. In-source CID (80-110 eV) was also utilized to enhance transmission and reduce cationic adduction of the NCB complexes. MS1 spectra were collected at a resolution of 240K (@ *m/z* 200) and averaged over 100 scans (**Supplementary Fig. S21**).

## Acknowledgments

This work was supported by the Welch Foundation to J.S.B. (F-1155) and H.-C.Y. (F-1833), the Texas 4000 to H.-C.Y., National Institutes of Health to H.-C.Y. and I.J.F. (GM129617), and National Science Foundation to H.-C.Y. and J.S.B. (2029266).

## Author contributions

Y.-A.K., C.J., Y.-A.C., J.T.P., I.J.F. and H.-C.Y. discussed and defined the project. I.J.F. and H.-C.Y. supervised the research. Y.-A.K., C.J., Y.-A.C., O.S.Z. and T.N.D. prepared the NCBs and carried out the imaging experiments. Y.-A.K., H.-C.K. prepared PCR samples for the *MiSeq* chip selection. Y.-A.K., O.S.Z., J.A.H and J.R.R. wrote the image alignment and analysis software. Y.-A.K. and O.S.Z. wrote flat-field correction script. Y.-A.K., O.S.Z., D.C.W. and S.H. made a comparison on machine learning algorithms and establish the predictive design workflow. Y.-A.K. and Y.-I.C performed fluorometry data analysis. J.N.W., S.W.S. and J.S.B. conducted direct infusion mass spectrum analysis. Y.-A.K., H.-C.Y., and I.J.F. wrote the paper with editorial assistance from all co-authors.

## Competing interests

The authors declare no competing interests.

## Data availability

CHAMP program is available at https://github.com/finkelsteinlab/champ. All other relevant raw/analyzed data are available from the corresponding authors upon reasonable request.

## Additional information

### Supplementary information

The online version contains supplementary material available at Nature Communications website.

## Notes

### Competing Interest Statement

The authors have declared no competing interest.

