## Supplementary Information for "Massively Parallel Selection of NanoCluster Beacons"

**Contents**

Figure S1: Schematic of NCB working principle……………….………………………..……………...2

Figure S2: Schematic of volumetric integrated intensity………………………………….………….2

Figure S3: *MiSeq* chip images before and after flat-field correction……………..………….…….3

Figure S4: Alignment of PhiX fiducial markers……………………………………………………....4

Figure S5: Identification of activator sequences from the FASTQ data………..…………………5

Figure S6: *MiSeq* chip images before and after restriction enzyme digestion…......……………......6

Figure S7: NCB images before and after restriction enzyme digestion……………………...………7

Figure S8: Illumination comparison of canonical NCB with different activator length…………………8

Figure S9: Intensity comparison of NCBs on the chip under different hybridization temperature…….8

Figure S10: Batch to batch variations in *MiSeq* chip selection for library_1…………………….....9

Figure S11: Batch to batch variations in *MiSeq* chip selection for library_2 and library_3……..…....10

Figure S12: Titration curve of G15 intensity on the chip…………………………..………………11

Figure S13: NCB intensity decay on the chip………………………………………………………11

Figure S14: Influence of activator mutations on NCB brightness ………………………………..…12

Figure S15: Nine-segment interrogation on the library_1 design……………………………..……14

Figure S16: Fluorescence correlation spectroscopy (FCS) results on the red NCBs………………..15

Figure S17: Fluorescence correlation spectroscopy (FCS) results on the yellow NCBs……….…….16

Figure S18: Small-scale test-tube investigation of G15 twin NCBs in the yellow channel..............17

Figure S19: Absorption spectra of selected NCBs and POT……………………..………………18

Figure S20: Native PAGE gel photo under UV excitation…………………….………………...18

Figure S21: ESI-MS analysis of selective NCBs………………….……………………………....19

Figure S22: 2D spectra of bright red activator candidates…………..……………………………....20

Figure S23: 2D spectra of dark activator candidates ………………………………..……………21

Figure S24: 2D spectra of bright yellow activator candidates……………………………..……22

Figure S25: 2D spectra of activators with various numbers of guanine bases……………….…..….23
Figure S26: 2D spectra of rationally designed red NCBs………..…..………………...……....…24
Figure S27: 2D spectra of rationally designed yellow NCBs………..…..………………...……....…25
Figure S28: 2D spectra of randomly designed NCBs ……………………………………….……26
Figure S29: 2D spectra of G5 NCBs……………………………………………………………..……27
Figure S30: 2D spectra of red POT candidates……………….………..……………………………...28
Figure S31: 2D spectra of yellow POT candidates………………..……………………………………29
Figure S32: 2D spectra of red NCBs near rank 3600………….………………………………….…30
Figure S33: 2D spectra of yellow NCBs near rank 3600………...………………………….….…30
Figure S34: Workflow for establishing machine learning models to classify screened NCBs or NCB candidates……………………………………………………………………………………….31
Figure S35: Workflow to rationally design bright NCBs…………………………………….……...32
Figure S36: CHAMP workflow…………………………………………………………………………33
Figure S37: Motif distribution for the top 1,000 library sequences for red and yellow channels...….34
Table S1: Sequences of probes and library designs used in this report…..………..……….…….35
Table S2: Test-tube investigation of selected bright red and dark activator candidates……..….…38
Table S3: Test-tube investigation of selected bright yellow activator candidates…………………....40
Table S4: Test-tube investigation of 10-guanine activators……..…………….…………….…..…41
Table S5: Test-tube investigation of 12-guanine activators ……………………………….………42
Table S6: Test-tube investigation of rationally designed bright red NCBs ……………….….……43
Table S7: Test-tube investigation of rationally designed bright yellow NCBs…………..……..…44
Table S8: Test-tube investigation of randomly designed NCBs and G5…………………….………45
Table S9: Test-tube investigation of red and yellow NCBs ranked near 3600…………….….……46
Table S10: Test-tube investigation of red POT candidates……..……….……...……..…......47
Table S11: Test-tube investigation of yellow POT candidates……..……………….…...…..…...48
Table S12: Machine learning model prediction results………………………………………………49
Table S13: Selected bright and dark features for yellow channel ……………………………...50
Table S14: Selected bright and dark features for red channel…………………………….……...51
References…………………….………………………………………………………………….. 52

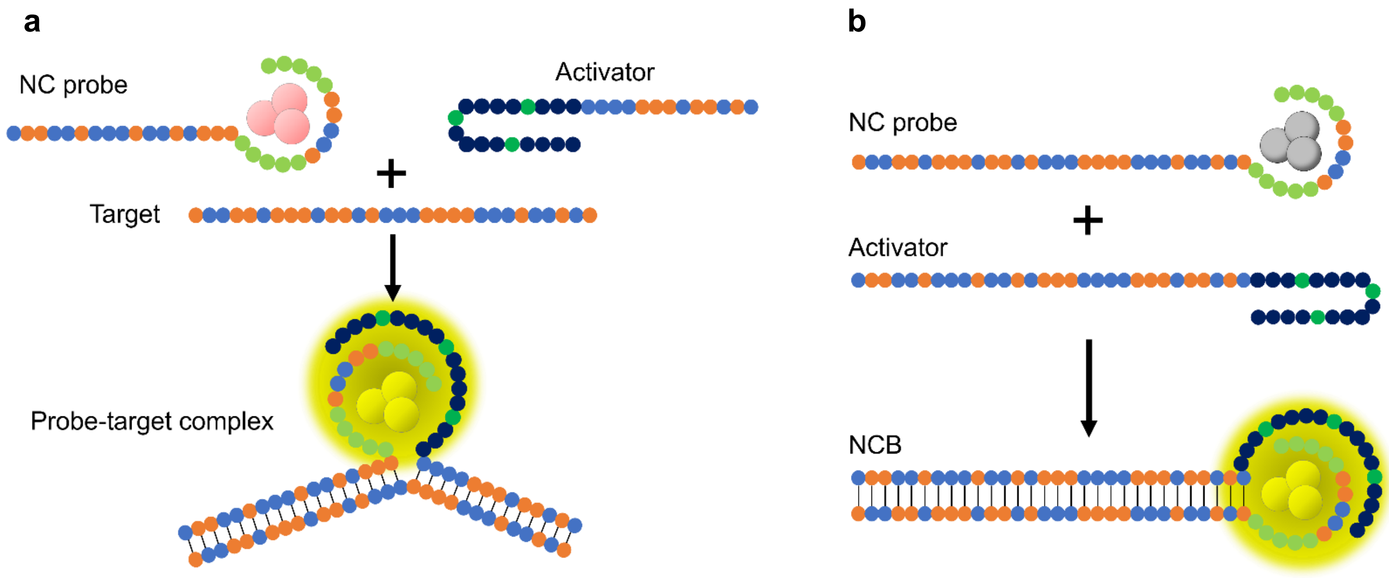


**Figure S1 | Schematic of NanoCluster Beacon (NCB) working principle. a** An NCB consists of an NC probe and an activator probe[1-5]. Upon binding to a target, the dark silver nanocluster (AgNC) interacts with the activator and lights up. NCBs remain dark when there is no target in the solution. **b** In the study, we eliminated the target and only focused on the interactions between the NC probe and the activator.

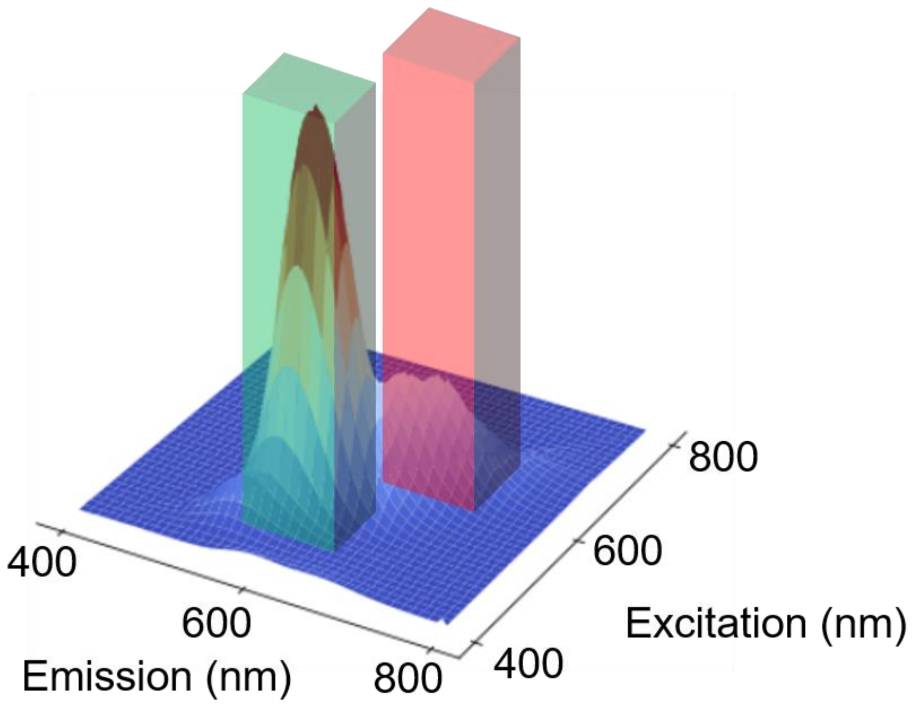


**Figure S2 | Schematic of volumetric integrated intensity.** From each 2D fluorescence spectrum, we can calculate the volumetric integrated intensities in the yellow channel (Ex/Em: 535/50, 605/70 nm) and the red channel (Ex/Em: 620/60, 700/75 nm), respectively. The volumetric integrated intensity refers to the volumetric integral under the 2D spectrum surface.

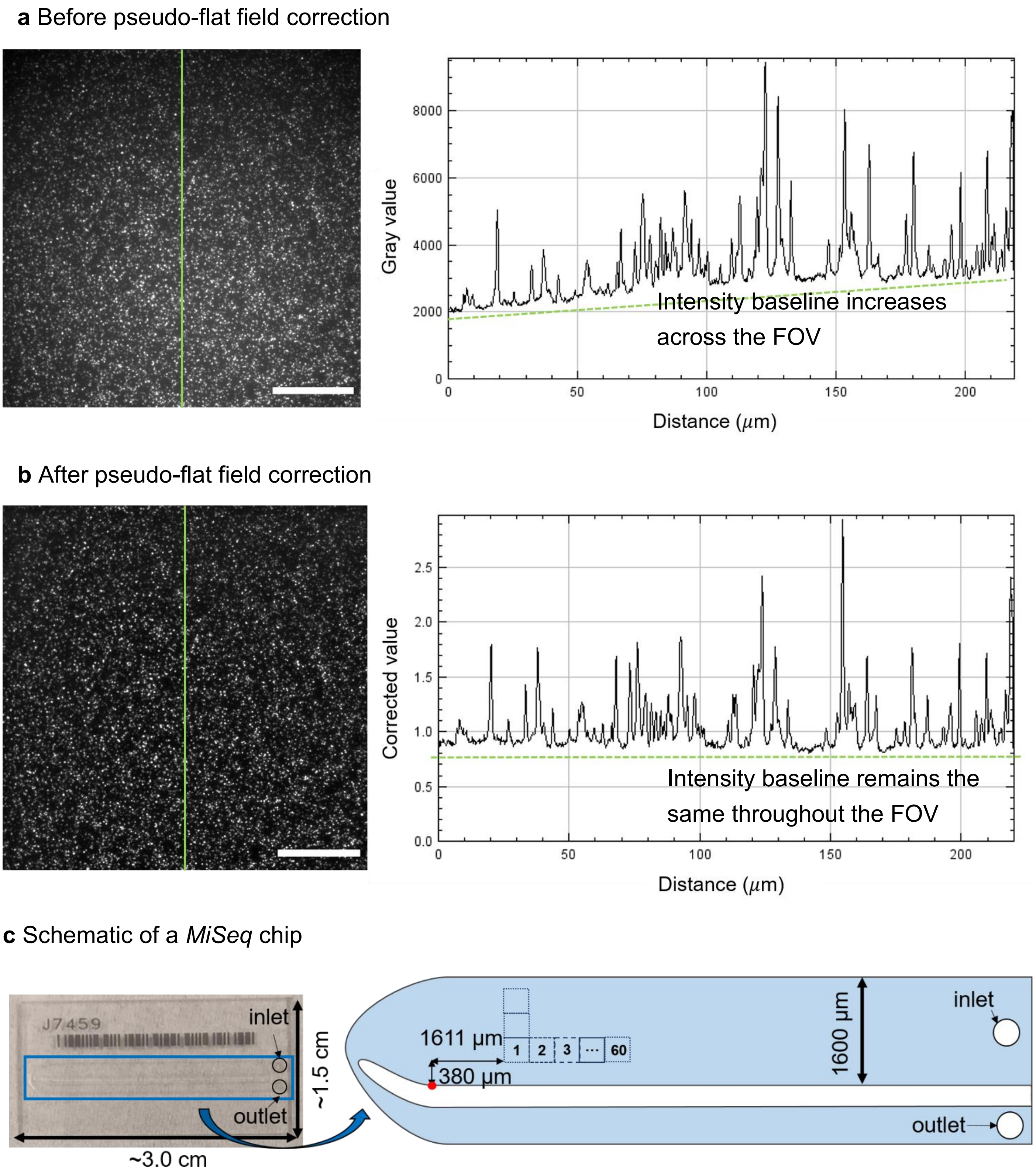


**Figure S3 | *MiSeq* chip images before and after flat-field correction. a** A representative field-of-view (FOV, 1,024×1,024 pixel) shows uneven illumination that leads to inconsistent intensity baseline across the FOV. **b** The same FOV is corrected using a pseudo-flat field correction method. The intensity baseline is uniform throughout the FOV after correction. Distance: from top to bottom. Scale bar: 50 μm. **c** A *MiSeq* chip is 1.5 cm long and 3.0 cm wide (left). To bypass most of these unregistered regions, we shifted the imaging starting position by 380 μm vertically and 1,611 μm horizontally with respect to the reference point (red dot) at the bottom left corner (right).

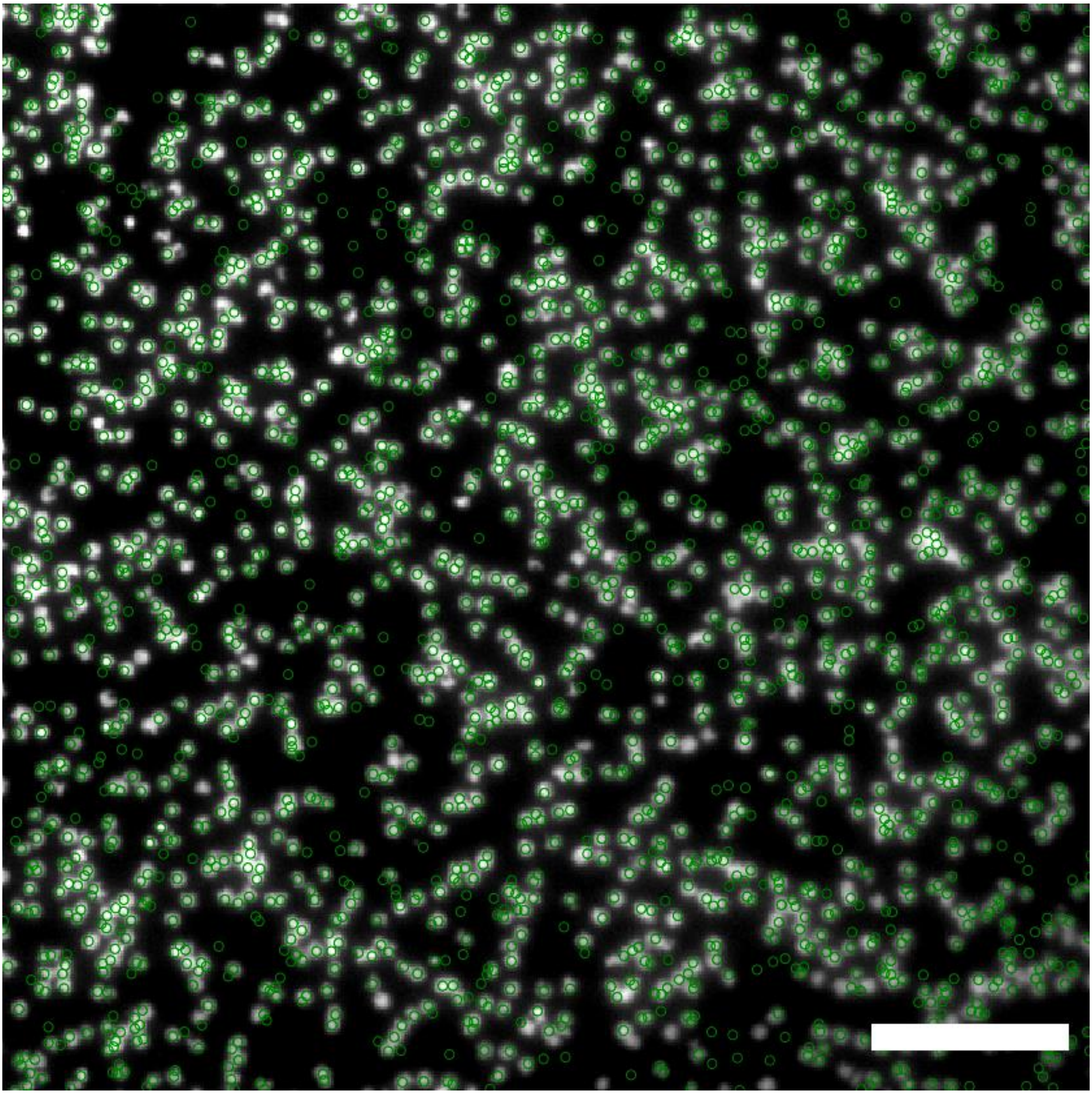

**Figure S4 | Alignment of PhiX fiducial markers.** A representative FOV (512×512 pixel; 110×110 μm$^2$) of aligned PhiX fiducial markers acquired under the green channel (EX/EM: 480/40, 535/50 nm). The PhiX fiducial markers are labeled with Atto488 and the registered PhiX positions from the FASTQ data file are circled in green. Scale bar: 20 μm.

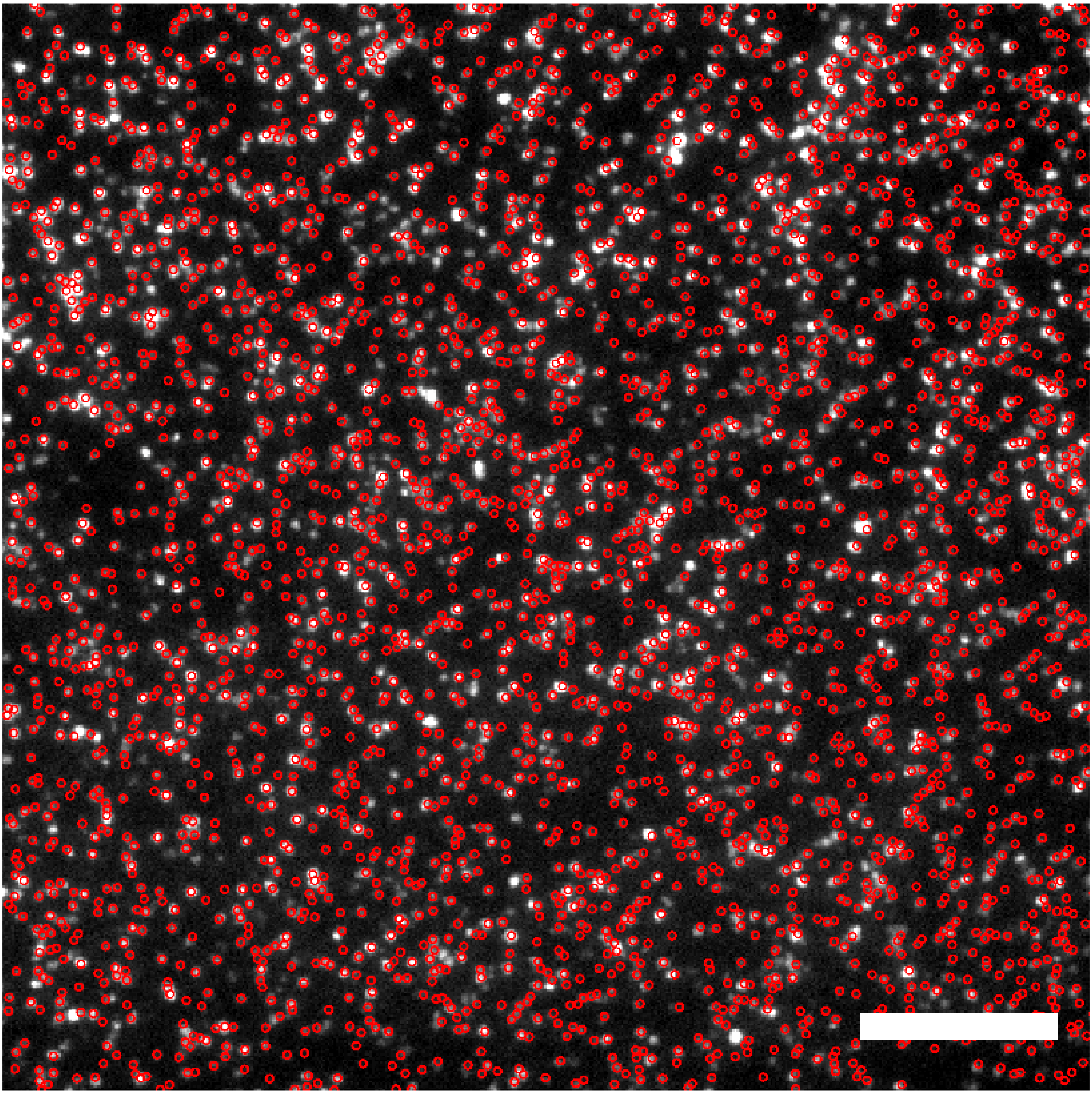

**Figure S5 | Identification of activator sequences from the FASTQ data.** A representative FOV (512×512 pixel; 110×110 $\mu m^2$) of aligned red NCBs polonies acquired under the red channel (EX/EM: 620/60, 700/75). The library sequences (i.e., activators) are hybridized with the common NC probe (i.e., C55) and form activated NCBs on the chip. The registered activator positions from the FASTQ data file are circled in red. These two examples (**Fig. S4** and **S5**) demonstrate the accuracy of the NCB-CHAMP[6] mapping algorithm. Scale bar: 20 μm.

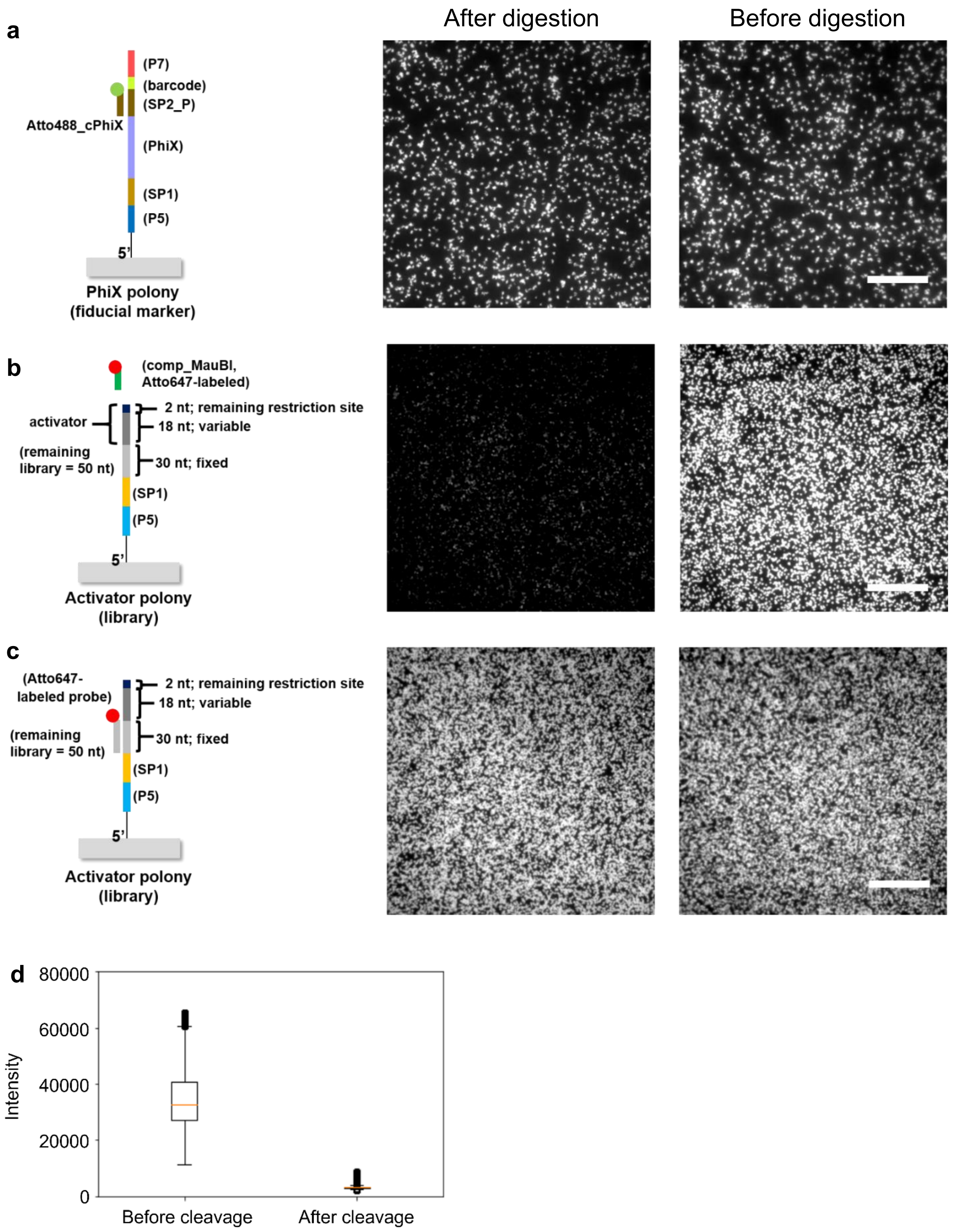


**Figure S6 | *MiSeq* chip images before and after restriction enzyme digestion. a** PhiX image is unaffected after cleavage. **b** Overhang is successfully cleaved as minimal Atto647N-labeled comp_SP2 probes can still bind with the activator polonies. **c** The library sequences are unaffected after digestion. **d**. We averaged one row of Atto647-tagged fluorescence images. The median intensity dropped ~90% after cleavage. Box plots represented median and 25th and 75th percentiles—interquartile range; IQR—and whiskers extended to 1.5× IQR from the hinges. Empty circles represented the outliers. **b** and **c** panels had the same contrast setting, while the contrast setting of **a** panel was different. Scale bar: 25 μm.

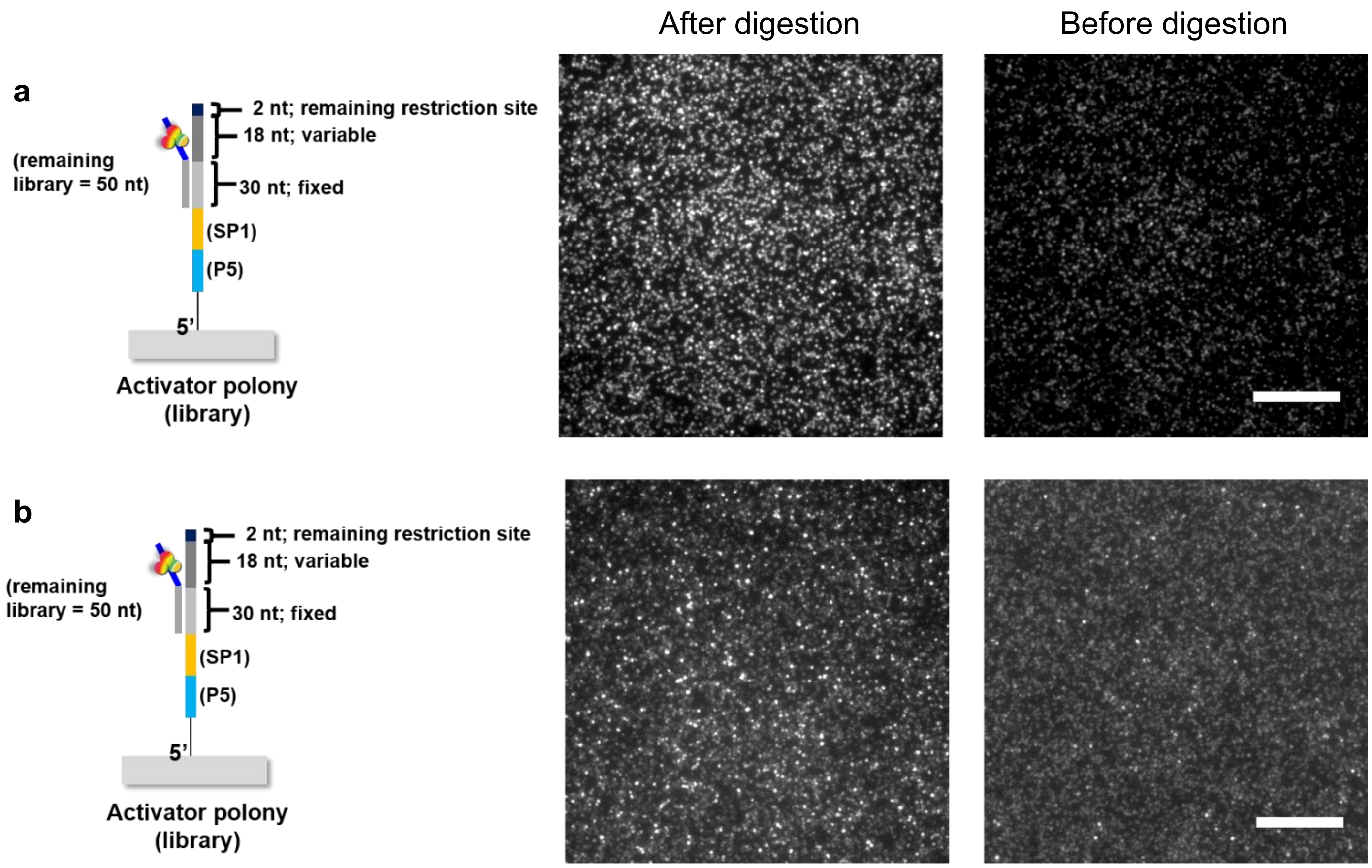


**Figure S7 | NCBs images before and after digestion. a** After restriction enzyme digestion, stronger NCBs signals are observed in the red channel. **b** Stronger NCBs signals are observed in the yellow channel. **a** and **b** panels had the same contrast setting. Scale bar: 25 μm.

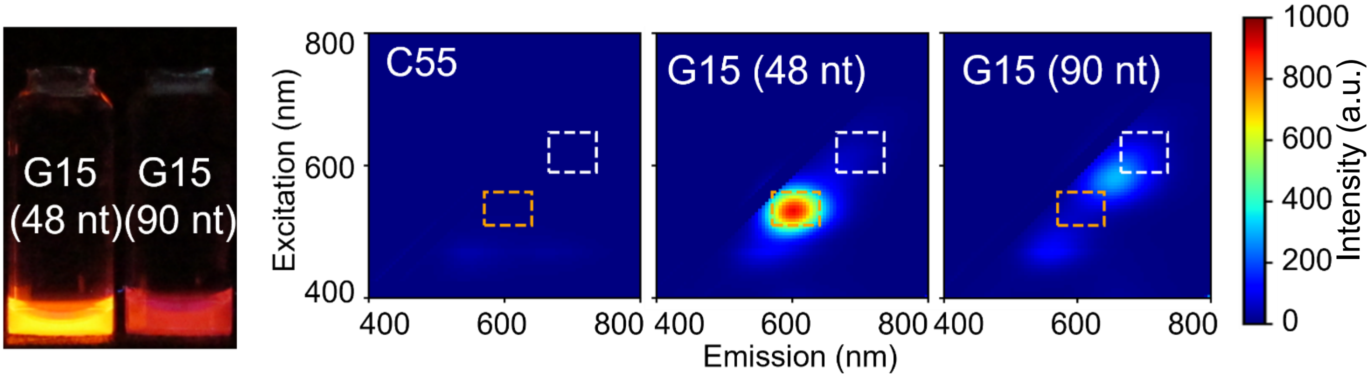


**Figure S8 | Illumination comparison of canonical NCBs with different activator length.** We observed red-shifted fluorescence for NCBs having longer G15 activator (90-nt long) compared to canonical G15 activator (48-nt long).

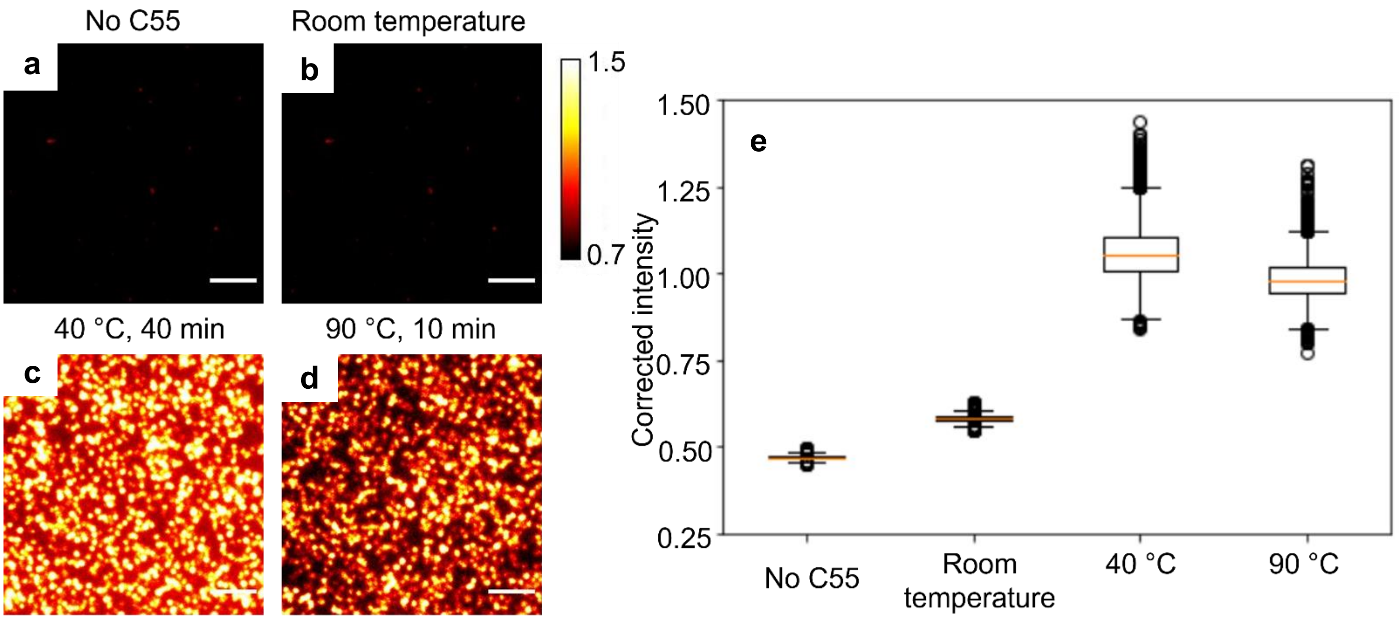


**Figure S9 | Intensity comparison of NCBs on the chip under different hybridizing temperature.** We tested the NCBs intensity on the chip with different hybridizing temperature. We found that 40°C gave the brightest mean intensity compared to 90 °C, which was the condition similar as test-tube validation. Furthermore, as 40 °C gave a more moderate condition to the delicate *MiSeq* chip, we applied 40 °C to the chip experiments throughout this report. Box plots represented median and 25th and 75th percentiles—interquartile range; IQR—and whiskers extended to 1.5× IQR from the hinges. Empty circles represented the outliers.

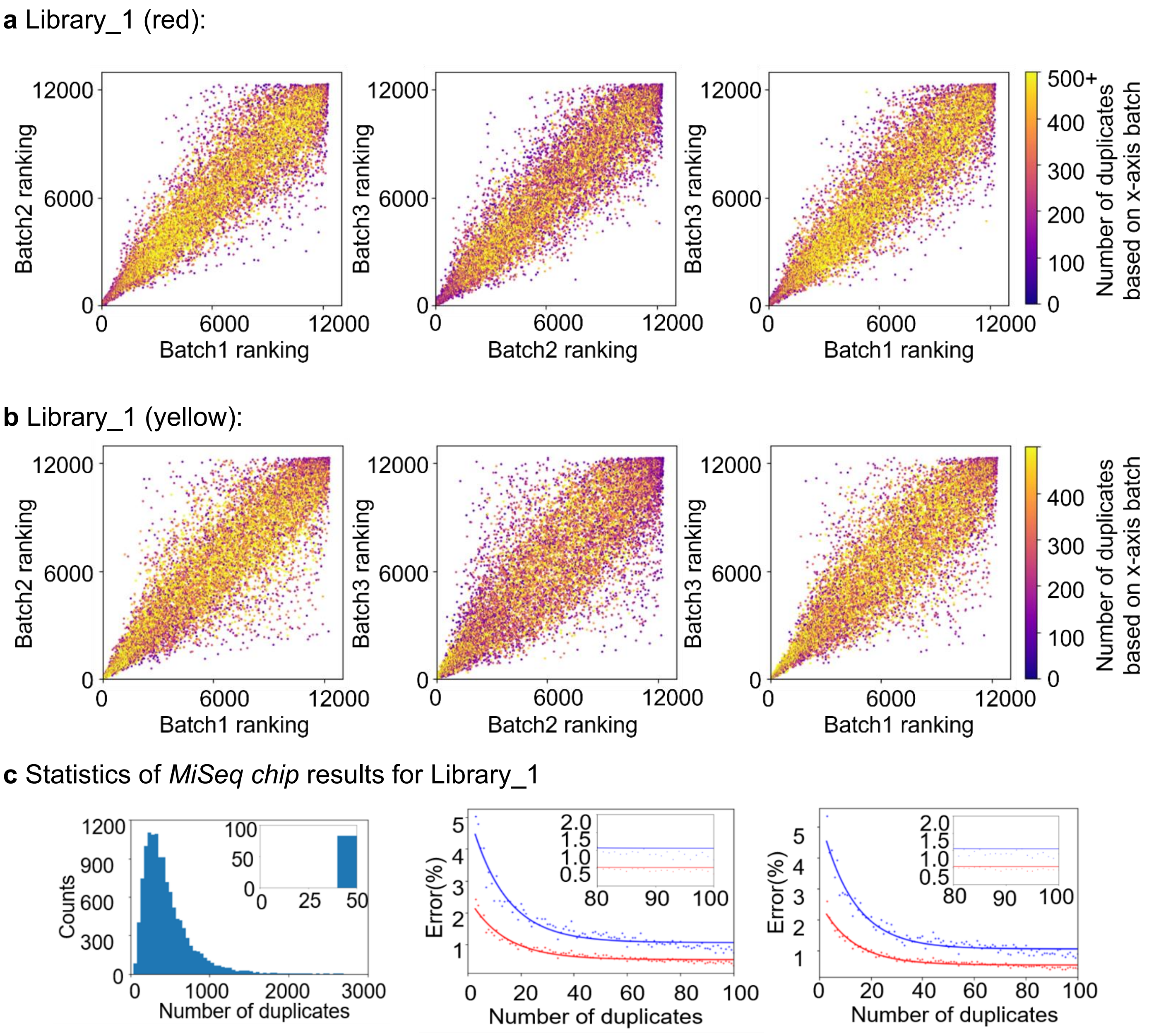


**Figure S10 | Batch to batch variations in *MiSeq* chip selection for library_1.** Nonparametric measure is applied to evaluate the ranking correlation among repeated experiments. **a** For library_1, all 12,286 distinct sequences are found on chip. The Spearman's rho for the 3 comparisons in red channel are 0.92, 0.93, and 0.93, with the $R^2$ of 0.85, 0.86 and 0.86 (left to right). **b** The Spearman's rho for the 3 comparisons in yellow channel are 0.86, 0.91, and 0.86, with the $R^2$ of 0.75, 0.83 and 0.75 (left to right). **c** (left) Distribution of the number of activator duplicates in library_1. All activators had at least 20 duplicates observed. On average, each activator had 457 ± 308 polonies on a *MiSeq* chip. (middle and right) Estimation of error in the NCB brightness characterization after bootstrapping. To improve the accuracy of our high-throughput screening, we performed 100 rounds of bootstrapping processes by random sampling 75% of observed duplictaes intensity and assigned median intensity as the NCB on-chip intensity. Bootstrap intensity values were calculated for the standard sequence (i.e., G15) with all numbers of clusters between 3 and 100. Shown are the average errors (red points) and 90% confidence intervals of error (blue points), using the median intensity with either 200 (middle) or 20,000 clusters (right) for 10,000 rounds as reference. Solid lines indicate a fit to the data.

**a** Library_2 (red):

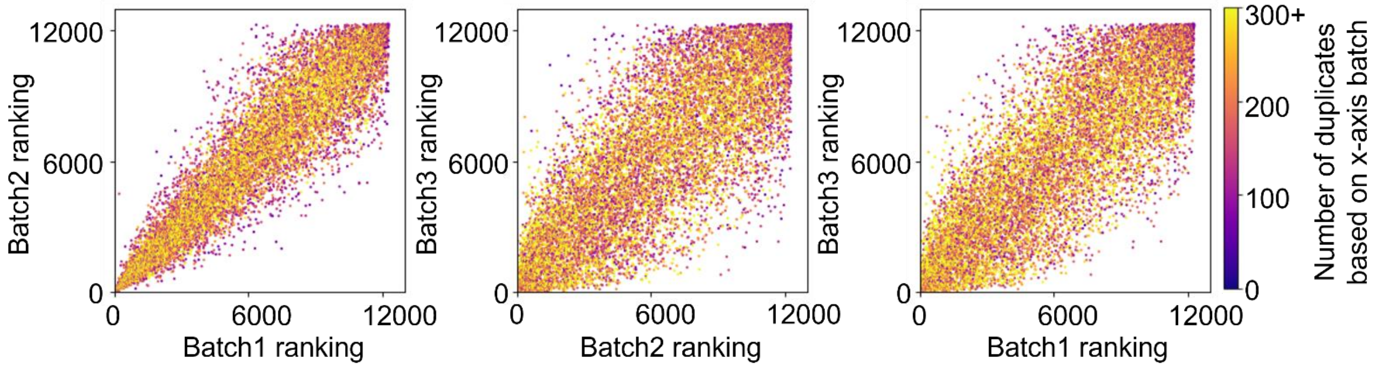


**b** Library_3 (red):

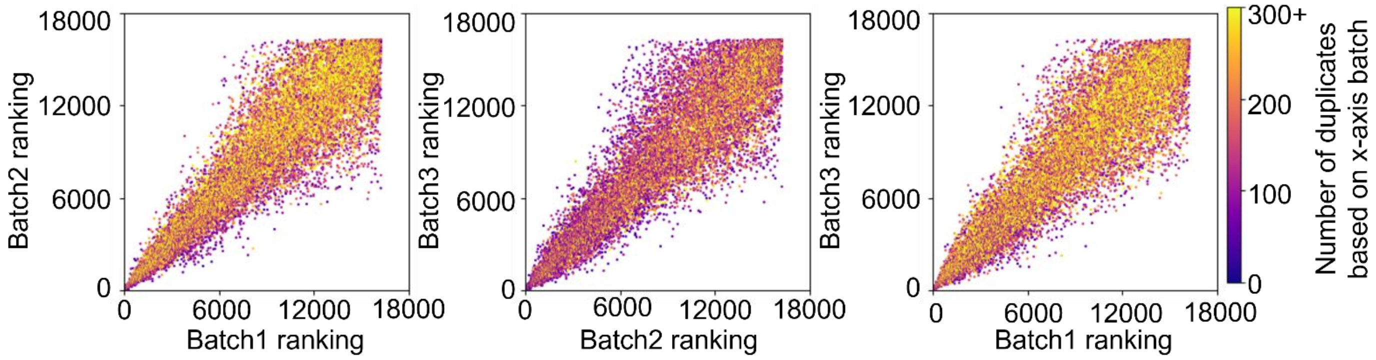


**Figure S11 | Batch to batch variations in *MiSeq* chip selection for library_2 and library_3. a** For library_2, all 12,286 distinct sequences are found on chip. The Spearman's rho for the 3 comparisons in red channel are 0.93, 0.83 and 0.80 (left to right), with the $R^2$ of 0.87, 0.68, and 0.65 (left to right). **b** For library_3, all 16,255 distinct sequences are found on chip. The Spearman's rho for the 3 comparisons in red channel are 0.91, 0.86 and 0.89 (left to right), with the $R^2$ of 0.82, 0.73 and 0.79 (left to right).

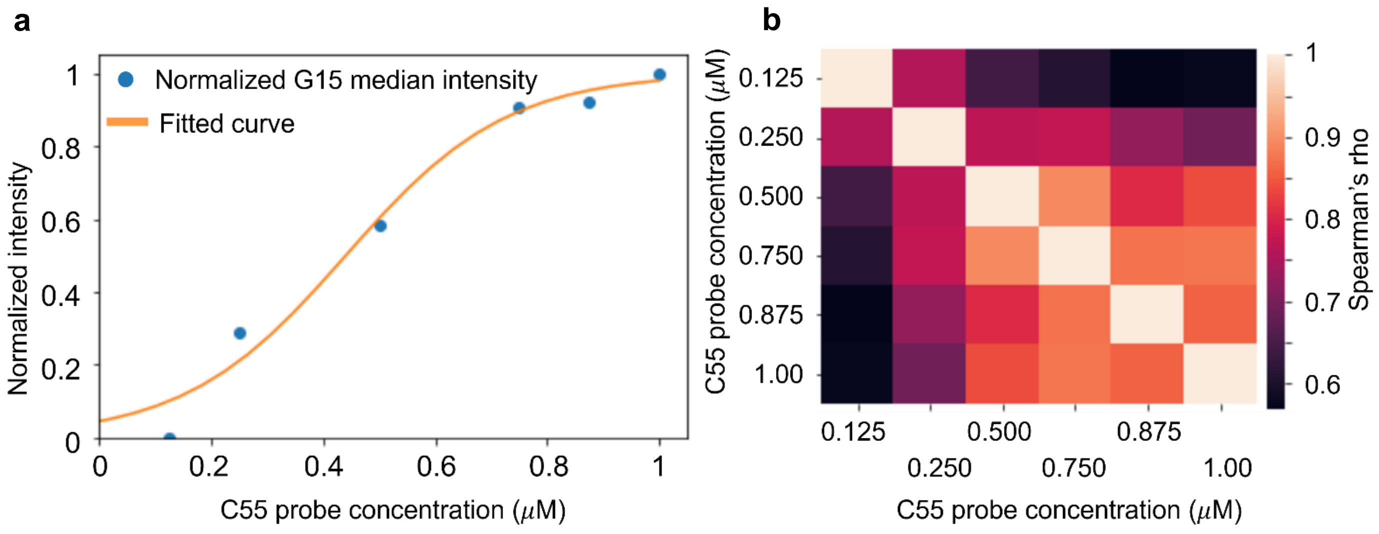


**Figure S12 | Titration curve of G15 NCB intensity on the chip. a** To find out the optimal condition for NCB screening on *MiSeq* chip, we used the G15 NCB intensity as the calibration standard in a titration experiment. C55 probes at 6 different concentrations were delivered to the chip. The normalized G15 NCB intensity reached a plateau when the C55 probe concentration was about 0.8 μM. **b** However, we also observed highly consistent ranking results if C55 probe concentrations were higher than 0.5 μM.

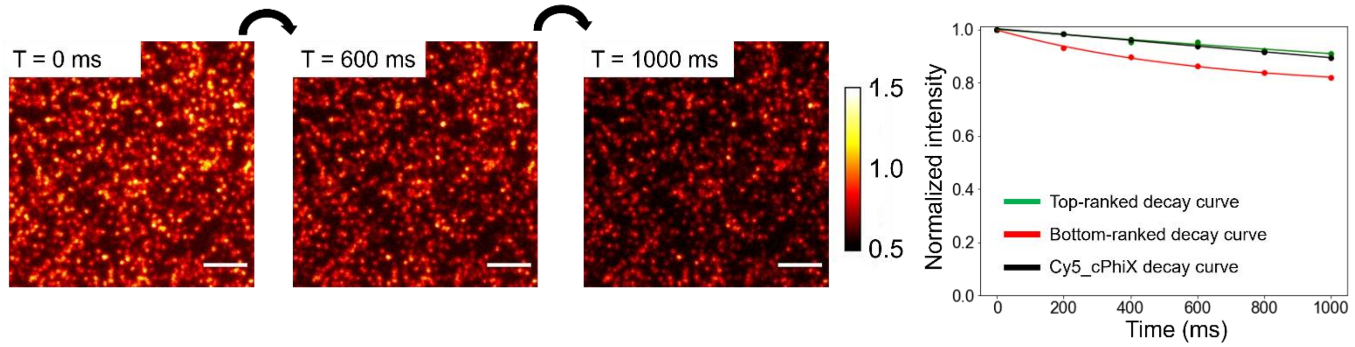


**Figure S13 | NCB intensity decay on the chip.** By acquiring a fluorescence image every 200 ms, intensity time traces of polonies were obtained, which could be fitted with a single-exponential decay. After one second of strong illumination (~10 W/cm$^2$), polony intensity decreased by ~20% at most.

**a** Library_1, 3- and 6-segment interrogation on yellow NCBs

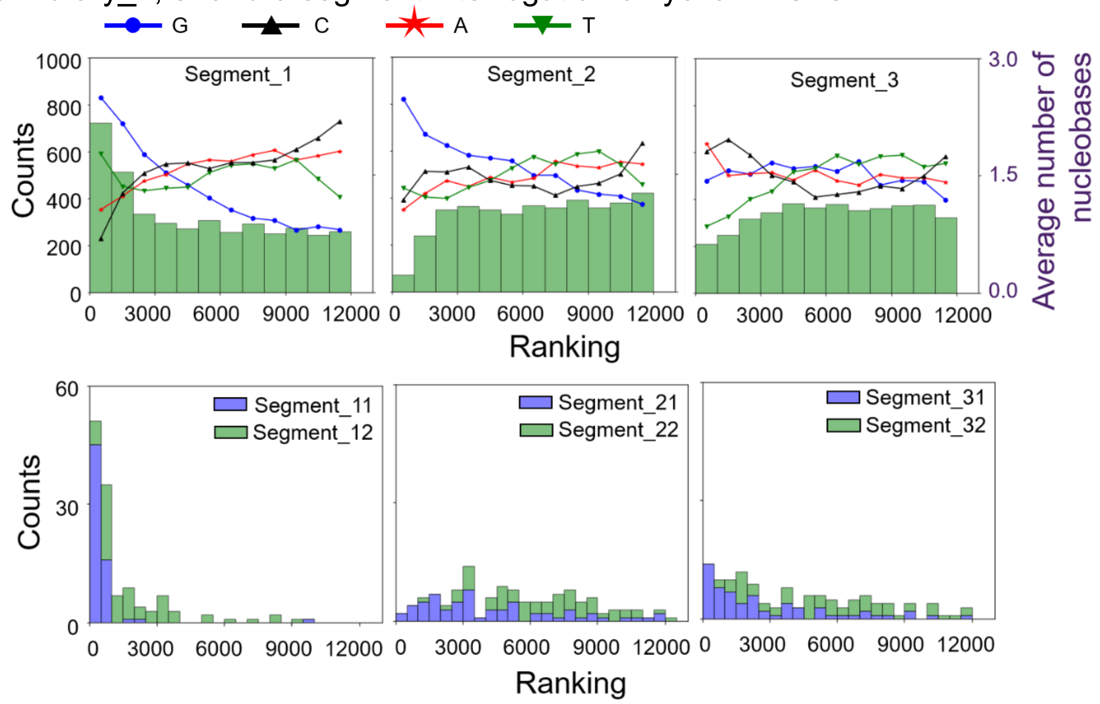


**b** Library_2, 3-segment interrogation on red NCBs

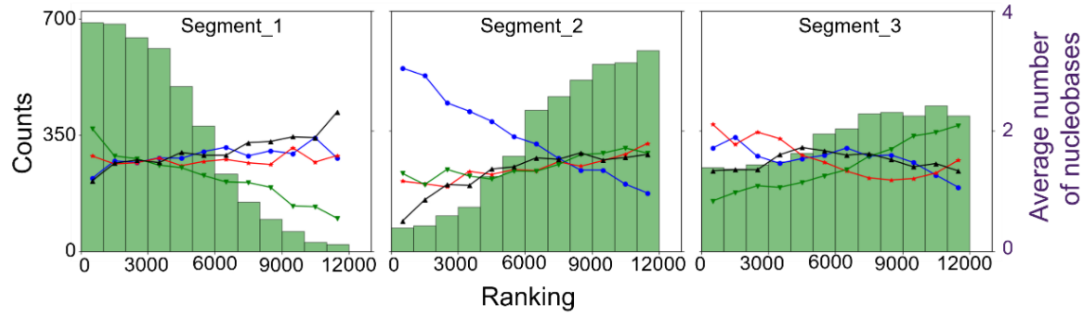


**c** Library_3, 4-segment interrogation on red NCBs

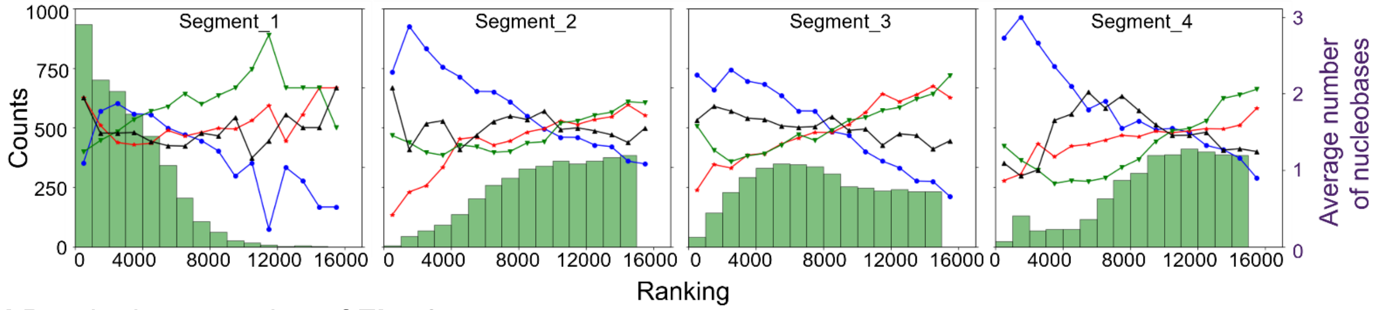


**d** Detailed explanation of **Fig. 2c**

**Activator1:**
ATCCGTGGTGGGGTGGGG
Number of adenine = 1
Number of thymine = 2
Number of guanine = 1
Number of cytosine = 2

**Activator2:**
ACCATTGGTGGGGTGGGG
Number of adenine = 2
Number of thymine = 2
Number of guanine = 0
Number of cytosine = 2

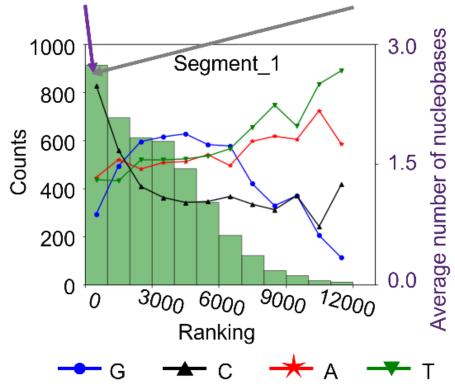


Average number of nucleobases = $\left(\frac{\Sigma_i(\text{number of nucleobases})_{\text{activator } i}}{\text{total number of activators considered}}\right)$

Average number of adenines = $\frac{1+2}{2}$ =1.5

Average number of thymines = $\frac{2+2}{2}$ =2

Average number of guanines = $\frac{1+0}{2}$ =0.5

Average number of cytosines = $\frac{2+2}{2}$ =2

**Figure S14 | Influence of activator mutations on NCB brightness. a** Here we showed the influence of activator mutations on yellow NCB brightness. **b** When randomizing the 3 segments in library_2, positions 10 to 15 were found to be the interaction hot zone. The segment definition for all three libraries could be found in **Table S1c**. **c** When randomizing the 4 segments in library_3, positions 10 to 12 were found to be the interaction hot zone. The results from (**b**) and (**c**) were consistent with the library_1 result (**Fig. 2**), which showed positions 10-12 were the interaction hot zone for creating bright NCBs. **d** Here we demonstrated calculation of the average number of nucleobases from two top-ranked sequences. For instance, Activator1 (Left, ATCCGT GGTGGG GTGGGG, rank no.12) has 1 adenine, 2 thymine, 1 guanine, and 2 cytosine, while the Activator2 (Right, ATCCGT GGTGGG GTGGGG, rank no. 14) has 2 adenine, 2 thymine, 0 guanine, and 2 cytosine. Both activators were highly ranked (i.e., they create bright NCBs) and located in the first bin (ranking 1-1,000) of the histogram. Each histogram contained 4,096 sequences.

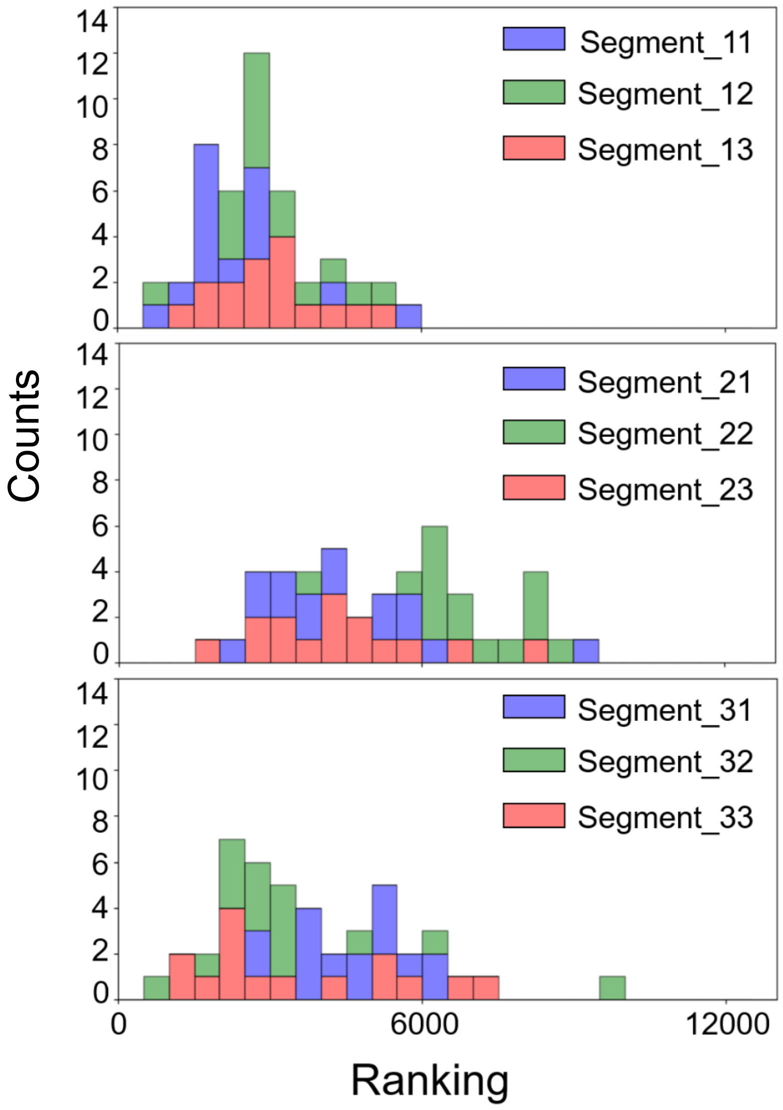


**Figure S15 | Nine-segment interrogation on the library_1 for red NCB brightness.** Here we further divide the library_1 activator into 9 segments (**Table S1b**) and investigate each segment's influence on red NCBs brightness. In segment_2, segment_22 (positons 9-10) and segment_23 (positions 11-12) are the critical zones as the ranking shifts toward the dark side when these segments are randomized.

**a**

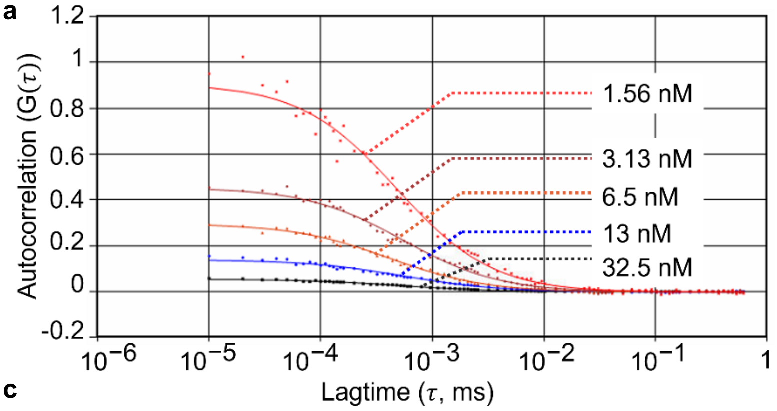


**b**

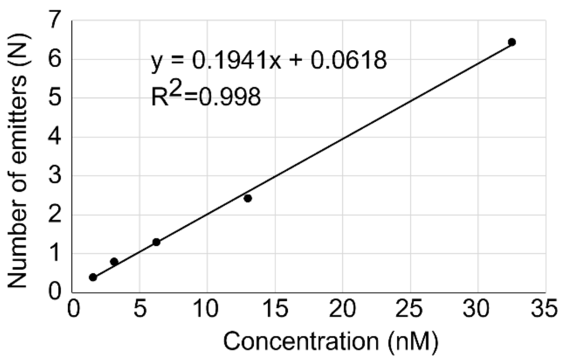


**c**

| Concentration (nM) | N | Avg. p.c. (kHz) | Tau ($\mu s$) | Single-emitter brightness (SEB, kHz) | Chi-square |
|---|---|---|---|---|---|
| 32.50 | 6.43 | 54.12 | 539.07 | 8.42 | 0.42 |
| 13.00 | 2.42 | 15.54 | 559.56 | 6.42 | 0.63 |
| 6.25 | 1.29 | 6.43 | 521.28 | 4.98 | 0.52 |
| 3.13 | 0.79 | 5.98 | 547.22 | 7.57 | 0.50 |
| 1.56 | 0.38 | 1.99 | 473.14 | 5.24 | 0.53 |
| 0.78 | 0.17 | 0.95 | 488.85 | 5.59 | 0.58 |

**d**

| ID | N | Avg. p.c. (kHz) | Dilution times | Tau ($\mu s$) | Chi-square | SEB (kHz) | Yield (%) | SEB*yield (kHz·%) | Enhancement ratio |
|---|---|---|---|---|---|---|---|---|---|
| rPOT5 NCB | 4.91 | 27.82 | 146.61 | 741.26 | 0.77 | 5.67 | 19.96 | 113.17 | 476 |
| rPOT6 NCB | 5.64 | 19.48 | 23.31 | 752.17 | 0.54 | 3.45 | 3.60 | 12.42 | 51 |

**e**

| | |
|---|---|
| **SEB improvement** | 5.67 (kHz) / 3.45 (kHz) **~1.64-fold** |
| **NCB chromophore chemical yield improvement** | 0.1996/0.0360 **~5.54** |
| **SEB*yield improvement** | 113.17 (kHz·%) / 12.42 (kHz·%) **~9.11** |
| **POT difference ratio in red channel (Bright/Dark)** | 476/51 **~9.33** |

**Avg. p. c.** (average photon count); **N** (number of emitters in detection volume);

**Tau** (dwell time of emitters in detection volume); **SEB** (single-emitter brightness)

**Figure S16 | Fluorescence correlation spectroscopy (FCS) results on the red NCBs. a** The amplitude of autocorrelation function, *G(0)*, is inversely proportional to the fluorophore concentration (Atto647N-labeled ssDNA, for calibration purpose), demonstrating no optical saturation in our FCS experiments. **b** Number of emitters in the detection volume (1/*G(0)*) shows a linear relationship with the emitter concentration, generating a calibration curve. **c** The fitting parameters of the calibration FCS experiment on Atto647N-labeled ssDNA. **d** The fitting parameters of the FCS experiment on rPOT5 and rPOT6 NCBs, which are an extreme POT (**Fig. 3**). **e** From the FCS experiment, it is clearly to see that a single rPOT5 emitter is 1.64-fold brighter than a single rPOT6 emitter (which we term “single-emitter brightness”, SEB), and the concentration of rPOT5 emitter is 5.54 higher than that of rPOT6 emitter (which we call “chromophore chemical yield” or “yield” – not all NC probes carry a AgNC that can be activated). The product of SEB improvement and chromophore chemical yield improvement (9.11) is about the same as the improvement in ensemble enhancement ratio identified by the fluorometer (9.33), indicating that the intensity difference seen in rPOT NCBs is a result of different yield and different SEB. For FCS setup and analysis, please refer to ***Methods.***

a

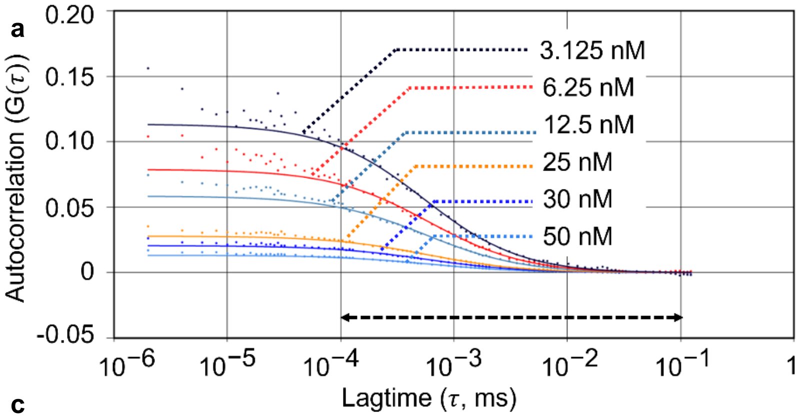


b

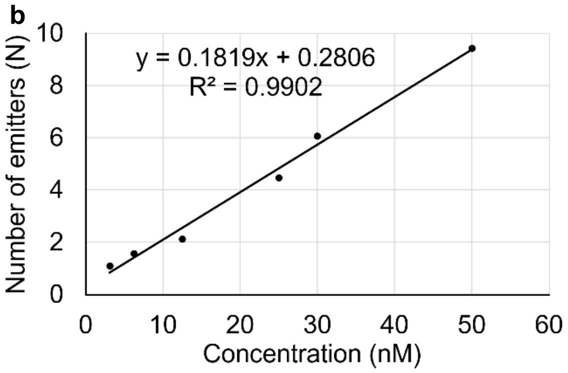


c

| Concentration (nM) | N | Avg. p.c. (kHz) | Tau ($\mu s$) | Single-emitter brightness (SEB, kHz) | Chi-square |
|---|---|---|---|---|---|
| 50 | 9.43 | 287.44 | 567.43 | 30.48 | 0.36 |
| 30 | 6.07 | 175.05 | 589.78 | 28.86 | 0.47 |
| 25 | 4.47 | 135.76 | 584.26 | 30.34 | 0.44 |
| 12.5 | 2.12 | 60.48 | 592.97 | 28.51 | 0.67 |
| 6.25 | 1.57 | 32.31 | 585.83 | 20.56 | 0.52 |
| 3.13 | 1.10 | 18.47 | 574.97 | 16.81 | 0.55 |

d

| ID | N | Avg. p.c. (kHz) | Dilution times | Tau ($\mu s$) | Chi-square | SEB (kHz) | Yield (%) | SEB*yield (kHz·%) | Enhancement ratio |
|---|---|---|---|---|---|---|---|---|---|
| yPOT5 NCB | 7.32 | 51525.4 | 100 | 1051.78 | 0.70 | 7.04 | 25.80 | 181.55 | 982 |
| yPOT6 NCB | 4.60 | 14848 | 10 | 779.71 | 0.70 | 3.24 | 1.58 | 5.11 | 28 |

e

| | |
|---|---|
| **SEB improvement** | 7.04 (kHz) / 3.24 (kHz) **~2.17-fold** |
| **NCB chromophore chemical yield improvement** | 0.2579/0.0158 **~16.33** |
| **SEB*yield improvement** | 181.55 (kHz·%) / 5.11 (kHz·%) **~35.53** |
| **POT difference ratio in yellow channel (Bright/Dark)** | 982/28 **~35.07** |

**Avg. p. c.** (average photon count); **N** (number of emitters in detection volume);

**Tau** (dwell time of emitters in detection volume); **SEB** (single-emitter brightness)

**Figure S17 | Fluorescence correlation spectroscopy (FCS) results on the yellow NCBs. a** The amplitude of autocorrelation function, *G(0)*, is inversely proportional to the fluorophore concentration (Atto532N-labeled ssDNA, for calibration purpose), demonstrating no optical saturation in our FCS experiments. **b** Number of emitters in the detection volume (1/*G(0)*) shows a linear relationship with the emitter concentration, generating a calibration curve. **c** The fitting parameters of the calibration FCS experiment on Atto532N-labeled ssDNA. **d** The fitting parameters of the FCS experiment on yPOT5 and yPOT6 NCBs, which are an extreme POT (**Fig. 3**). **e** From the FCS experiment, it is clearly to see that the SEB of yPOT5 emitter is 2.17-fold brighter than that of yPOT6 emitter, and the chromophore chemical yield of yPOT5 emitter is 16.33 higher than that of yPOT6 emitter. The product of SEB improvement and chromophore chemical yield improvement (35.53) is about the same as the improvement in ensemble enhancement ratio identified by the fluorometer (35.07), indicating that the intensity difference seen in yPOT NCBs is a result of different yield and different SEB. For FCS setup and analysis, please refer to ***Methods***.

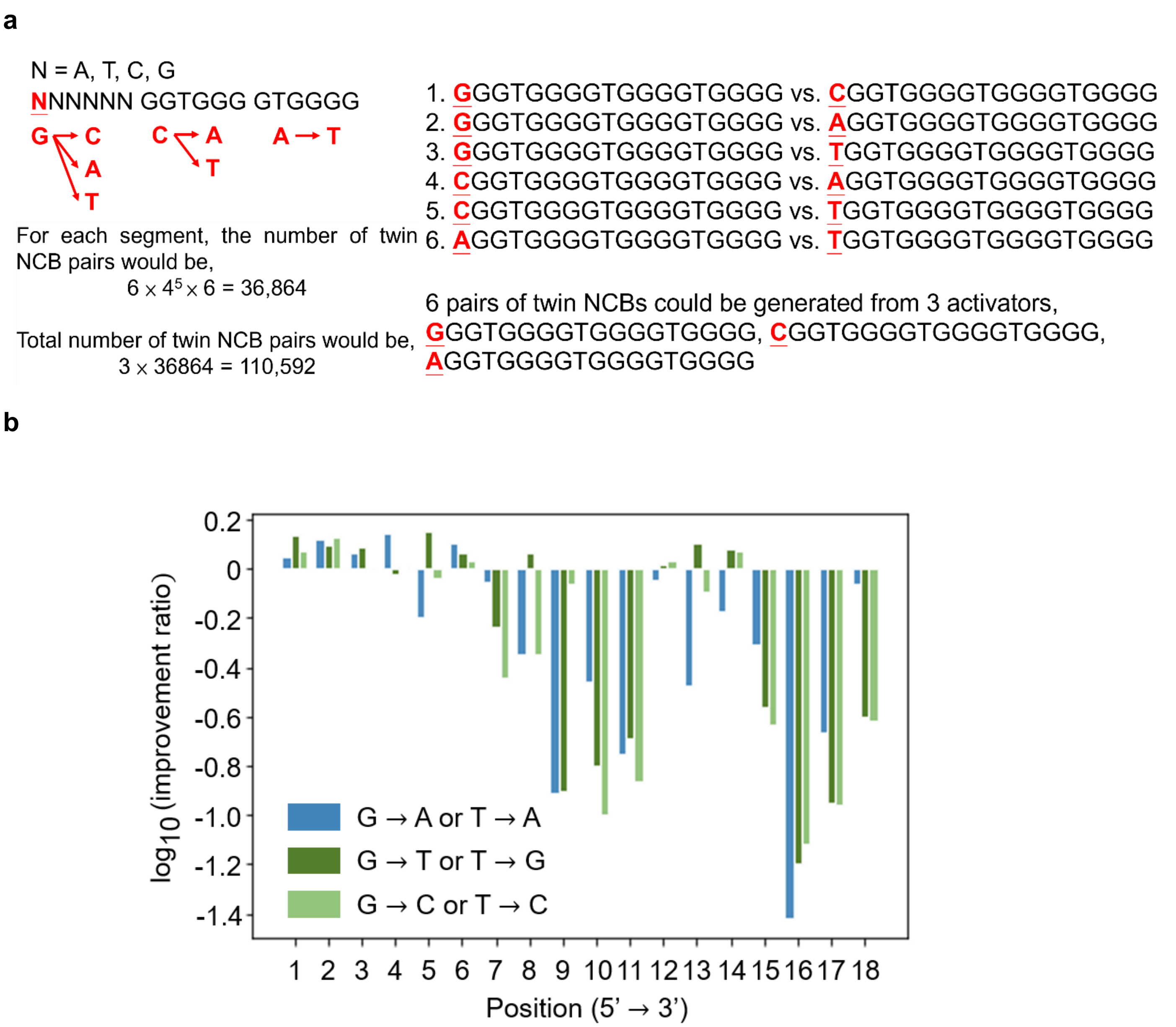


**Figure S18 | Calculation for the number of twin NCB pairs in library_1 and small-scale test-tube investigation of G15 twin NCBs in the yellow channel. a** Although library_1 only has 12,286 activators, they give us totally 110,592 twin NCB pairs. For each position in segment_1, there are 6 scenarios for creating twin NCBs at that position. Considering we have 6 positions in segment_1 and we fill up the rest of the 5 positions using the $4^5$ combinations, we have $6\times4^5\times6 = 36,864$ distinct twin NCB pairs just for segment_1. For 3 segments, there are totally $3\times36,864 = 110,592$ distinct twin NCB pairs in library_1. **b** The improvement ratios of 54 G15 twin NCBs are put into this base-10 logarithm chart. By substituting G to T at position 5 (i.e., GGGTTGGGTGGGGTGGGG), the largest improvement in the enhancement ratio is observed, which is only 1.41-fold higher than the enhancement ratio of G15 NCB in the yellow channel. This result demonstrates that a small-scale investigation cannot improve the brightness of an existing NCB by more than 2-fold. Interestingly, by substituting G to A at position 16 (i.e., GGGTGGGGTGGGGTGAGG), we observed a pair of POTs with POT difference ratio ~25.

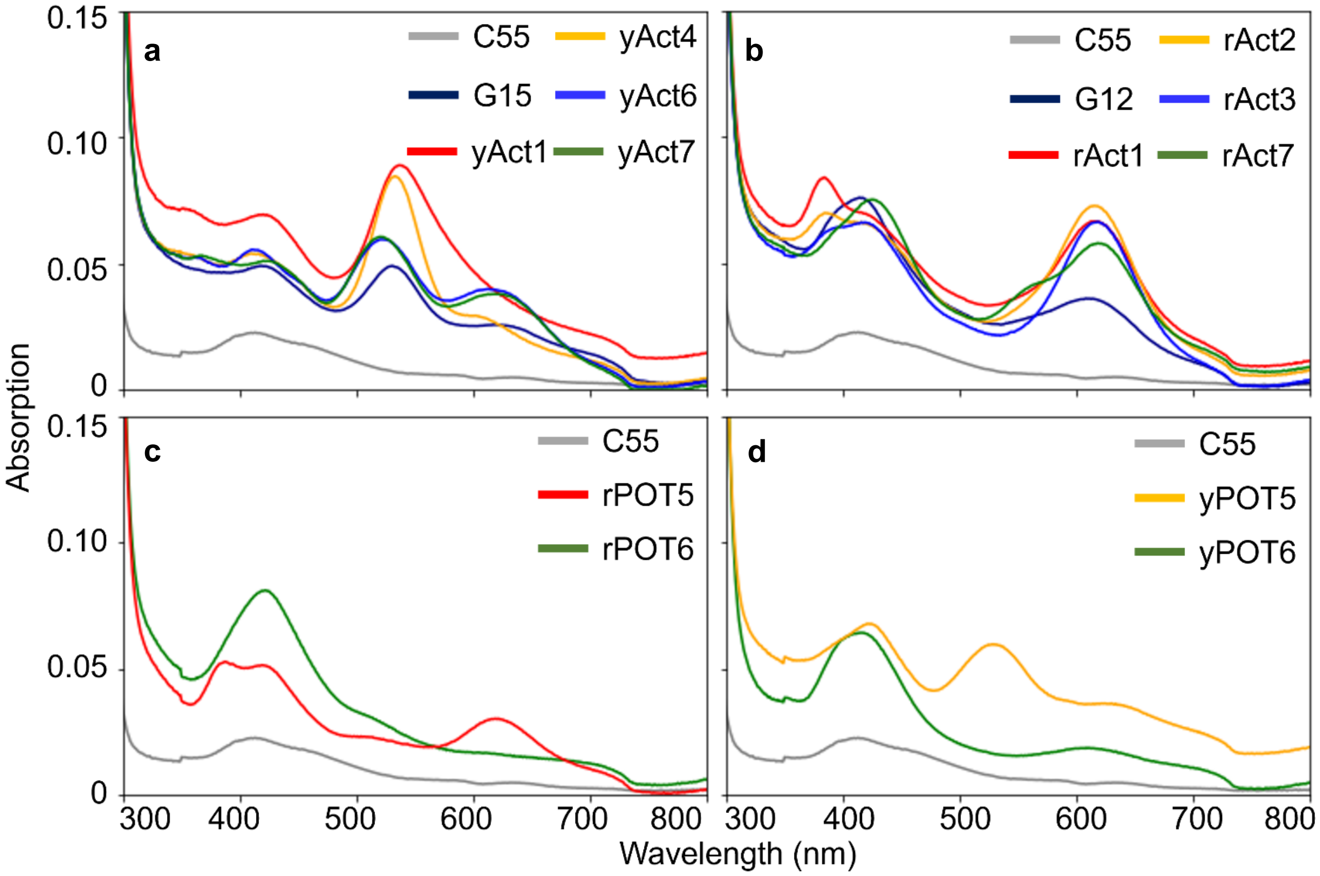


**Figure S19 | Absorption spectra of selected NCBs and POTs. a** For the 5 selected yellow NCBs, we observed the highest absorbance (0.089) for yAct1 NCB at 535 nm, while that of the 5 selected red NCBs reached 0.073 at 610 nm for rAct2 as shown in (**b**). **c** For rPOT5 and rPOT6 NCBs, differences in their absorption spectra around 610 nm were observed (0.030 for rPOT5 and 0.017 for rPOT6). **d** For yPOT5 and yPOT6 NCBs, differences in their absorption spectra around 530 nm were observed (0.060 for yPOT5 and 0.015 for yPOT6).

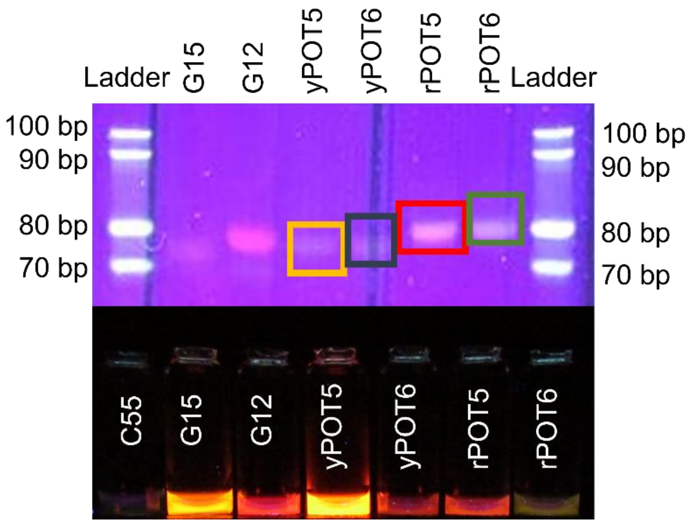


**Figure S20 | Native PAGE gel photo under UV excitation.** We assessed the mobility of selected NCBs using 20% native PAGE gel. Four of the NCBs (yPOT5, yPOT6, rPOT5 and rPOT6, highlighted by color boxes) were selected to process purification and ESI-MS analysis afterward.

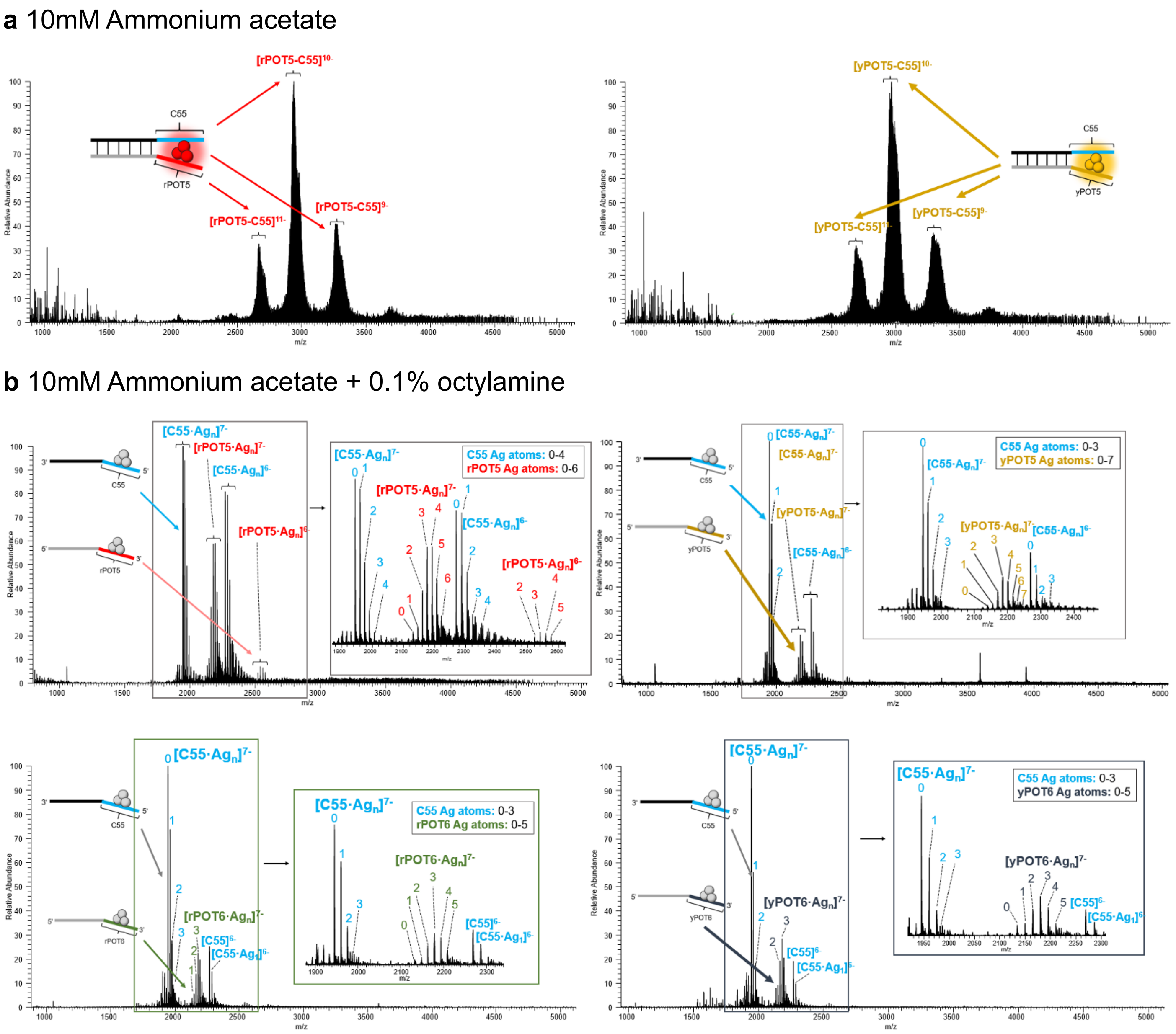


**Figure S21 | ESI-MS analysis of selective NCBs. a** Following the purification using gel electrophoresis, several NCB samples (yPOT5-C55, rPOT5-C55, yPOT6-C55 and rPOT6-C55) were desalted and buffer exchanged into 10 mM ammonium acetate. The resultant mixture was then analyzed by ESI-MS to evaluate the silver stoichiometry of the NCB complexes. **b** Octylamine was added to aliquots of yPOT5-C55 and rPOT5-C55 NCBs (upper row), yPOT6-C55 and rPOT6-C55 (bottom row) at a concentration of 0.1% (v/v), in attempt to reduce the extensive cationic metal adduction that is commonly seen for ESI-MS analysis of oligonucleotides > 20 nt[40-43]. Although the addition of octylamine disassembles the NCB complexes into their respective NC probe and an activator sequence, we found the C55 NC probes from the yPOT5 NCB carry 0-4 and the others carry 0-3 silver atoms, while the activators from yPOT5, rPOT5, yPOT6 and rPOT6 NCBs carry 0-7, 0-6, 0-5 and 0-5 silver atoms, respectively. These results indicate that the original silver stoichiometry for the intact yPOT5 NCB may be larger than that of the intact rPOT5 NCB. Moreover, the bright member of POTs may be larger than its counterpart.

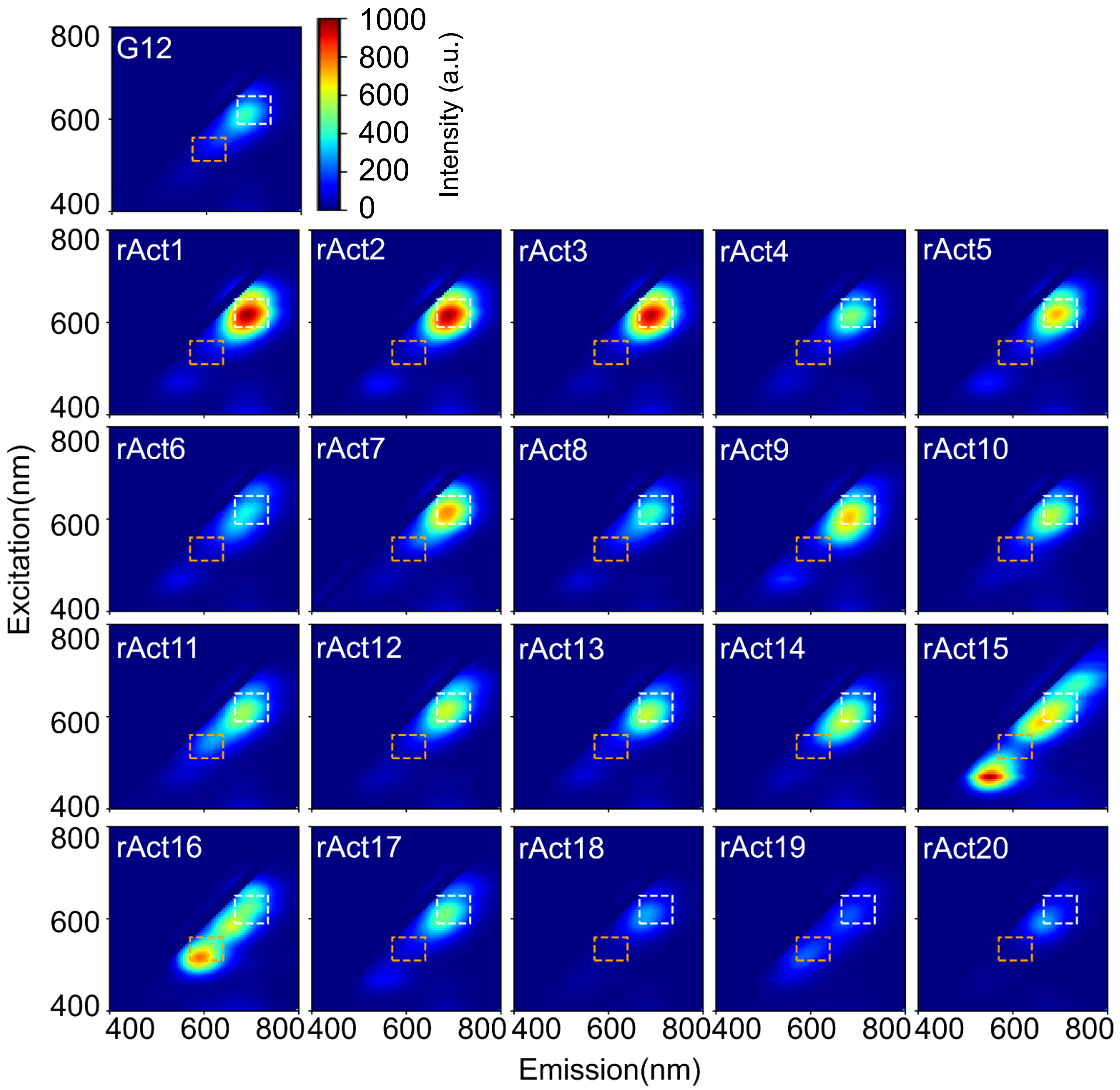


**Figure S22 | 2D spectra of bright red activator candidates (including the 3 false positive selections: rACT18, rACT19 and rACT20)** Compared to G12 NCB (ATCCGGGGTGGGGTGGGG), 17 out of 20 bright red activator candidates (selected by the chip screening method) have the improvement ratio greater than one (85% accuracy). In particular, rAct1 NCB (TCCATTGGTGGGGTGGGG) has the improvement ratio of 2.94. The white dashed box represents the integrated region of red channel (Ex/Em: 620/60, 700/75 nm), and the orange dashed box represents the integrated region of yellow channel (Ex/Em: 535/50, 605/70 nm). See **Table S2a** for details.

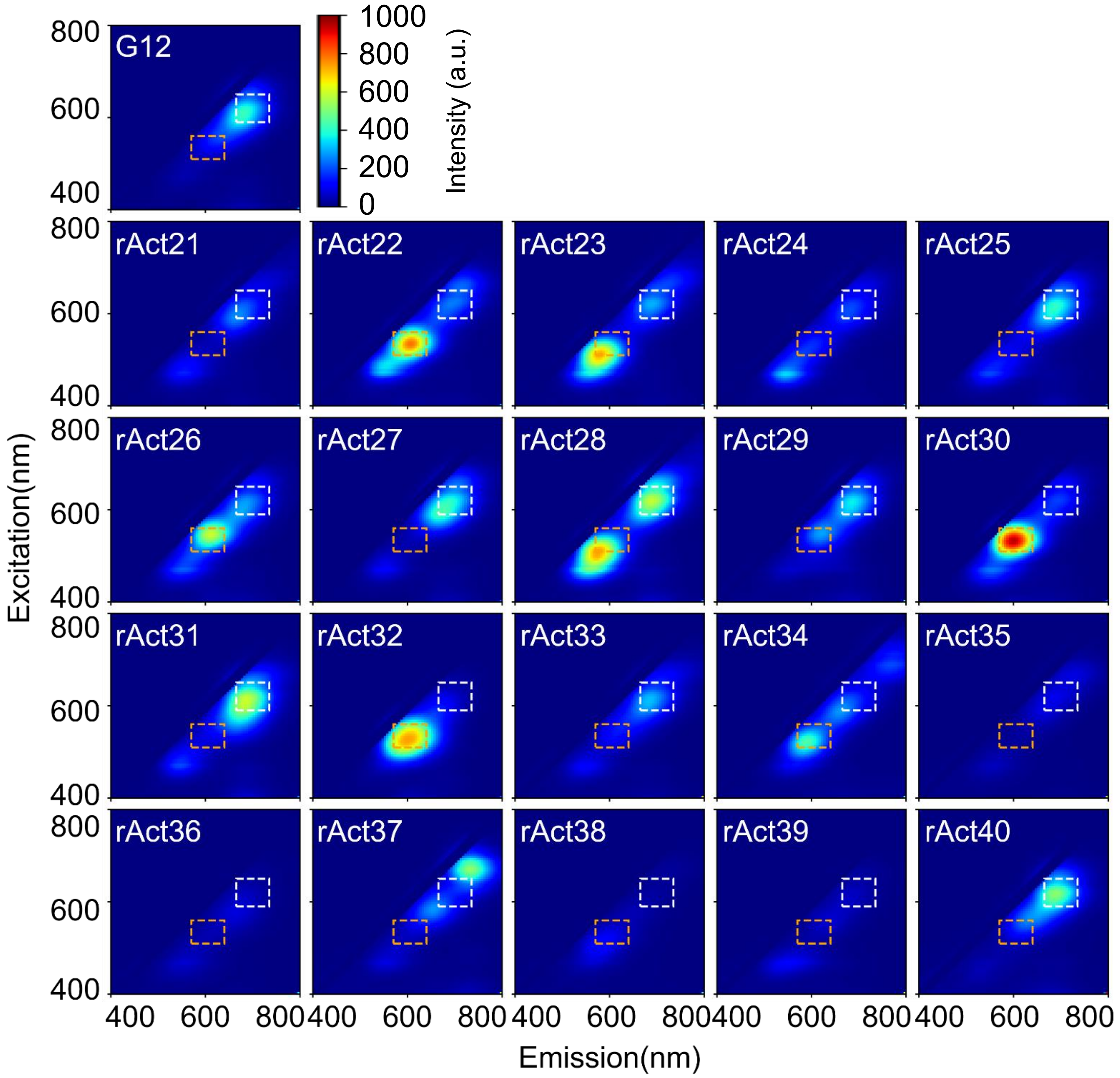


**Figure S23 | 2D spectra of dark activator candidates (including 3 false negative selections: rACT28, rACT31 and rACT40)** Compared to G12 NCB (ATCCGGGGTGGGGTGGGG), 17 out of 20 dark candidates (selected by the chip screening method) have the improvement ratio less than one (85% accuracy). In particular, rAct38 NCB (GGGTGGGTTTATGTGGGG) has the improvement ratio of 0.10. The white dashed box represents the integrated region of red channel (Ex/Em: 620/60, 700/75 nm), and the orange dashed box represents the integrated region of yellow channel (Ex/Em: 535/50, 605/70 nm). See **Table S2b** for details.

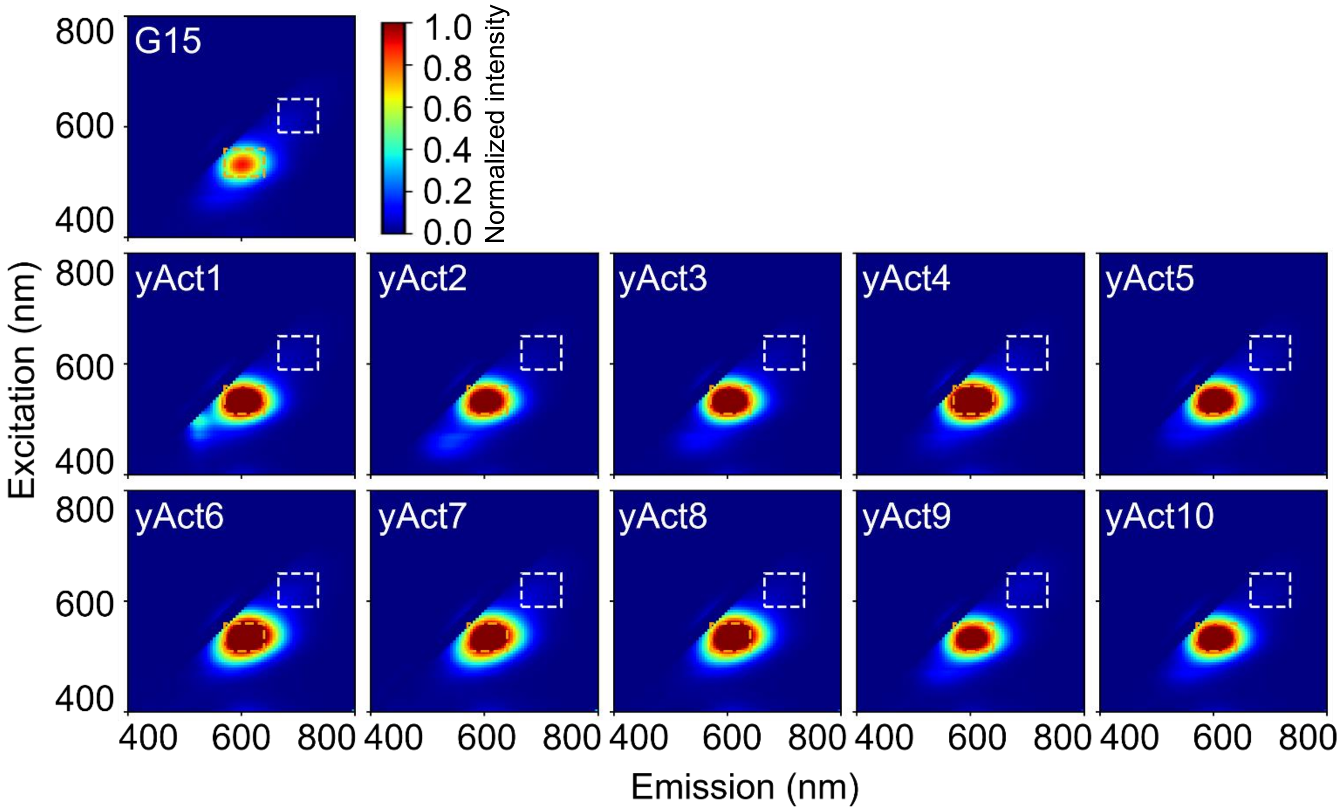


**Figure S24 | 2D spectra of bright yellow activator candidates.** Compared to G15 NCB (GGGTGGGGTGGGGTGGGG), all 10 bright yellow activator candidates (selected by the chip screening method) have the improvement ratio greater than one (100% accuracy). In particular, yAct4 NCB (TTGGTGGGTGGGGTGGGG) has the improvement ratio of 2.03. Fluorescence intensity normalized to G15 peak intensity. The white dashed box represents the integrated region of red channel (Ex/Em: 620/60, 700/75 nm), and the orange dashed box represents the integrated region of yellow channel (Ex/Em: 535/50, 605/70 nm). See **Table S3** for details.

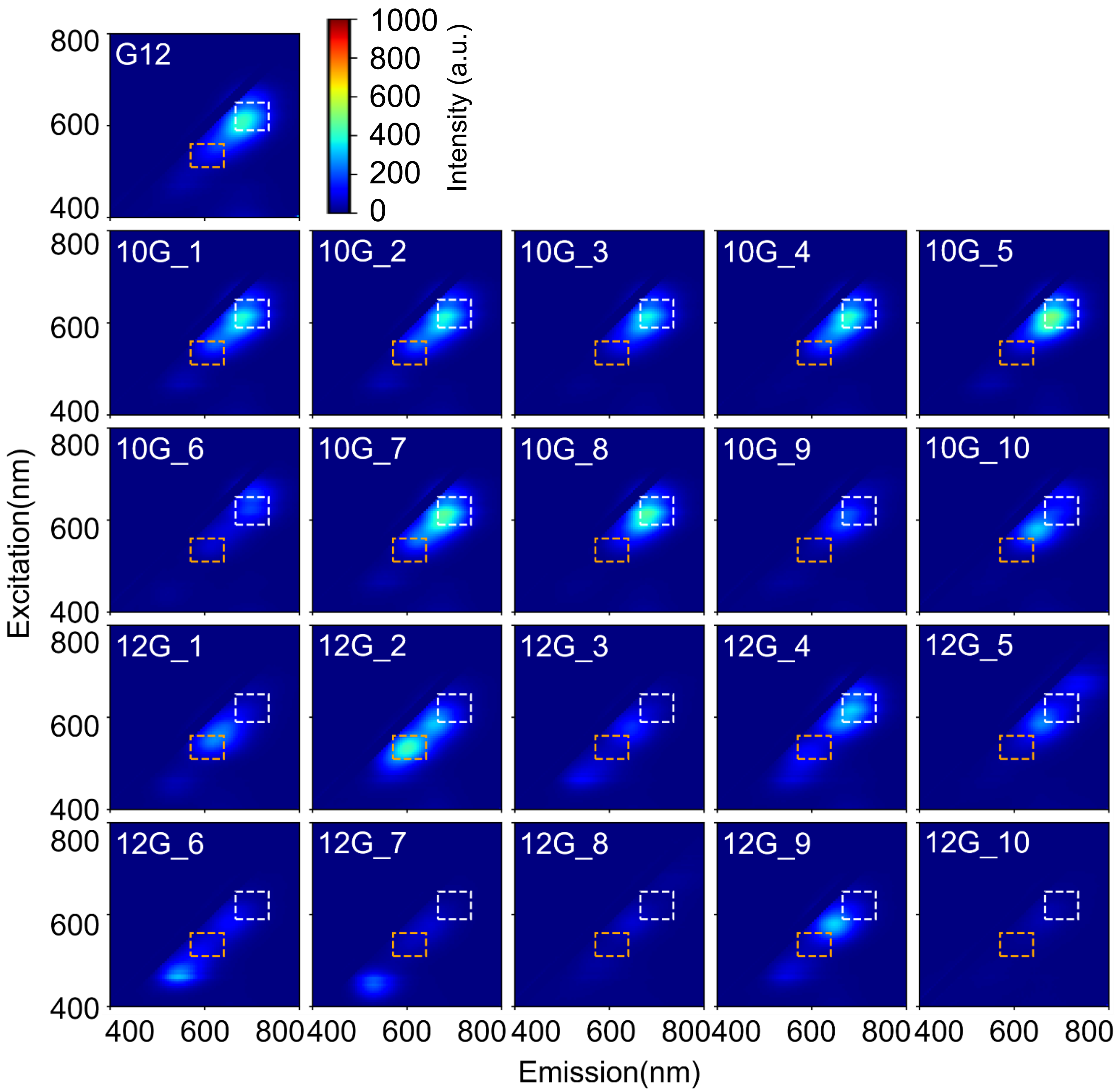


**Figure S25 | 2D spectra of activators with various numbers of guanine bases.** Based on chip selection results, ten 10G activators can potentially be brighter than G12 NCB (**Table S4**) and ten 12G activators can potentially be darker than G12 NCB (**Table S5**). Test-tube investigation proves that 7 of the selected 10G activators have their enhancement ratios comparable to that of G12 in the red channel (improvement ratio ≥ 0.9). and all selected 12G activators are darker than G12 in the red channel (improvement ratio < 0.6). This result indicates that it is possible to create bright red NCBs with fewer numbers of guanine. The white dashed box represents the integrated region of red channel (Ex/Em: 620/60, 700/75 nm), and the orange dashed box represents the integrated region of yellow channel (Ex/Em: 535/50, 605/70 nm).

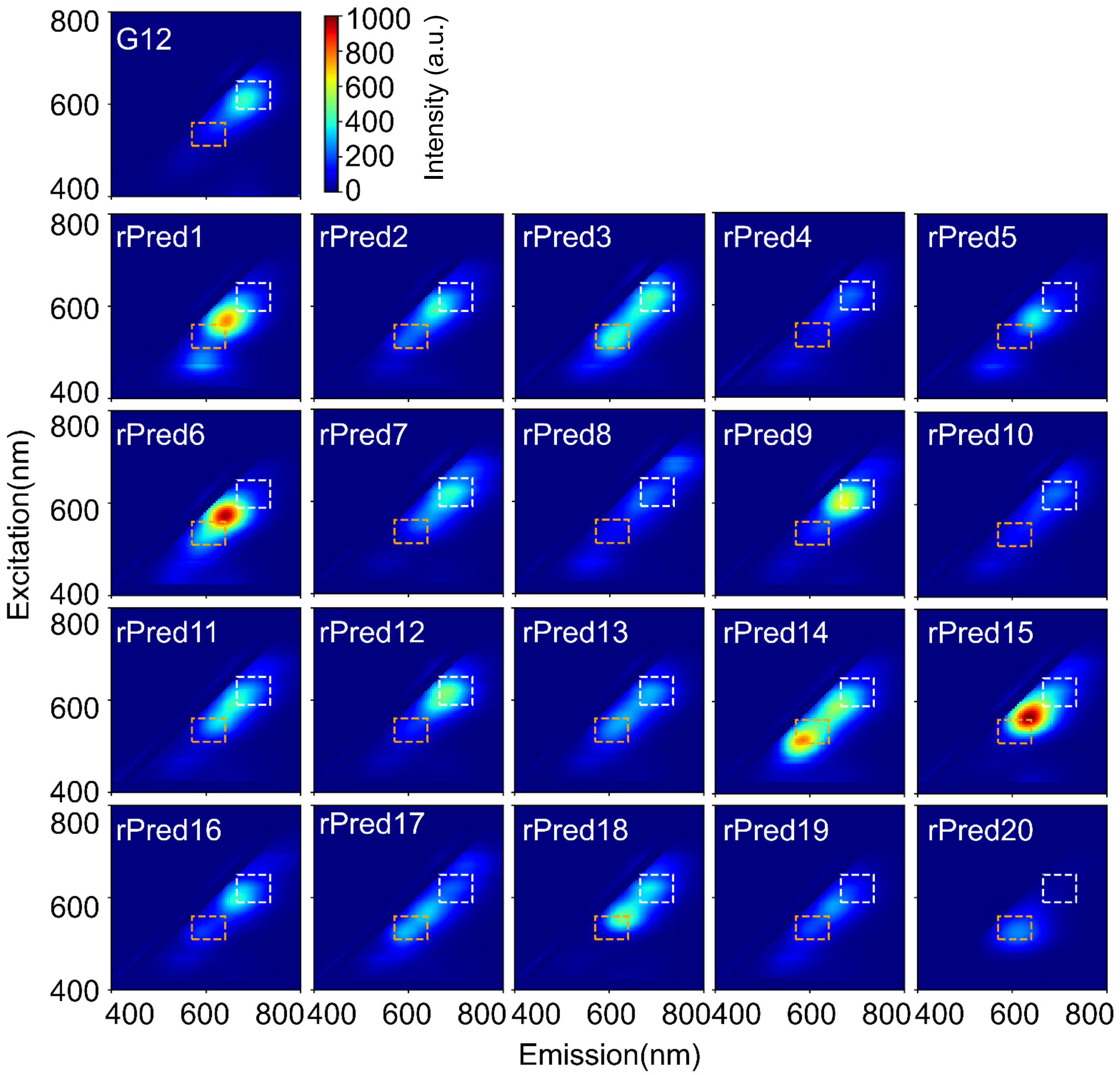


**Figure S26 | 2D spectra of rationally designed red NCBs.** Twenty activators are designed based on the machine learning results and evaluated in test tubes. On average, the enhancement ratio was 291 for these twenty designs. The white dashed box represents the integrated region of red channel (Ex/Em: 620/60, 700/75 nm), and the orange dashed box represents the integrated region of yellow channel (Ex/Em: 535/50, 605/70 nm).

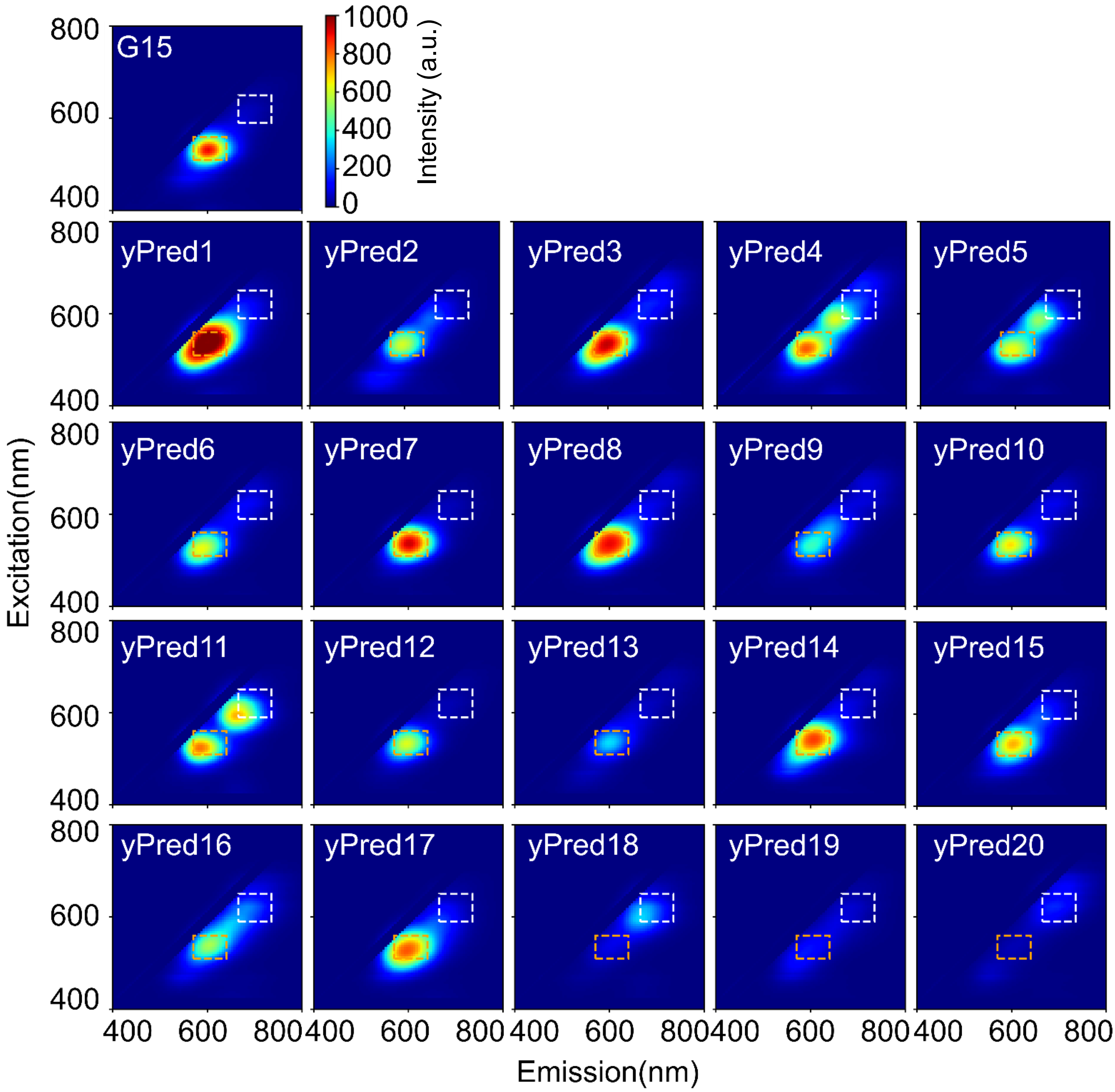


**Figure S27 | 2D spectra of rationally designed yellow NCBs.** Twenty activators are designed based on the machine learning results and evaluated in test tubes. On average, the enhancement ratio was 532 for these twenty. The white dashed box represents the integrated region of red channel (Ex/Em: 620/60, 700/75 nm), and the orange dashed box represents the integrated region of yellow channel (Ex/Em: 535/50, 605/70 nm).

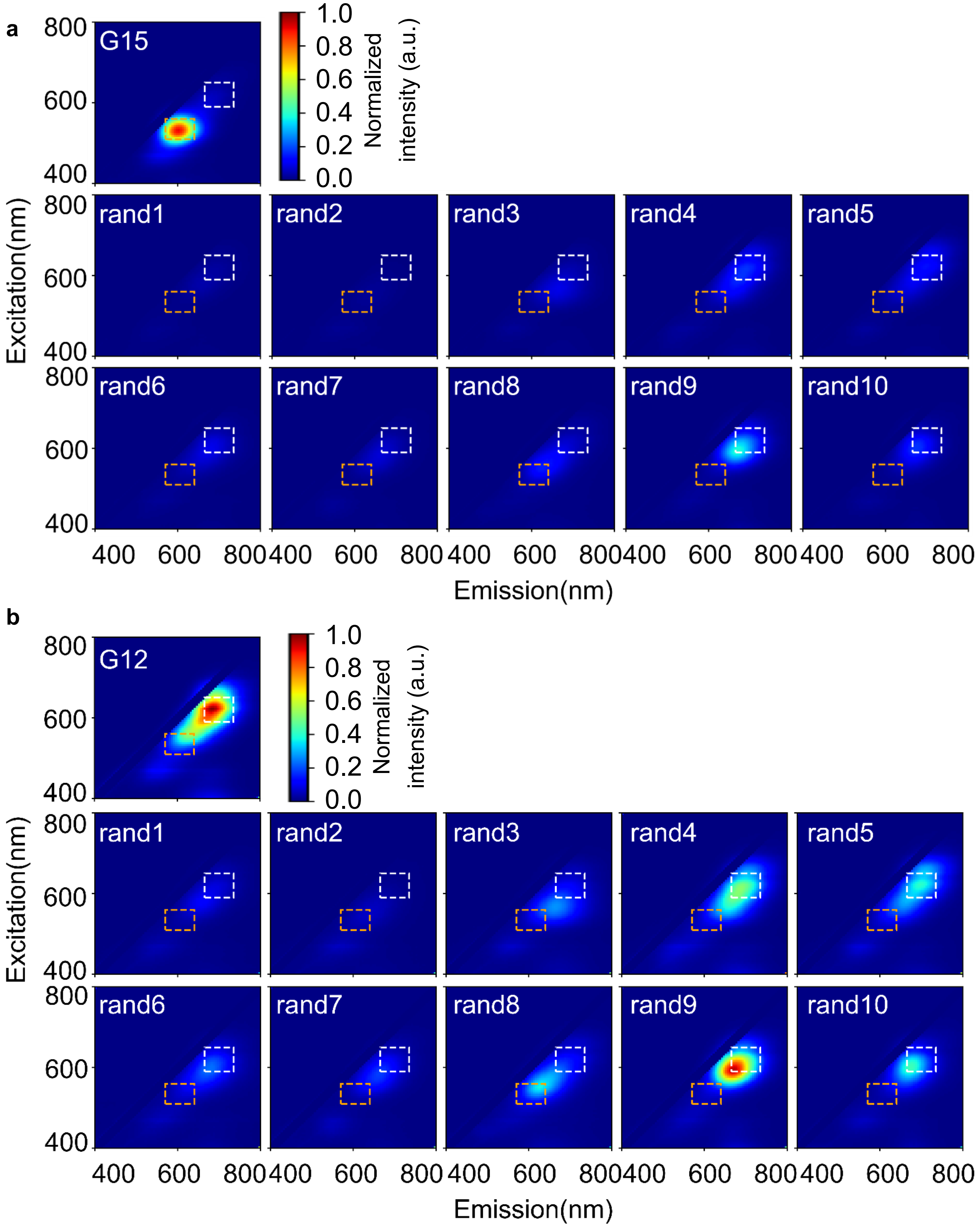


**Figure S28 | 2D spectra of randomly designed NCBs.** Ten activators are randomly designed and evaluated in test tubes. The enhancement ratio of rand8 passed the threshold in the yellow channel, while rand4, rand5, rand9 and rand10 passed the threshold in the red channel (**Table S8**). Fluorescence intensity normalized to G15 peak intensity (**a**) and G12 peak intensity (**b**). The white dashed box represents the integrated region of red channel (Ex/Em: 620/60, 700/75 nm), and the orange dashed box represents the integrated region of yellow channel (Ex/Em: 535/50, 605/70 nm).

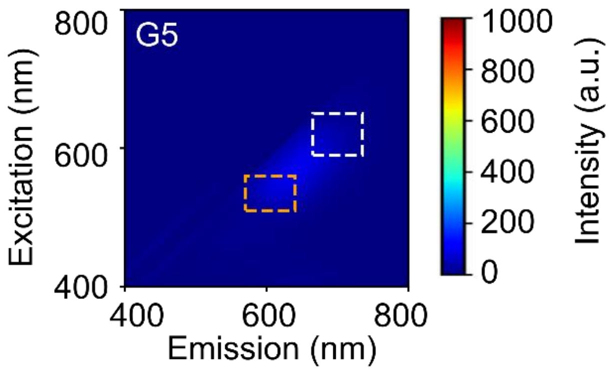


**Figure S29 | 2D spectra of G5 NCBs.** Based on the design rules discussed in **Fig. 2,** we speculated that this G5 activator (CCCCCCGCGGGTTTCCC) would lead to a bright NCB. However, the result was actually a low red enhancement ratio (39, as compared to 439 for G12). This result clearly indicated that segments do not work alone – cooperativities among the segments determine the activation color and intensity of an NCB.

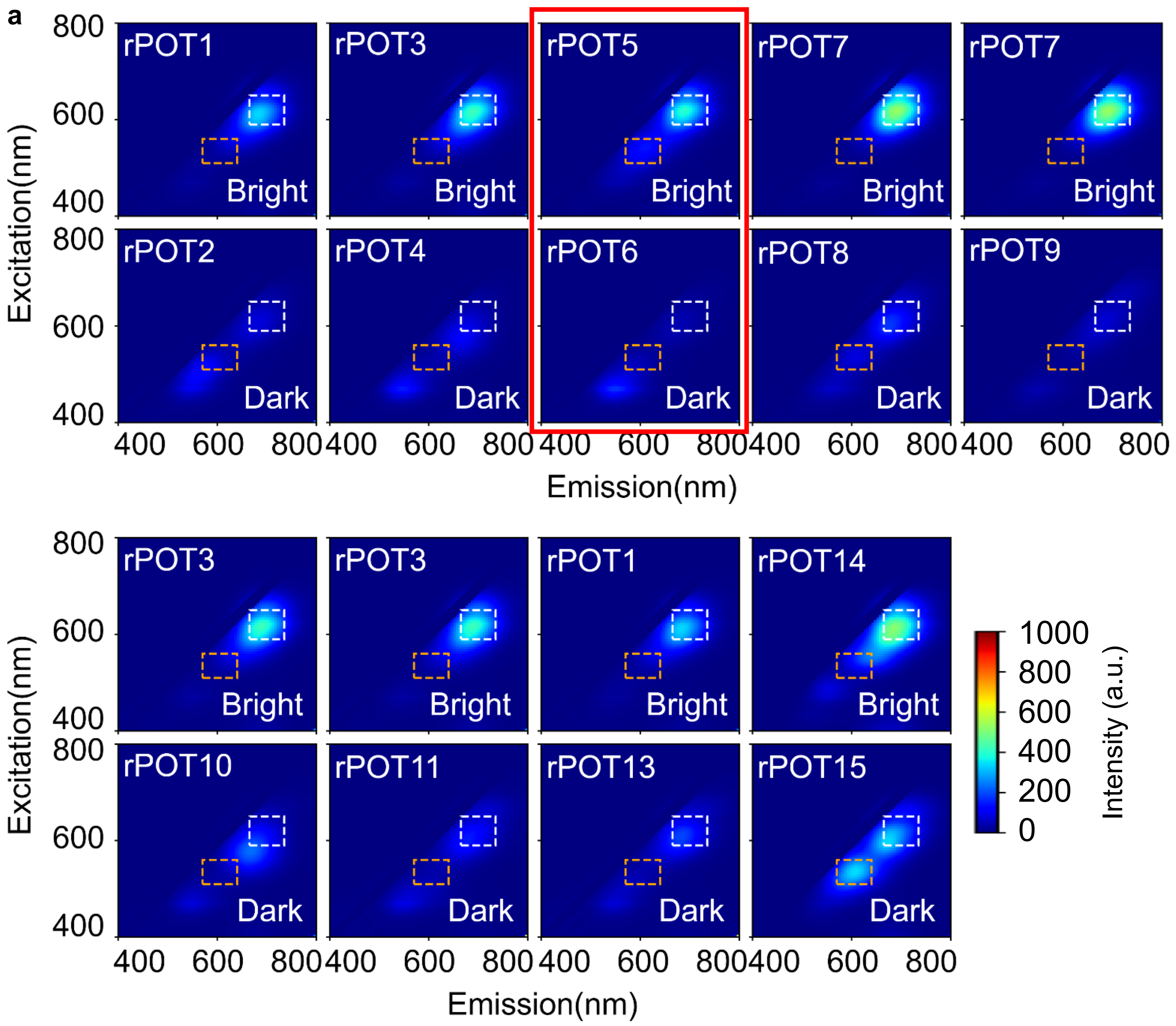


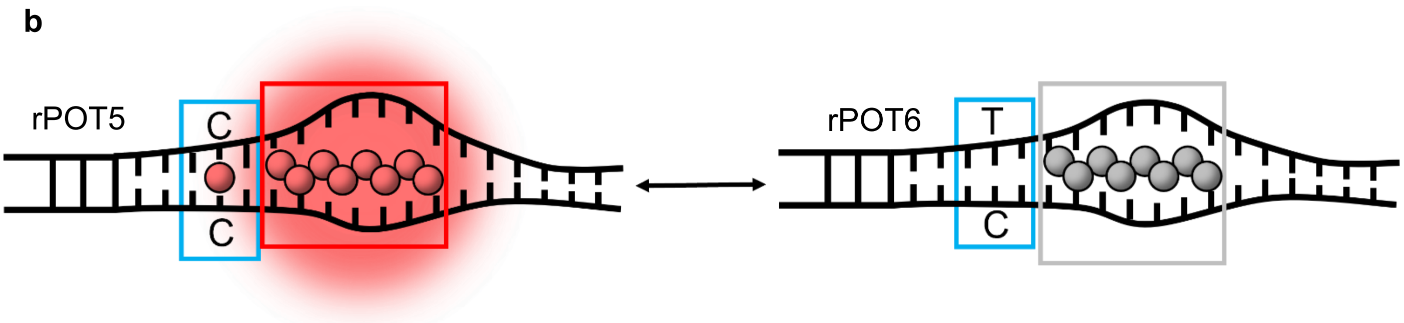


**Figure S30 | 2D spectra of red POT candidates. a** Based on chip selection results, 9 sets of red POT candidates are evaluated in test tubes. All these candidates have their POT difference ratios greater than 1.7, with the largest difference ratio of 9.12 (rPOT5/rPOT6 NCBs, highlighted in solid red box, **Table S8**). **b** Based on the hotspots from **Fig. 3**, we hypothesized that the disruption of silver-mediated pair C-$Ag^+$-C would darken red NCB samples and form red POT pairs. The white dashed box represents the integrated region of red channel (Ex/Em: 620/60, 700/75 nm), and the orange dashed box represents the integrated region of yellow channel (Ex/Em: 535/50, 605/70 nm). The blue box represents the hotspots of red POTs. The red and gray boxes represent the bag position of bright member of red POTs and its counterpart, respectively.

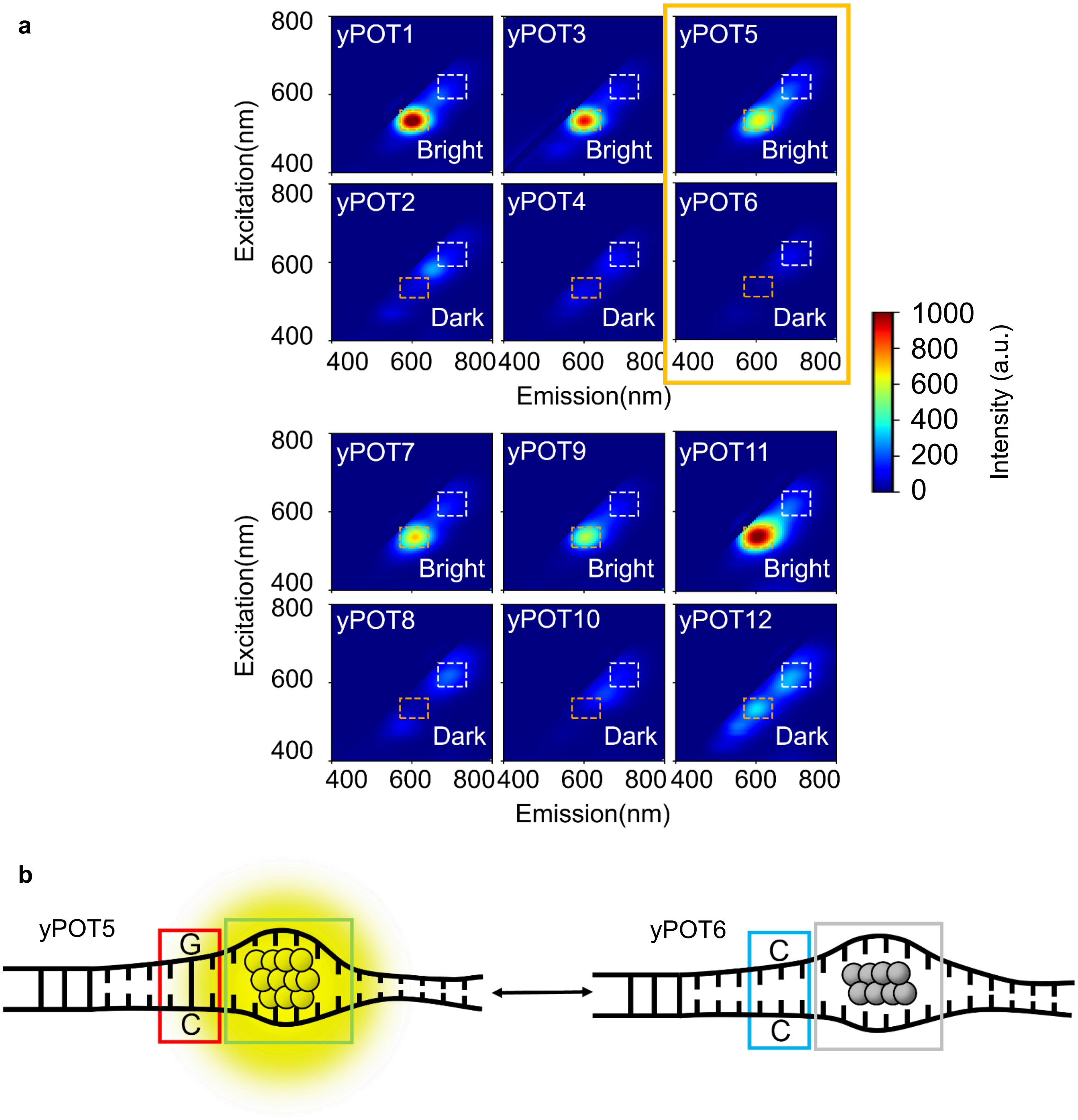


**Figure S31 | 2D spectra of yellow POT candidates. a** Based on chip selection results, 9 sets of yellow POT candidates are evaluated in test tubes. All these candidates have their POT difference ratios greater than 3.0, with the largest difference ratio of 37.04 (yPOT5/yPOT6 NCBs, highlighted in solid orange box, **Table S9**). **b** Based on the hotspots from **Fig. 3**, we hypothesized that the disruption of WC pair, GC pairing, would darken yellow NCB samples and form yellow POT pairs. The white dashed box represents the integrated region of red channel (Ex/Em: 620/60, 700/75 nm), and the orange dashed box represents the integrated region of yellow channel (Ex/Em: 535/50, 605/70 nm). The blue box represents the hotspots of yellow POTs. The gold and gray boxes represent the bag position of bright member of yellow POTs and its counterpart, respectively.

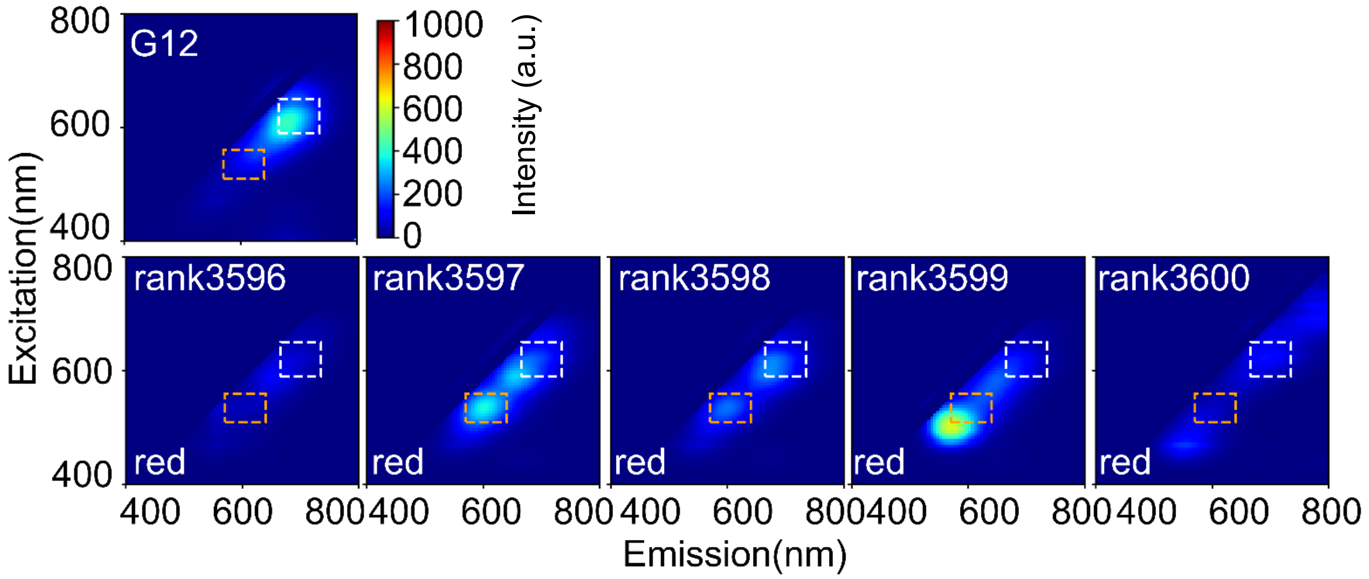


**Figure S32 | 2D spectra of red NCBs near ranking 3,600.** As we apply top 3,600 sequences as our bright class to perform ML modeling. We evaluate the fluorescence intensity of the NCBs ranking near 3,600. The median enhancement ratio 145 was set as our threshold to evaluate the rationally designed red NCBs.

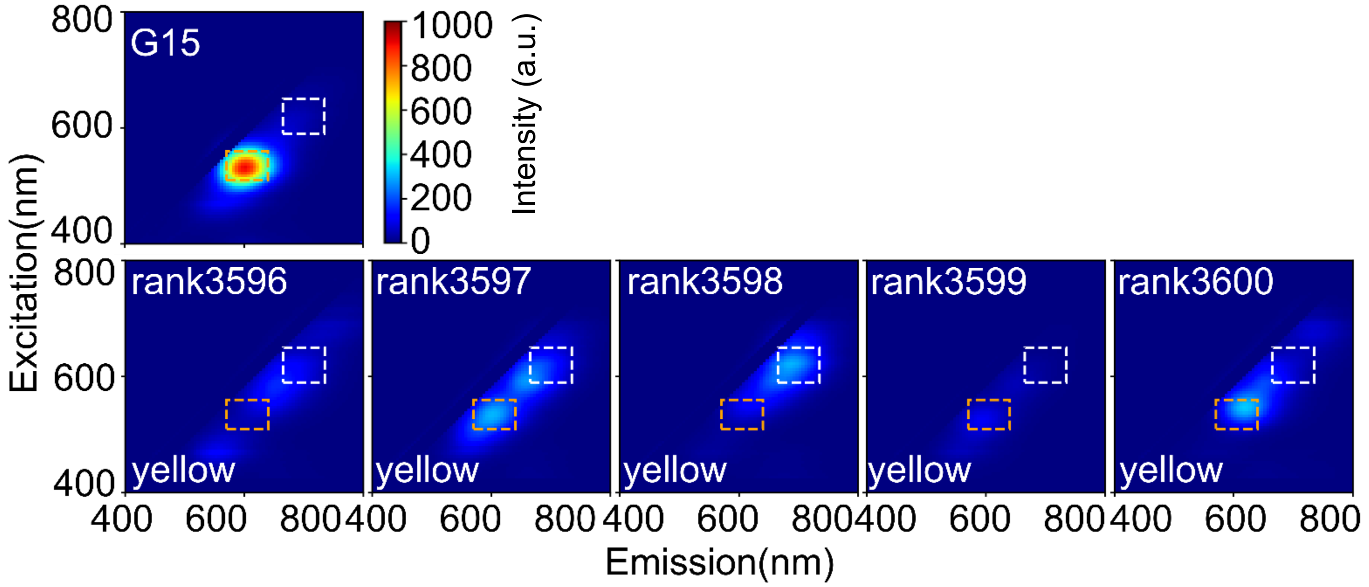


**Figure S33 | 2D spectra of yellow NCBs near ranking 3,600.** As we apply top 3,600 sequences as our bright class to perform ML modeling. We evaluate the fluorescence intensity of the NCBs ranking near 3,600. The median enhancement ratio 66 was set as our threshold to evaluate the rationally designed yellow NCBs.

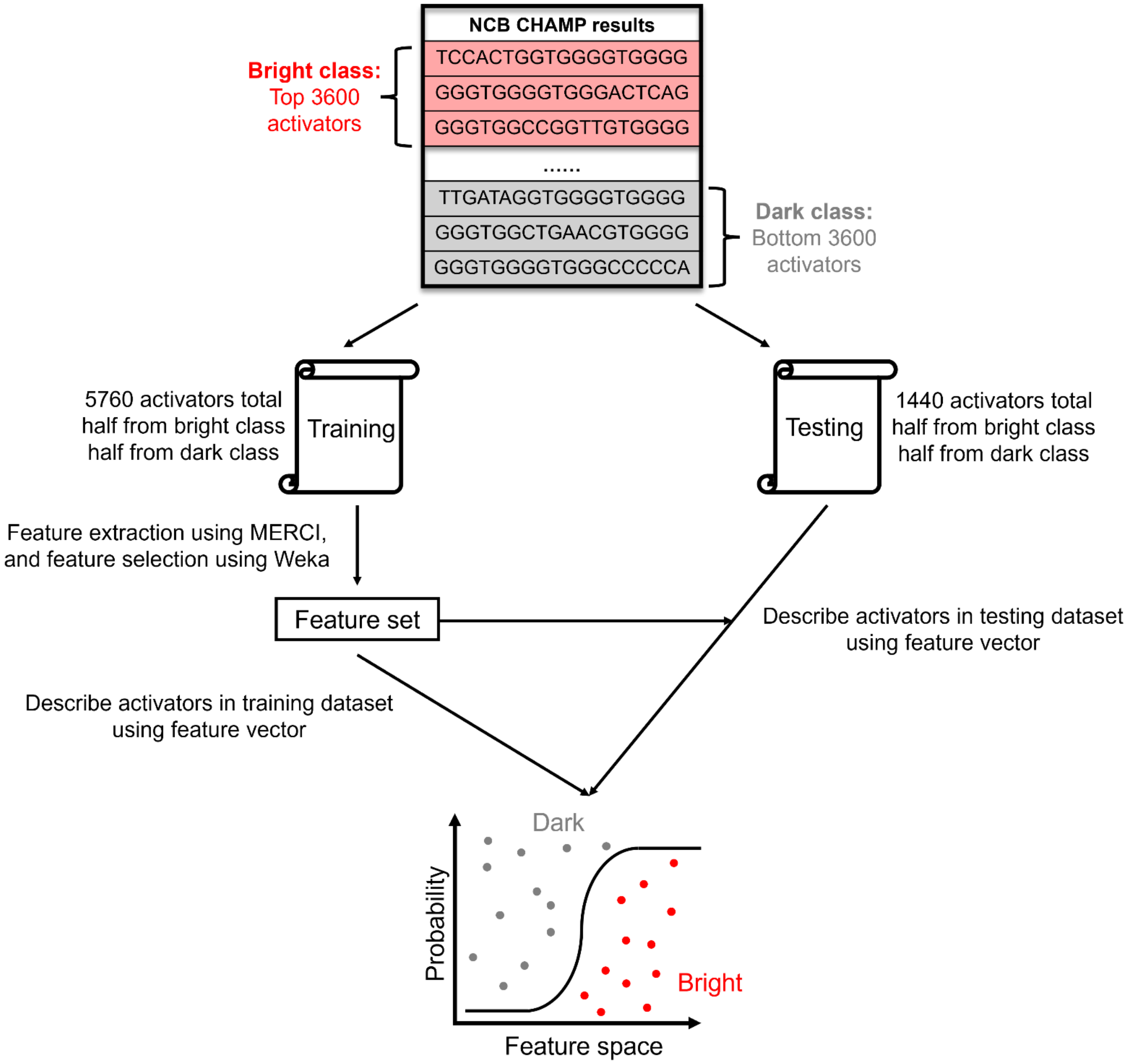


**Figure S34 | Workflow for establishing machine learning models to classify screened NCBs or NCB candidates.** In this report, we performed 5-fold CV to classify our library sequences. Following the approaches proposed by Copp and Gwinn[11-13], we labeled the top 30% NCBs as "bright" class and the bottom 30% as "dark" class. The feature extraction process was proceeded using MERCI[14]. The extracted motifs were processed with Python scripts to include the position information. We then identified the most discriminative set of features using Weka.[15] Based on the selected features, a number of models were established for classifying the chip screening results and we found the model built on LR has the best performance.

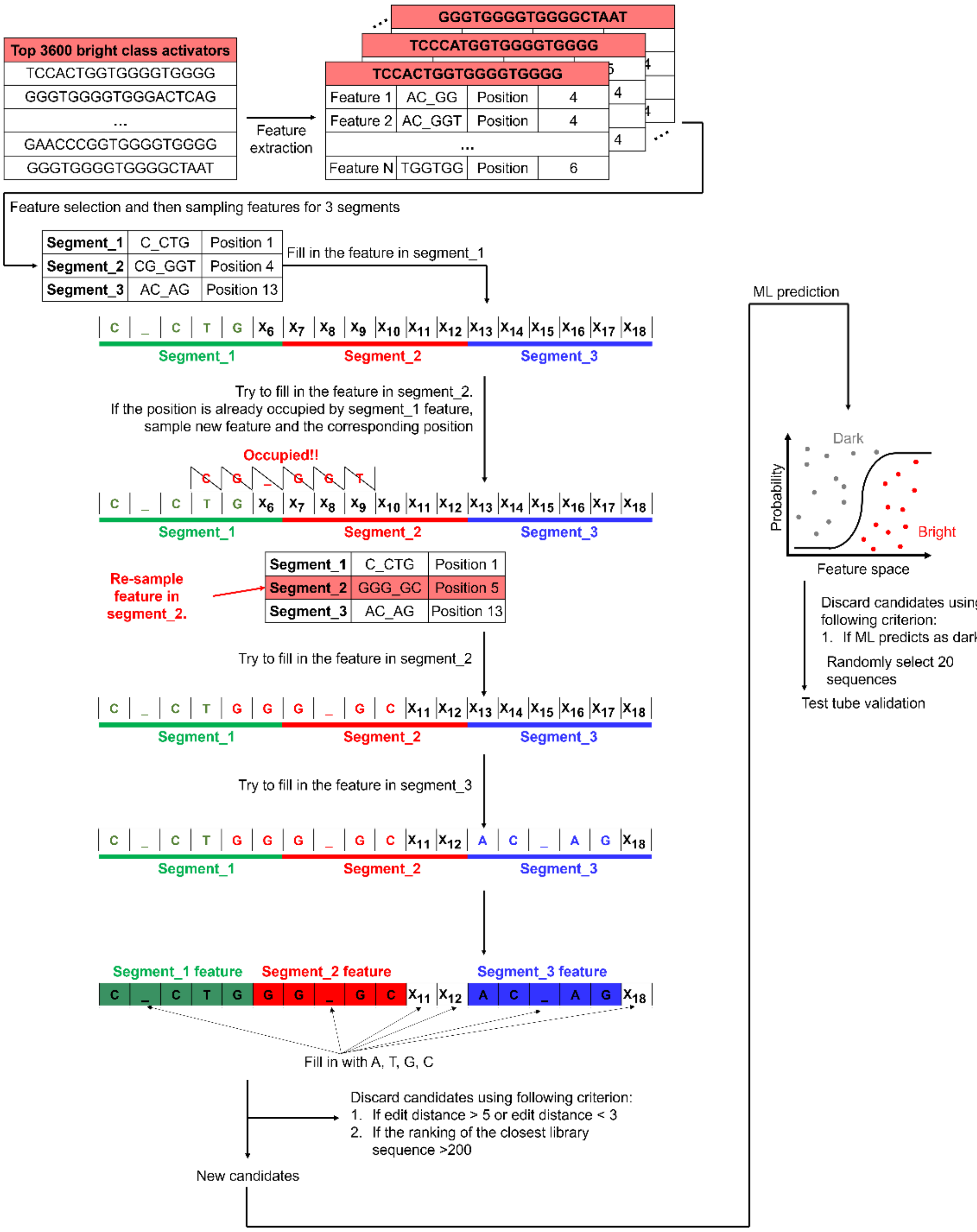


**Figure S35 | Workflow to rationally design bright NCBs.** Based on the most discriminative features identified by Weka, we sampled the distribution of these features in each segment and generated a list of common motifs with their corresponding positions. To construct a red NCB candidate, we assigned 3 features to the blank 18-nt template, starting with feature_1 insertion into segment_1. As feature_2 might have an overlap with feature_1 when being inserted into segment_2. In that situation, the design algorithm would replace feature_2 with another feature to ensure no overlap. However, if any two features shared identical bases at their overlapping site, they were considered as “compatible” and could be inserted into the same template. For example, as shown above, feature C_CTG (positions 1-5) and feature GGG_GC (positions 5-10) shared a guanine base at the overlapping site (position 5). Consequently, they were compatible and were used in constructing a bright NCB candidate. The same procedure was repeated until a compatible feature for segment_3 was found. Once all three features were inserted into the template, the remaining blank positions were filled up based on the composition popularity (at the same positions) from the bright class sequences. The edit distance[13,16] of the new candidate was then assessed. We only selected new candidates with edit distance between 3 to 5 from the top 200 bright activators screened on chip for test-tube investigation (**Table S6**-**S7**).

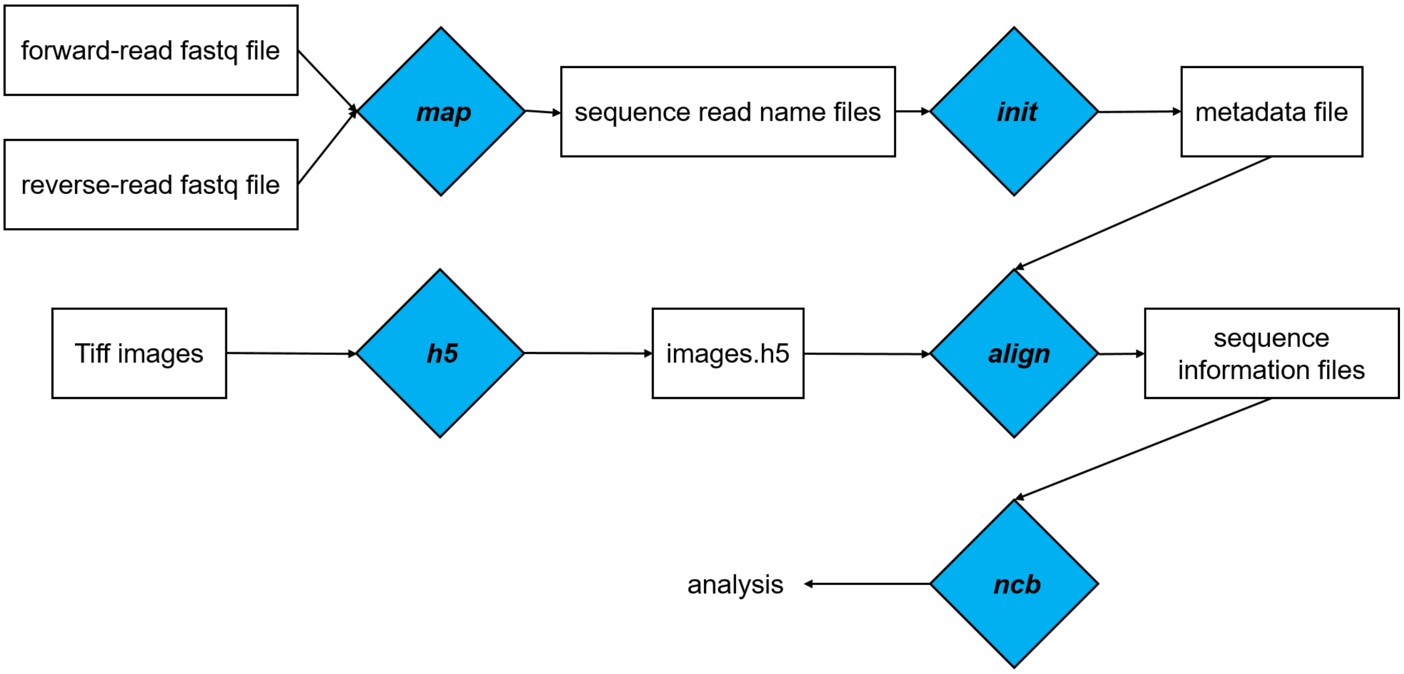


**Figure S36 | CHAMP workflow.** A custom bioinformatics and imaging processing pipeline named CHAMP (Chip-Hybridized Associated Mapping Platform) was developed by Finkelstein's group and the detailed algorithm description can be found in ref. 6. CHAMP helped decipher the activator sequence behind each activated NCB spot (termed the NCB-CHAMP selection method, **Fig. 1c** and **Supplementary Fig. S4-S5**). In brief, mapping the alignment markers was done at four stages. First, a rough alignment was carried out using Fourier-based cross correlation, followed by a precision alignment using least-squares constellation mapping between FASTQ and *de novo* extracted NCB spots. We built up the consensus sequences and their corresponding information (e.g., lane number, tile number, and x-y coordinates) at all reported positions in the FASTQ file using the ***map*** command. Second, the ***init*** command was executed to record the metadata of imaging settings (e.g., rotation and scaling). Third, the ***h5*** command was applied to generate a single hdf5 file containing all 512×512 PhiX fiducial marker images. Fourth, the ***align*** command transformed the processed sequence information into pseudo-images and performed precise alignment. The output files were saved individually by image positions. The content included x, y coordinates of each sequence and the corresponding sequence ID. To analyze our NCB images, we developed an additional function named ***ncb***, which corrected the uneven illumination using flat-field correction. A bootstrap method was then performed to derive the median intensity of each activator in order to rank the NCB brightness (**Supplementary Fig. S36**).

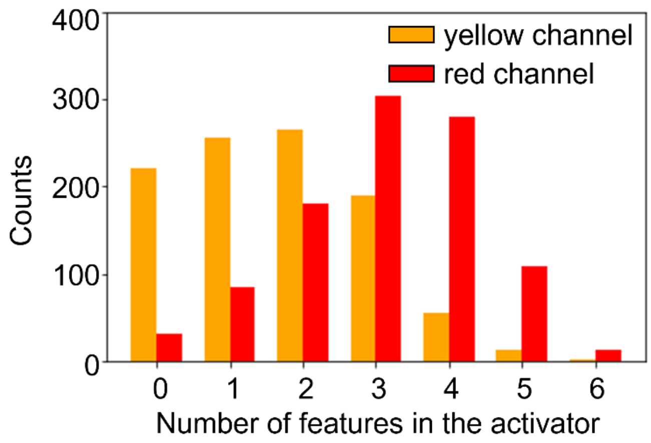


**Figure S37 | Feature distribution for the top 1,000 library sequences for red and yellow channels.** By evaluating the selected bright features within the top 1,000 library sequences for red and yellow channels, we found the optimal number of features to create bright NCBs would be 2 and 3 for yellow and red channels, respectively.

**Table S1: Sequences of probes and library designs used in this report.**

**a.** RE strand is used for restriction enzyme digestion (MauBI). The three Atto probes are used for digestion evaluation and NCB-CHAMP alignment. **b.** The 6-segment and 9-segment interrogation of library_1. **c.** Three different library designs. For each of our library sequences on *MiSeq* chip, it consists of six parts: P5 (light blue), SP1 (gold), library sequence (gray for hybridization segment, purple for activator, and dark blue for restriction site), SP2 (orange), barcode (red), and P7 (green). P5 and P7 are adapters for surface attachment. SP1 and SP2 are sequencing-by-synthesis primer binding sites. Barcodes are reserved and used by *Illumina*. The 30-nt-long hybridization segment is for C55 hybridization and the 18-nt-long activator part is where we call "the library". As for the library size, library_1 contains 12,286 sequences, library_2 contains 12,286 sequences, and library_3 contains 16,255 sequences. The '**CG**' represents the remaining nucleotides after cleavage by a restriction enzyme, and the vertical line represents the cutting site.

**a**

| **Acronym** | **Sequence (5' → 3')** |
|---|---|
| C55 | CCC CCT TAA TCC CCC TAT AAT AAA TTT TAA ATA TTA TTT ATT AAT |
| G15 | ATT AAT AAA TAA TAT TTA AAA TTT ATT ATA GGG TGG GGT GGG GTG GGG |
| G12 | ATT AAT AAA TAA TAT TTA AAA TTT ATT ATA ATC CGG GGT GGG GTG GGG |
| RE strand | CAG ACG TGT GCT CTT CCG ATC TCG CGC GCG NN |
| Atto647N-tagged comp_MauBI | /5ATTO647N/ CAG ACG TGT GCT CTT CCG ATC TCG CGC GCG NN |
| Atto647N-tagged AT | /5ATTO647N/ CCC CCT TAA TCC CCC TAT AAT AAA TTT TAA ATA TTA TTT ATT AAT |
| Atto488_cPhiX | /5Alex488N/CG GTC TCG GCA TTC CTG CTG AAC CGC TCT TCC GAT C |
| Forward primer | AAT GAT ACG GCG ACC ACC GAG A |
| Reverse primer | CAA GCA GAA GAC GGC ATA CGA GA |
| G15 (90 nt) | ATT AAT AAA TAA TAT TTA AAA TTT ATT ATA GGG TGG GGT GGG GTG GGG AGA TCG GAA GAG CAC ACG TCT GAA CTC CAG TCA CTT GTT CAT |

**b 6-segment interrogation and 9-segment interrogation in library_1**

| **Acronym** | **Sequence (5' → 3')** |
|---|---|
| **The 6-segment interrogation** | |
| Segment_11 | ATT AAT AAA TAA TAT TTA AAA TTT ATT ATA **NNN** TGG GGT GGG GTG GGG |
| Segment_12 | ATT AAT AAA TAA TAT TTA AAA TTT ATT ATA GGG **NNN** GGT GGG GTG GGG |
| Segment_21 | ATT AAT AAA TAA TAT TTA AAA TTT ATT ATA GGG TGG **NNN** GGG GTG GGG |
| Segment_22 | ATT AAT AAA TAA TAT TTA AAA TTT ATT ATA GGG TGG GGT **NNN** GTG GGG |
| Segment_31 | ATT AAT AAA TAA TAT TTA AAA TTT ATT ATA GGG TGG GGT GGG **NNN** GGG |
| Segment_32 | ATT AAT AAA TAA TAT TTA AAA TTT ATT ATA GGG TGG GGT GGG GTG **NNN** |

| **The 9-segment interrogation** | |
|---|---|
| Segment_11 | ATT AAT AAA TAA TAT TTA AAA TTT ATT ATA NNG TGG GGT GGG GTG GGG |
| Segment_12 | ATT AAT AAA TAA TAT TTA AAA TTT ATT ATA GGN NGG GGT GGG GTG GGG |
| Segment_13 | ATT AAT AAA TAA TAT TTA AAA TTT ATT ATA GGG TNN GGT GGG GTG GGG |
| Segment_21 | ATT AAT AAA TAA TAT TTA AAA TTT ATT ATA GGG TGG NNT GGG GTG GGG |
| Segment_22 | ATT AAT AAA TAA TAT TTA AAA TTT ATT ATA GGG TGG GGN NGG GTG GGG |
| Segment_23 | ATT AAT AAA TAA TAT TTA AAA TTT ATT ATA GGG TGG GGT GNN GTG GGG |
| Segment_31 | ATT AAT AAA TAA TAT TTA AAA TTT ATT ATA GGG TGG GGT GGG NNG GGG |
| Segment_32 | ATT AAT AAA TAA TAT TTA AAA TTT ATT ATA GGG TGG GGT GGG GTN NGG |
| Segment_11 | ATT AAT AAA TAA TAT TTA AAA TTT ATT ATA GGG TGG GGT GGG GTG GNN |

**c sequence information of library_1, library_2 and library_3**

| **Acronym** | **Sequence (5' → 3')** |
|---|---|
| Canonical activator G15 in library_1 | AATGATACGGCGACCACCGAGA (P5)<br>TCTACACTCTTTCCCTACACGACGCTCTTCCGATCT (SP1)<br>**ATTAATAAATAATATTTAAAATTTATTATAGGGTGGGGTGGGGTGGGG (activator) CG \| CGCGCG (restriction site)**<br>AGATCGGAAGAGCACACGTCTGAACTCCAGTCAC(SP2) *GTAGAG* (barcode) ATCTCGTATGCCGTCTTCTGCTTG (P7) |
| Canonical activator G15 in library_2 and library_3 | AATGATACGGCGACCACCGAGA (P5)<br>TCTACACTCTTTCCCTACACGACGCTCTTCCGATCT (SP1)<br>**ATTAATAAATAATATTTAAAATTTATTATAGGGTGGGGTGGGGTGGGG (activator)**<br>AGATCGGAAGAGCACACGTCTGAACTCCAGTCAC(SP2) *GTAGAG* (barcode) ATCTCGTATGCCGTCTTCTGCTTG (P7) |
| Segment_1 activators In library_1 | AATGATACGGCGACCACCGAGA (P5)<br>TCTACACTCTTTCCCTACACGACGCTCTTCCGATCT (SP1)<br>**ATTAATAAATAATATTTAAAATTTATTATANNNNNNGGTGGGGTGGGG (activator) CG \| CGCGCG (restriction site)**<br>AGATCGGAAGAGCACACGTCTGAACTCCAGTCAC(SP2) *TTGTTC*(barcode) ATCTCGTATGCCGTCTTCTGCTTG (P7) |
| Segment_2 activators in library_1 | AATGATACGGCGACCACCGAGA (P5)<br>TCTACACTCTTTCCCTACACGACGCTCTTCCGATCT (SP1)<br>**ATTAATAAATAATATTTAAAATTTATTATAGGGTGGNNNNNNGTGGGG (activator) CG \| CGCGCG (restriction site)**<br>AGATCGGAAGAGCACACGTCTGAACTCCAGTCAC(SP2) *TTGTTC*(barcode ATCTCGTATGCCGTCTTCTGCTTG (P7) |
| Segment_3 activators In library_1 | AATGATACGGCGACCACCGAGA (P5)<br>TCTACACTCTTTCCCTACACGACGCTCTTCCGATCT (SP1)<br>**ATTAATAAATAATATTTAAAATTTATTATAGGGTGGGGTGGGNNNNNN (activator) CG \| CGCGCG (restriction site)**<br>AGATCGGAAGAGCACACGTCTGAACTCCAGTCAC(SP2) *TTGTTC*(barcode) ATCTCGTATGCCGTCTTCTGCTTG (P7) |

| | |
|---|---|
| Segment_1 activators In library_2 | AATGATACGGCGACCACCGAGA (P5)<br>TCTACACTCTTTCCCTACACGACGCTCTTCCGATCT (SP1)<br>**ATTAATAAATAATATTTAAAATTTATTATA<u>GGGNNNNNNGGGTGGGG</u>**<br>**(activator)** AGATCGGAAGAGCACACGTCTGAACTCCAGTCAC(SP2)<br>*TTGTTC* (barcode) ATCTCGTATGCCGTCTTCTGCTTG (P7) |
| Segment_2 activators In library_2 | AATGATACGGCGACCACCGAGA (P5)<br>TCTACACTCTTTCCCTACACGACGCTCTTCCGATCT (SP1)<br>**ATTAATAAATAATATTTAAAATTTATTATA<u>GGGTGGGGTNNNNNNGGG</u>**<br>**(activator)** AGATCGGAAGAGCACACGTCTGAACTCCAGTCAC(SP2)<br>*TTGTTC*(barcode) ATCTCGTATGCCGTCTTCTGCTTG (P7) |
| Segment_3 activators In library_2 | AATGATACGGCGACCACCGAGA (P5)<br>TCTACACTCTTTCCCTACACGACGCTCTTCCGATCT (SP1)<br>**ATTAATAAATAATATTTAAAATTTATTATA<u>NNNTGGGGTGGGGTGNNN</u>**<br>**(activator)** AGATCGGAAGAGCACACGTCTGAACTCCAGTCAC(SP2)<br>*TTGTTC*(barcode) ATCTCGTATGCCGTCTTCTGCTTG (P7) |
| Segment_1 activators In library_3 | AATGATACGGCGACCACCGAGA (P5)<br>TCTACACTCTTTCCCTACACGACGCTCTTCCGATCT (SP1)<br>**ATTAATAAATAATATTTAAAATTTATTATA<u>NNNTGGNNNGGGGTGGGG</u>**<br>**(activator)** AGATCGGAAGAGCACACGTCTGAACTCCAGTCAC(SP2)<br>*TTGTTC*(barcode) ATCTCGTATGCCGTCTTCTGCTTG (P7) |
| Segment_2 activators in library_3 | AATGATACGGCGACCACCGAGA (P5)<br>TCTACACTCTTTCCCTACACGACGCTCTTCCGATCT (SP1)<br>**ATTAATAAATAATATTTAAAATTTATTATA<u>GGGNNNGGTNNNGTGGGG</u>**<br>**(activator)** AGATCGGAAGAGCACACGTCTGAACTCCAGTCAC(SP2)<br>*TTGTTC*(barcode) ATCTCGTATGCCGTCTTCTGCTTG (P7) |
| Segment_3 activators library_3 | AATGATACGGCGACCACCGAGA (P5)<br>TCTACACTCTTTCCCTACACGACGCTCTTCCGATCT (SP1)<br>**ATTAATAAATAATATTTAAAATTTATTATA<u>GGGTGGNNNGGGNNNGGG</u>**<br>**(activator)** AGATCGGAAGAGCACACGTCTGAACTCCAGTCAC(SP2)<br>*TTGTTC*(barcode) ATCTCGTATGCCGTCTTCTGCTTG (P7) |
| Segment_4 activators In library_3 | AATGATACGGCGACCACCGAGA (P5)<br>TCTACACTCTTTCCCTACACGACGCTCTTCCGATCT (SP1)<br>**ATTAATAAATAATATTTAAAATTTATTATA<u>GGGTGGGGTNNNGTGNNN</u>**<br>**(activator)** AGATCGGAAGAGCACACGTCTGAACTCCAGTCAC(SP2)<br>*TTGTTC*(barcode) ATCTCGTATGCCGTCTTCTGCTTG (P7) |

**Table S2: Test-tube investigation of selected bright red and dark activator candidates.**

To validate our NCB-CHAMP selection method, twenty top-ranked and twenty bottom-ranked activators are further investigated in test tubes. **a.** Using G12 NCB (ATCCGGGGTGGGGTGGGG) as the standard for red emitter comparison, 17 out 20 bright red candidates are found brighter than G12 NCB in test tubes (also see **Fig. S13**). **b.** 17 out 20 dark candidates are found darker than G12 NCB in test tubes (**Fig. S14**). The formulas to compute "enhancement ratio" and "improvement ratio" are described in ***Methods***. In short, we first calculate the volumetric integrated intensity (**Fig. S2**) from the 2D spectrum of the sample in the red channel (Ex: 620/60 nm, Em: 700/75 nm). From there we calculate the enhancement ratio:

$$\text{Enhancement ratio}=(I_{NCB}-I_{NC\ probe})/(I_{NC\ probe}-I_{background})$$

where $I_{NCB}$ stands for the volumetric integrated intensity of NCB in red of yellow window, $I_{NC\ probe}$ represents the volumetric integrated intensity of dark AgNC on C55 probe, and $I_{background}$ is the volumetric integrated intensity of buffer (i.e., sodium phosphate buffer). The improvement ratio is simply the ratio of the enhancement ratio of an NCB to that of the standard red NCB (G12). False selections are highlighted in gray below.

**a Selected red bright NCB candidates:**

| ID | Activator (5' → 3') | Enhancement ratio in red channel | Improvement ratio (compared to G12) |
|---|---|---|---|
| G12 | ATCCGGGGTGGGGTGGGG | 439 | 1 |
| rAct1 | TCCATTGGTGGGGTGGGG | 1292 | 2.94 |
| rAct2 | TCCAATGGTGGGGTGGGG | 1247 | 2.84 |
| rAct3 | TCCATAGGTGGGGTGGGG | 1229 | 2.80 |
| rAct4 | ATCCGTGGTGGGGTGGGG | 601 | 1.37 |
| rAct5 | TCCTATGGTGGGGTGGGG | 756 | 1.72 |
| rAct6 | TCTCATGGTGGGGTGGGG | 498 | 1.13 |
| rAct7 | ATCCCAGGTGGGGTGGGG | 973 | 2.22 |
| rAct8 | CCTTCTGGTGGGGTGGGG | 532 | 1.21 |
| rAct9 | CACATTGGTGGGGTGGGG | 764 | 1.74 |
| rAct10 | CCCCAAGGTGGGGTGGGG | 669 | 1.52 |
| rAct11 | CCTTGAGGTGGGGTGGGG | 575 | 1.31 |
| rAct12 | TCACTAGGTGGGGTGGGG | 616 | 1.40 |
| rAct13 | ACTCGTGGTGGGGTGGGG | 583 | 1.33 |
| rAct14 | CCTGCAGGTGGGGTGGGG | 678 | 1.54 |
| rAct15 | GGGTGGGGTGGGGCTAGA | 833 | 1.90 |
| rAct16 | TGGGACGGTGGGGTGGGG | 694 | 1.58 |
| rAct17 | TGAACAGGTGGGGTGGGG | 516 | 1.18 |
| rAct18 | GCTACAGGTGGGGTGGGG | 269 | 0.61 |
| rAct19 | CGGTTTGGTGGGGTGGGG | 166 | 0.38 |
| rAct20 | CGCTTCGGTGGGGTGGGG | 201 | 0.46 |

**b Selected dark candidates:**

| ID | Activator (5' → 3') | Enhancement ratio in red channel | improvement ratio (compared to G12) |
|---|---|---|---|
| G12 | ATCCGGGGTGGGGTGGGG | 439 | 1 |
| rAct21 | GGGTGGGGTGGGACGCTA | 221 | 0.51 |
| rAct22 | ATCTGAGGTGGGGTGGGG | 236 | 0.54 |
| rAct23 | GGGTGGGGTGGGGACATT | 256 | 0.58 |
| rAct24 | AAGTTTGGTGGGGTGGGG | 139 | 0.32 |
| rAct25 | AACGATGGTGGGGTGGGG | 404 | 0.92 |
| rAct26 | TGGCTTGGTGGGGTGGGG | 272 | 0.62 |
| rAct27 | CTGCTTGGTGGGGTGGGG | 423 | 0.96 |
| rAct28 | GGGTGGGGTGGGGAGATC | 587 | 1.34 |
| rAct29 | CTGGCCGGTGGGGTGGGG | 347 | 0.79 |
| rAct30 | AAAAGGGGTGGGGTGGGG | 192 | 0.44 |
| rAct31 | ACGTTTGGTGGGGTGGGG | 575 | 1.30 |
| rAct32 | GGTGCAGGTGGGGTGGGG | 124 | 0.28 |
| rAct33 | ACAATAGGTGGGGTGGGG | 375 | 0.86 |
| rAct34 | GGGTGGGGTGTGGTGGGG | 243 | 0.55 |
| rAct35 | GGGTGGCGATTAGTGGGG | 148 | 0.35 |
| rAct36 | GGGTGGTAATGTGTGGGG | 78 | 0.19 |
| rAct37 | GGGTGGGGTGGGTGTAGG | 310 | 0.71 |
| rAct38 | GGGTGGGTTTATGTGGGG | 44 | 0.10 |
| rAct39 | GGGTGGTCAAAAGTGGGG | 81 | 0.19 |
| rAct40 | GGGTGGCCTCCAGTGGGG | 571 | 1.29 |

**Table S3: Test-tube investigation of selected bright yellow activator candidates.**

To validate our NCB-CHAMP selection method, ten top-ranked yellow activators are further investigated in test tubes. **A.** Using G15 NCB (GGGTGGGGTGGGGTGGGG) as the standard for yellow emitter comparison, all 10 bright yellow candidates are found brighter than G15 NCB in test tubes (also see **Fig. S15**). The formulas to compute "enhancement ratio" and "improvement ratio" are described in ***Methods***. In short, we first calculate the volumetric integrated intensity (**Fig. S2**) from the 2D spectrum of the sample in the yellow channel (Ex: 535/50 nm, Em: 605/70 nm). From there we calculate the enhancement ratio:

$$\text{Enhancement ratio}=(I_{NCB}-I_{NC\ probe})/(I_{NC\ probe}-I_{background})$$

where $I_{NCB}$ stands for the volumetric integrated intensity of NCB in yellow window, $I_{NC\ probe}$ represents the volumetric integrated intensity of dark AgNC on C55 probe, and $I_{background}$ is the volumetric integrated intensity of buffer (i.e., sodium phosphate buffer). The improvement ratio is simply the ratio of the enhancement ratio of an NCB to that of the standard yellow activator (G15).

| ID | Sequence (5' → 3') | Enhancement ratio | Improvement ratio (compared to G15) |
|---|---|---|---|
| G15 | GGGTGGGGTGGGGTGGGG | 553 | 1 |
| yAct1 | CAGGTGGGTGGGGTGGGG | 988 | 1.79 |
| yAct2 | TTTGTGGGTGGGGTGGGG | 872 | 1.58 |
| yAct3 | TGTGTGGGTGGGGTGGGG | 951 | 1.72 |
| yAct4 | TTGGTGGGTGGGGTGGGG | 1125 | 2.03 |
| yAct5 | AAGTTGGGTGGGGTGGGG | 924 | 1.67 |
| yAct6 | AGTTGAGGTGGGGTGGGG | 1105 | 2.00 |
| yAct7 | TTGTGAGGTGGGGTGGGG | 1009 | 1.82 |
| yAct8 | GTTTGAGGTGGGGTGGGG | 1045 | 1.89 |
| yAct9 | AGTTTGGGTGGGGTGGGG | 840 | 1.52 |
| yAct10 | ATGTTGGGTGGGGTGGGG | 922 | 1.67 |

**Table S4: Test-tube investigation of 10-guanine activators**

Based on chip selection results, ten 10G activators can potentially be brighter than G12 NCB (**Fig. S15**). Test-tube investigation proves that 7 of the selected 10G activators have their enhancement ratios comparable to that of G12 in the red channel (improvement ratio ≥ 0.9). This result indicates that it is possible to create bright red NCBs with fewer numbers of guanine. False selections are highlighted in gray below.

| **ID** | **Sequence (5' → 3')** | **Enhancement ratio** | **Improvement ratio (compared to G12)** |
|---|---|---|---|
| G12 | ATCCGGGGTGGGGTGGGG | 439 | 1 |
| 10G_1 | AACCTTGGTGGGGTGGGG | 415 | 0.95 |
| 10G_2 | TCCAATGGTGGGGTGGGG | 409 | 0.93 |
| 10G_3 | ATCCATGGTGGGGTGGGG | 397 | 0.91 |
| 10G_4 | ATCCCAGGTGGGGTGGGG | 413 | 0.95 |
| 10G_5 | TACCATGGTGGGGTGGGG | 527 | 1.21 |
| 10G_6 | GGGTGGTCCCCCGTGGGG | 240 | 0.54 |
| 10G_7 | AACCATGGTGGGGTGGGG | 480 | 1.09 |
| 10G_8 | CTCCATGGTGGGGTGGGG | 473 | 1.09 |
| 10G_9 | ACATCAGGTGGGGTGGGG | 204 | 0.47 |
| 10G_10 | GGGTGGCCCCCCGTGGGG | 221 | 0.51 |

**Table S5: Test-tube investigation for activators having at least 12 guanines**

Based on chip selection results, ten 12G activators can potentially be darker than G12 NCB (**Fig. S15**). Test-tube investigation proves that all of the selected 12G activators are darker than G12 in the red channel (improvement ratio < 0.6).

| ID | Sequence (5' → 3') | Enhancement ratio | Improvement ratio (compared to G12) |
|---|---|---|---|
| G12 | ATCCGGGGTGGGGTGGGG | 439 | 1 |
| 12G_1 | GGGTGGTCGGACGTGGGG | 61 | 0.15 |
| 12G_2 | GGGTGGTGTCAGGTGGGG | 124 | 0.29 |
| 12G_3 | GGGTGGAAGAGGGTGGGG | 51 | 0.12 |
| 12G_4 | GGGTGGTTGCTGGTGGGG | 260 | 0.59 |
| 12G_5 | GGGTGGGTCGCCGTGGGG | 126 | 0.29 |
| 12G_6 | GGGTGGAGTGATGTGGGG | 45 | 0.11 |
| 12G_7 | GGGTGGTGAGACGTGGGG | 26 | 0.06 |
| 12G_8 | GGGTGGGCTGACGTGGGG | 31 | 0.08 |
| 12G_9 | GGGTGGAAGAGTGTGGGG | 57 | 0.14 |
| 12G_10 | GGGTGGACGACGGTGGGG | 17 | 0.05 |

**Table S6: Test-tube investigation of rationally designed bright red NCBs.**

Twenty activators are designed based on the machine learning results and evaluated in test tubes. Following the observation shown in **Supplementary Fig. S37**, the new red candidates were generated if three bright red features were presented in the sequences as shown below. On average, the enhancement ratio was 291 for these twenty and the mean edit distance was 4.0. Here the enhancement ratio of 145 was set as the cutoff for bright yellow NCBs (**Table S9a**). Three out of the 20 rationally designed red NCBs below showed either low emission (rPred19 and rPred20, highlighted in gray) or yellow emission (rPred14, highlighted in yellow). Thus, the overall test-tube validation accuracy was 85%. Among the 20 NCBs below, rPred9 NCB was the brightest (1.30-fold brighter than G12 NCB, highlighted in red).

| ID | Activator (5' → 3') | Emission peak (nm) | Minimal edit distance | Enhancement ratio | Improvement ratio compared to G12 | Motif #1 | Motif #2 | Motif #3 |
|---|---|---|---|---|---|---|---|---|
| rPred1 | TCCCATGCGGGGCTCGGG | 655 | 4 | 220 | 0.50 | CCC | GGG_C | TC_GG |
| rPred2 | CCCGAAGGGGGGATCGCG | 640 | 4 | 250 | 0.57 | CCC | GGGGA | TC_CG |
| rPred3 | CCCGAAGGTGGGGCTCTG | 655 | 4 | 487 | 1.11 | CCC | GA_GGT | C_CTG |
| rPred4 | TACCAAGGGGGGAACGGG | 685 | 4 | 167 | 0.38 | CA_GG | GGGGA | AA_GG |
| rPred5 | ACCAGAGGGGTGGGCCCG | 645 | 5 | 154 | 0.35 | ACC_G | AG_GGT | CCC |
| rPred6 | TCCCAAGGTGGGGGGCAG | 640 | 3 | 198 | 0.45 | CCC | CA_GG | GGGG_C |
| rPred7 | TCCCGAGGTTGGGTCTGG | 685 | 3 | 408 | 0.93 | CCC | CG_GGT | CTGG |
| rPred8 | TCCAGCGGGGGAGGGGGC | 735 | 4 | 184 | 0.42 | TCC_G | GGGGA | GGG_C |
| rPred9 | ATCCCTCGGGGAGGGGGC | 670 | 5 | 571 | 1.30 | CCC | GGGGA | GGGG_C |
| rPred10 | CATCCGTTGGGGGACGGG | 685 | 5 | 180 | 0.41 | A_CCG | TTGG_G | GGGAC |
| rPred11 | GCCCGAGGGGGGGACGGG | 655 | 3 | 373 | 0.85 | CCC | C_AGG | GGGAC |
| rPred12 | TCCAGTGGGGGGAGCGGG | 680 | 4 | 505 | 1.15 | TCC_G | GGGGA | GCGG |
| rPred13 | CCCGTAGGGTAGGTTGGG | 685 | 4 | 316 | 0.72 | CCC | TA_GGT | TT_GG |
| rPred14 | CCCGAAGGGGGGGGCATG | 580 | 5 | 531 | 1.21 | CCC | AA_GG | GGG_C |
| rPred15 | TCCCGCGGGGGGGACGGG | 635 | 3 | 325 | 0.74 | CCC | GGGA | GGGAC |
| rPred16 | TCCGGACGGGGGTGGGGG | 660 | 3 | 277 | 0.63 | TCC_G | AC_GG | TGGGGG |
| rPred17 | TCCCCAGGGGGACTGGGG | 640 | 3 | 170 | 0.39 | CCC | GGGGA | CTGG |
| rPred18 | ATCCTTCGGGGGATCGGG | 630 | 5 | 380 | 0.87 | CTT_G | GGGGA | A_CGG |
| rPred19 | TCCAAGGGGTGGACTGGC | 650 | 4 | 101 | 0.23 | AA_GG | AG_GGT | CTGG |
| rPred20 | TACCCAGGGGGACTGGGC | 650 | 4 | 31 | 0.07 | CCC | CA_GG | ACT_G |

**Table S7: Test-tube investigation of rationally designed yellow NCBs.**

Twenty activators are designed based on the machine learning results and evaluated in test tubes. Following the observation shown in **Supplementary Fig. S37**, the new yellow candidates were generated if two bright yellow features were presented in the sequences as shown below. On average, the enhancement ratio was 532 for these twenty designs and the mean edit distance was 3.5. Here the enhancement ratio of 66 was set as the cutoff for bright yellow NCBs (**Table S9b**). Three out of the 20 rationally designed yellow NCBs below showed either low emission (yPred18, yPred19 and rPred20, highlighted in gray) or red emission (yPred18 and rPred20). Thus, the overall test-tube validation accuracy was 85%. Among the 20 NCBs below, yPred1 NCB was the brightest (2.30-fold brighter than G15 NCB, highlighted in red).

| ID | Activator (5' → 3') | Emission peak (nm) | Minimal edit distance | Enhancement ratio | Improvement ratio compared to G15 | Motif #1 | Motif #2 |
|---|---|---|---|---|---|---|---|
| yPred1 | GTGTTGGGTGGTCGGGGG | 585 | 3 | 1272 | 2.30 | GTG_TG | TGGGTG |
| yPred2 | GGTGTGGGTGGGAAGGGC | 595 | 3 | 371 | 0.67 | GT_TG | TGGGTG |
| yPred3 | TGTGTGTGGGGGATGGGG | 595 | 3 | 968 | 1.75 | GT_TGG | GGGGG |
| yPred4 | GCTGTGTGGGGTGTGGGG | 585 | 3 | 724 | 1.31 | GT_TGG | GTGTGG |
| yPred5 | GGAGTGGGTGGTGGTGGG | 590 | 3 | 487 | 0.88 | TGGGTG | GTG_TG |
| yPred6 | TCGGTGTGGTGTGTGGGG | 585 | 4 | 299 | 0.54 | GTGGTG | GTGTGG |
| yPred7 | TGGTGTGGTTGGCGGGGG | 600 | 3 | 946 | 1.71 | GT_TGG | T_GCG |
| yPred8 | AGTGTGGTGTTGGGGGGG | 595 | 5 | 619 | 1.12 | GTGGTG | TTG_GG |
| yPred9 | GCTTGGGTGGGTGTGGGC | 600 | 3 | 448 | 0.81 | GT_TGG | TGGGTG |
| yPred10 | AGTGGGTGTGTGTGGGGG | 595 | 4 | 680 | 1.23 | GT_TGG | TGGGTG |
| yPred11 | GAGTTAGGGGTGTGGGGC | 580 | 5 | 885 | 1.60 | GT_AG | GT_TGG |
| yPred12 | AGTGTGGGTGTGTGGGGG | 595 | 3 | 481 | 0.87 | GT_TGG | TGGGTG |
| yPred13 | GGTGTGGGTGTGTGGGGG | 600 | 3 | 249 | 0.45 | GT_TGG | TGGGTG |
| yPred14 | GTTGTGGTGGGAGGGGGG | 600 | 4 | 559 | 1.01 | GTGGTG | GA_GGG |
| yPred15 | GTATGAGTGGGTGTGGGC | 600 | 4 | 498 | 0.90 | TGGGTG | GTGTGG |
| yPred16 | GTCGTGGTGGTGGTGGGC | 600 | 4 | 470 | 0.85 | GTGGTG | GTG_TG |
| yPred17 | GAGGTGGTGGTGGTGGGG | 595 | 3 | 514 | 0.93 | GTGGTG | GTG_TG |
| yPred18 | TGTGGTGAGGGGGAGGGG | 665 | 3 | 53 | 0.10 | TGA_G | GGGG_A |
| yPred19 | GGTGTGGTGGTGGTGGGC | 580 | 4 | 65 | 0.12 | GTGGTG | GTG_TG |
| yPred20 | CGTGTGGGTTTGGGGGGG | 685 | 3 | 50 | 0.09 | GT_TGG | TTG_GG |

**Table S8: Test-tube investigation of randomly designed NCBs and G5.**

Ten randomly designed activators and a hypothetical bright candidate (G5) designed based on **Fig. 2** conclusion were evaluated in test tubes. Note that we do not preset any threshold of predicted success before selection here. On average, the enhancement ratio of the ten designs in yellow and red channels were 19 and 126, respectively. Since we selected top 30% (3,600) activator sequences as the bright class, we used the median enhancement ratio value from ranking 3,595 to ranking 3,600 sequences as our new criteria for bright/dark categorization, which corresponded to 145 and 66 for red and yellow channels, respectively (see **Table S9**). As a result, 1 out the 10 random sequences was identified as a "bright yellow" activator and 4 out of the 10 random sequences were identified as "bright red" activators.

| ID | Sequence (5' → 3') | Emission peak (nm) | Enhancement ratio (yellow) | Improvement ratio (compared to G15) | Enhancement ratio (red) | Improvement ratio (compared to G12) |
|---|---|---|---|---|---|---|
| G5 | CCCCCCGCGGGGTTTCCC | 645 | 39 | 0.09 | 83 | 0.19 |
| rand1 | AGGGACTAGGTGGGCGCT | 660 | 9 | 0.02 | 44 | 0.10 |
| rand2 | CGCGTGAGCGAGGTCGAG | 630 | 10 | 0.02 | 9 | 0.02 |
| rand3 | GTACGGAGGTGAGCTTGG | 660 | 23 | 0.04 | 66 | 0.15 |
| rand4 | TGTGCACAAGAGGGGAGG | 685 | 30 | 0.05 | 250 | 0.57 |
| rand5 | GCTGATTGGGCGCTTGGG | 695 | 24 | 0.04 | 206 | 0.47 |
| rand6 | GGCCGACTTGTGGGTAGG | 675 | 24 | 0.04 | 92 | 0.21 |
| rand7 | TGAGGGCTGAGACGCCGG | 660 | 19 | 0.03 | 53 | 0.12 |
| rand8 | GCTCGGGCCAGGTGGAAG | 625 | 68 | 0.12 | 79 | 0.18 |
| rand9 | AGTGGGGATGAGTGTGCA | 665 | 28 | 0.05 | 316 | 0.72 |
| rand10 | GCCGGGTTGTAGATGGGT | 670 | 18 | 0.03 | 149 | 0.34 |

**Table S9: Test-tube investigation of red and yellow NCBs ranked near 3600.**

As we selected top 30% (3,600) activator sequences as the bright class, we used the median enhancement ratio value from ranking 3,595 to ranking 3,600 sequences as our new criteria for bright/dark categorization, which corresponded to 145 and 66 for red and yellow channels, respectively.

**a Red channel**

| ID | Sequence (5' → 3') | Emission peak (nm) | Enhancement ratio (red) | Improvement ratio (compared to G12) |
|---|---|---|---|---|
| Rank3596 | CTCGAAGGTGGGGTGGGG | 650 | 83 | 0.19 |
| Rank3597 | TGGAAAGGTGGGGTGGGG | 600 | 239 | 0.54 |
| Rank3598 | CGTAGTGGTGGGGTGGGG | 670 | 233 | 0.53 |
| Rank3599 | GAACCCGGTGGGGTGGGG | 570 | 143 | 0.33 |
| Rank3600 | GGGTGGGGTGGGGGTGGA | 550 | 145 | 0.33 |

**b Yellow channel**

| ID | Sequence (5' → 3') | Emission peak (nm) | Enhancement ratio (yellow) | Improvement ratio (compared to G15) |
|---|---|---|---|---|
| Rank3596 | GGGTGGGGTGGGTCAATC | 655 | 55 | 0.10 |
| Rank3597 | CGAAGCGGTGGGGTGGGG | 600 | 210 | 0.38 |
| Rank3598 | AAACCGGGTGGGGTGGGG | 685 | 66 | 0.12 |
| Rank3599 | GGGTGGATGGCAGTGGGG | 590 | 61 | 0.11 |
| Rank3600 | GGGTGGTGCAGCGTGGGG | 615 | 182 | 0.33 |

**Table S10: Test-tube investigation of red POT candidates.**

Based on chip selection results, 9 sets of red POT candidates are evaluated in test tubes. All these candidates have their POT difference ratios greater than 1.63, with the largest difference ratio of 8.92 (rPOT5/rPOT6, **Fig. S18**). Single-nucleotide differences in these POTs are marked in red. POT difference ratio is simply the ratio of the enhancement ratios of the twins, which is,

POT difference ratio=(Enhancement ratio of bright candidate)/(Enhancement ratio of dark candidate)

| **ID (Bright)** | **Sequence (5' → 3')** | **ID (Dark)** | **Sequence (5' → 3')** | **POT difference ratio in red channel (Bright/Dark)** |
|---|---|---|---|---|
| rPOT1 | ATCCGTGGTGGGTGGGG | rPOT2 | ATTCGTGGTGGGTGGGG | 4.43±0.68 |
| rPOT3 | TCCATTGGTGGGTGGGG | rPOT4 | TTCATTGGTGGGTGGGG | 6.55±0.92 |
| rPOT5 | AATCCTGGTGGGTGGGG | rPOT6 | AATTCTGGTGGGTGGGG | 8.91±1.31 |
| rPOT7 | TCCATAGGTGGGTGGGG | rPOT8 | TGCATAGGTGGGTGGGG | 3.10±0.55 |
| rPOT7 | TCCATAGGTGGGTGGGG | rPOT9 | TCAATAGGTGGGTGGGG | 8.32±1.81 |
| rPOT3 | TCCATTGGTGGGTGGGG | rPOT10 | TACATTGGTGGGTGGGG | 3.10±0.39 |
| rPOT3 | TCCATTGGTGGGTGGGG | rPOT11 | TCGATTGGTGGGTGGGG | 3.39±1.11 |
| rPOT1 | ATCCGTGGTGGGTGGGG | rPOT12 | ATCAGTGGTGGGTGGGG | 2.82±1.34 |
| rPOT13 | ATCCGAGGTGGGTGGGG | rPOT14 | ATCGGAGGTGGGTGGGG | 1.63±0.20 |

| **ID (Bright)** | **Sequence (5' → 3')** | **ID (Dark)** | **Sequence (5' → 3')** | **POT difference ratio in yellow channel (Bright/Dark)** |
|---|---|---|---|---|
| rPOT1 | ATCCGTGGTGGGTGGGG | rPOT2 | ATTCGTGGTGGGTGGGG | 0.73±0.07 |
| rPOT3 | TCCATTGGTGGGTGGGG | rPOT4 | TTCATTGGTGGGTGGGG | 1.46±0.24 |
| rPOT5 | AATCCTGGTGGGTGGGG | rPOT6 | AATTCTGGTGGGTGGGG | 2.78±0.24 |
| rPOT7 | TCCATAGGTGGGTGGGG | rPOT8 | TGCATAGGTGGGTGGGG | 0.77±0.30 |
| rPOT7 | TCCATAGGTGGGTGGGG | rPOT9 | TCAATAGGTGGGTGGGG | 1.65±0.20 |
| rPOT3 | TCCATTGGTGGGTGGGG | rPOT10 | TACATTGGTGGGTGGGG | 1.19±0.11 |
| rPOT3 | TCCATTGGTGGGTGGGG | rPOT11 | TCGATTGGTGGGTGGGG | 1.52±0.40 |
| rPOT1 | ATCCGTGGTGGGTGGGG | rPOT12 | ATCAGTGGTGGGTGGGG | 1.59±0.57 |
| rPOT13 | ATCCGAGGTGGGTGGGG | rPOT14 | ATCGGAGGTGGGTGGGG | 0.51±0.05 |

**Table S11: Test-tube investigation of yellow POT candidates.**

Based on chip selection results, 9 sets of yellow POT candidates are evaluated in test tubes. All these candidates have their POT difference ratios greater than 3.29, with the largest difference ratio of 31.25 (yPOT5/yPOT6 NCBs, **Fig. S19**). Single-nucleotide differences in these POTs are marked in red.

| ID (Bright) | Sequence (5' → 3') | ID (Dark) | Sequence (5' → 3') | POT difference ratio in yellow channel (Bright/Dark) |
|---|---|---|---|---|
| yPOT1 | TAAGTGGGTGGGGTGGGG | yPOT2 | TAACTGGGTGGGGTGGGG | 9.16±1.65 |
| yPOT3 | TTAGTGGGTGGGGTGGGG | yPOT4 | TTAGTCGGTGGGGTGGGG | 9.41±0.69 |
| yPOT5 | CAGTGAGGTGGGGTGGGG | yPOT6 | CAGTCAGGTGGGGTGGGG | 31.25±5.37 |
| yPOT7 | AGCTGAGGTGGGGTGGGG | yPOT8 | AGCTAAGGTGGGGTGGGG | 14.17±2.95 |
| yPOT9 | ACAGTGGGTGGGGTGGGG | yPOT10 | ACAGCGGGTGGGGTGGGG | 6.15±1.20 |
| yPOT11 | ACAGTGGGTGGGGTGGGG | yPOT12 | ACAGAGGGTGGGGTGGGG | 3.29±0.26 |

| ID (Bright) | Sequence (5' → 3') | ID (Dark) | Sequence (5' → 3') | POT difference ratio in red channel (Bright/Dark) |
|---|---|---|---|---|
| yPOT1 | TAAGTGGGTGGGGTGGGG | yPOT2 | TAACTGGGTGGGGTGGGG | 0.88±0.07 |
| yPOT3 | TTAGTGGGTGGGGTGGGG | yPOT4 | TTAGTCGGTGGGGTGGGG | 0.69±0.24 |
| yPOT5 | CAGTGAGGTGGGGTGGGG | yPOT6 | CAGTCAGGTGGGGTGGGG | 2.10±0.34 |
| yPOT7 | AGCTGAGGTGGGGTGGGG | yPOT8 | AGCTAAGGTGGGGTGGGG | 0.83±0.15 |
| yPOT9 | ACAGTGGGTGGGGTGGGG | yPOT10 | ACAGCGGGTGGGGTGGGG | 0.84±0.07 |
| yPOT11 | ACAGTGGGTGGGGTGGGG | yPOT12 | ACAGAGGGTGGGGTGGGG | 0.68±0.05 |

**Table S12: Machine learning model prediction results**

In this report, we evaluated the predictability across various machine learning models, including logistic regression[17] (LR), linear discriminant analysis[18] (LDA), decision tree[19] (DT), AdaBoost[20] (ADA), and support vector machines[21] (SVM). We observed that after feature selection using Weka, LR revealed the best predictability for both the red channel (accuracy: 0.87; marked in red) and yellow channel (accuracy: 0.89; marked in yellow) based on 5-fold cross validation.

| Model (red) | Accuracy (Acc) | Sensitivity | Specificity | Positive prediction rate | Negative prediction rate |
|---|---|---|---|---|---|
| **LR** | 0.87 | 0.88 | 0.85 | 0.90 | 0.83 |
| **LDA** | 0.86 | 0.86 | 0.85 | 0.90 | 0.81 |
| **DT** | 0.84 | 0.86 | 0.80 | 0.86 | 0.80 |
| **ADA** | 0.86 | 0.87 | 0.85 | 0.90 | 0.83 |
| **SVM** | 0.86 | 0.86 | 0.85 | 0.90 | 0.81 |

| Model (yellow) | Accuracy (Acc) | Sensitivity | Specificity | Positive prediction rate | Negative prediction rate |
|---|---|---|---|---|---|
| **LR** | 0.89 | 0.88 | 0.90 | 0.92 | 0.85 |
| **LDA** | 0.87 | 0.85 | 0.90 | 0.92 | 0.82 |
| **DT** | 0.83 | 0.84 | 0.82 | 0.86 | 0.79 |
| **ADA** | 0.87 | 0.86 | 0.88 | 0.90 | 0.83 |
| **SVM** | 0.88 | 0.87 | 0.90 | 0.92 | 0.84 |

**Table S13: selected bright and dark features for yellow channel**

We defined the top 30% as the bright class and bottom 30% as the dark class. The feature extraction was processed using MERCI. We then used Weka to selected important features. The attribute evaluator was set to "CfsSubsetEval"[22] and the search method was set to "GreedyStepwise"[15]. All other parameters were set to default values.

**a** Selected bright features (the number indicates the segment that the motif belongs to)

| 3-nt | 4-nt | 5-nt | 6-nt |
|---|---|---|---|
| | AAGG_1 | AG_AG_1 | AAG_GT_1 |
| | AGGG_1 | CG_AG_1 | AAG_GT_3 |
| | AGGG_3 | GAGGT_1 | CGGG_T_1 |
| | ATGG_1 | G_AAG_1 | GAGG_G_2 |
| | CGGG_1 | G_ATG_1 | GA_GGG_2 |
| | GAGG_1 | GC_AG_1 | GA_GGG_3 |
| | GCGG_1 | GCC_G_3 | GA_TGG_1 |
| | GTGT_1 | GCT_G_3 | G_AGTG_1 |
| | TAGG_1 | G_CTG_1 | GCG_GG_2 |
| | TTGG_1 | GGGGG_2 | GC_TGG_1 |
| | | GT_AG_1 | G_CGGT_1 |
| | | GTT_G_1 | GGA_GG_1 |
| | | G_TAG_1 | GGC_GG_1 |
| | | G_TGT_1 | GGC_GG_2 |
| | | TGA_G_1 | GGGA_A_1 |
| | | TGA_G_2 | GGGA_G_1 |
| | | TG_AG_1 | GGGG_A_1 |
| | | T_GCG_3 | GGGG_A_3 |
| | | TT_GG_1 | GG_GCG_1 |
| | | | G_GGCA_1 |
| | | | GTGGTG_2 |
| | | | GTGTGG_3 |
| | | | GTG_TG_1 |
| | | | GTG_TG_3 |
| | | | GT_TGG_1 |
| | | | GT_TGG_2 |
| | | | TGGGTG_1 |
| | | | TGGGTG_2 |
| | | | TG_GCG_1 |
| | | | TTG_GG_2 |
| | | | TTG_GG_3 |
| | | | TT_GGT_3 |

**b** Selected dark features (the number indicates the segment that the motif belongs to)

| 3-nt | 4-nt | 5-nt | 6-nt |
|---|---|---|---|
| AAT_2 | ACGT_3 | AA_CG_2 | AA_TGG_1 |
| ACT_2 | CCGG_1 | AA_TG_1 | AA_TGG_3 |
| ATC_1 | GGAT_1 | A_AGT_1 | A_AGTG_1 |
| ATC_2 | GGTT_1 | AC_CG_2 | A_AGTG_2 |
| ATT_2 | TCGG_1 | AC_TG_1 | ACG_GG_1 |
| CAT_2 | | AT_CG_2 | AC_GGT_1 |
| CGC_2 | | AT_CG_3 | AC_GTG_3 |
| CTC_2 | | ATG_T_1 | AC_TGG_1 |
| CTT_3 | | AT_TG_1 | A_CGGT_1 |
| TCA_3 | | C_AGT_1 | ATG_TG_1 |
| TCT_2 | | C_AGT_2 | AT_GGT_1 |
| TTC_2 | | CCG_T_1 | AT_GTG_1 |
| TTT_3 | | CC_TG_3 | CAG_TG_1 |
| | | CTC_G_2 | CA_GGT_1 |
| | | CT_TG_1 | CA_TGG_1 |
| | | C_TGT_1 | C_AGGT_1 |
| | | C_TGT_2 | C_AGTG_2 |
| | | GAT_G_1 | CCG_TG_1 |
| | | GT_CT_1 | CC_GGT_1 |
| | | GT_CT_3 | CC_TGG_1 |
| | | GT_TC_1 | CGTGGG_1 |
| | | G_TCG_1 | CGTGGG_2 |
| | | TA_TG_1 | CT_GGT_1 |
| | | TA_TG_3 | CT_TGG_1 |
| | | T_AGT_1 | C_TGGT_1 |
| | | TC_GT_3 | C_TGTG_1 |
| | | TC_TG_1 | C_TGTG_2 |
| | | TG_AT_3 | G_CGTG_1 |
| | | TG_TT_1 | G_CGTG_2 |
| | | TG_TT_3 | GGG_CC_1 |
| | | T_GTT_1 | GGGT_A_1 |
| | | TT_CG_3 | GGGT_C_1 |
| | | T_TGT_1 | GGGT_T_1 |
| | | | GG_GTC_1 |
| | | | GGTGGA_1 |
| | | | GGTGGA_2 |
| | | | G_GGAA_1 |
| | | | G_GGAC_1 |
| | | | G_GGAT_1 |
| | | | G_GGCT_1 |
| | | | G_GGTA_1 |
| | | | G_GGTC_1 |
| | | | G_GGTT_1 |
| | | | T_AGTG_2 |
| | | | TCGG_G_1 |
| | | | TCG_GG_1 |
| | | | TC_TGG_1 |
| | | | T_CGGT_1 |
| | | | TGGA_A_1 |
| | | | TGGA_C_3 |
| | | | TGGA_G_1 |
| | | | TGG_AA_1 |
| | | | TGG_CT_1 |
| | | | TGGT_A_1 |
| | | | TGGT_A_3 |
| | | | TGGT_C_1 |
| | | | TGGT_T_3 |
| | | | TGG_TT_2 |
| | | | TG_GAA_1 |
| | | | TG_GAC_1 |
| | | | TG_GAT_1 |
| | | | TG_GCC_1 |
| | | | TG_GTA_1 |
| | | | TG_GTC_1 |
| | | | TG_GTT_1 |
| | | | T_GGAC_1 |
| | | | T_GGCC_1 |
| | | | T_GGCT_1 |
| | | | T_GGTA_1 |
| | | | TTG_TG_1 |

**Table S14: selected bright and dark features for red channel**

We defined the top 30% as the bright class and bottom 30% as the dark class. The feature extraction was processed using MERCI. We then used Weka to selected important features. The attribute evaluator was set to “CfsSubsetEval” and the search method was set to “GreedyStepwise”. All other parameters were set to default values.

**a** Selected bright features (the number indicates the segment that the motif belongs to)

| 3-nt | 4-nt | 5-nt | 6-nt |
|---|---|---|---|
| CCC_1 | AAGG_1 | AA_GG_1 | AA_GGT_2 |
| CCC_3 | ATGG_1 | AA_GG_3 | ACG_GT_1 |
| | CTGG_3 | A_AGG_3 | ACG_GT_3 |
| | GAGG_1 | ACA_G_1 | AC_GGT_2 |
| | GCGG_1 | AC_AG_3 | AC_GGT_3 |
| | TAGG_1 | ACC_G_1 | AG_GGT_1 |
| | TTGG_1 | AC_GG_2 | AG_GGT_2 |
| | | ACT_G_1 | ATG_GT_3 |
| | | ACT_G_3 | A_TGGT_2 |
| | | A_CCG_1 | A_TGGT_3 |
| | | A_CCG_3 | CAG_GT_3 |
| | | A_CGG_1 | CCG_GT_3 |
| | | A_CGG_3 | CC_GGT_2 |
| | | AT_GG_1 | CG_GGT_2 |
| | | CAC_G_1 | CT_GGT_2 |
| | | CAC_G_3 | GA_GGT_2 |
| | | CA_CG_1 | GA_GGT_3 |
| | | CA_CG_3 | G_AGGT_1 |
| | | CA_GG_1 | GC_GGT_2 |
| | | CA_GG_2 | GC_GGT_3 |
| | | C_AGG_2 | GGGG_C_3 |
| | | CCA_G_1 | GGG_GC_1 |
| | | CC_AG_3 | GGG_GC_2 |
| | | CC_CG_3 | GTGGTG_1 |
| | | CC_GG_1 | GTGGTG_3 |
| | | CCT_G_1 | TAG_GT_3 |
| | | CCT_G_3 | TA_GGT_2 |
| | | C_CAG_3 | TC_GGT_1 |
| | | C_CGG_1 | TC_GGT_3 |
| | | C_CGG_2 | TGGGGG_1 |
| | | C_CTG_1 | TGGGGG_3 |
| | | C_CTG_3 | TGGGTG_1 |
| | | C_GGG_1 | TGGGTG_3 |
| | | CT_AG_1 | TTGG_G_2 |
| | | CT_AG_3 | TTG_GT_3 |
| | | CTC_G_1 | T_TGGT_2 |
| | | CTC_G_3 | T_TGGT_3 |
| | | CT_CG_1 | |
| | | CT_CG_3 | |
| | | CT_GG_2 | |
| | | GCC_G_1 | |
| | | GCC_G_3 | |
| | | GC_GG_3 | |
| | | GGGAC_3 | |
| | | GGGGA_1 | |
| | | GGGGA_2 | |
| | | GGGGA_3 | |
| | | GGGGG_1 | |
| | | TA_GG_3 | |
| | | T_AGG_3 | |
| | | TCC_G_1 | |
| | | TC_CG_1 | |
| | | TC_CG_3 | |
| | | TC_GG_3 | |
| | | T_CCG_3 | |
| | | TT_GG_1 | |
| | | TT_GG_3 | |

**b** Selected dark features (the number indicates the segment that the motif belongs to)

| 3-nt | 4-nt | 5-nt | 6-nt |
|---|---|---|---|
| AAA_1 | CCGT_1 | AA_TG_1 | AAGT_G_1 |
| AGA_1 | CGTG_3 | AGT_G_1 | AA_GTG_2 |
| AGA_3 | GAGT_1 | AGT_G_3 | ACG_GG_2 |
| ATA_1 | GGAA_1 | A_TGT_1 | AC_TGG_1 |
| TAA_1 | GGAG_1 | CGT_G_1 | AC_TGG_2 |
| TGA_1 | GGAT_1 | CGT_G_2 | A_TGGG_2 |
| | GGCA_1 | C_TGT_1 | CAG_GG_1 |
| | GGCC_1 | GAA_C_1 | CAG_GG_2 |
| | GGCT_1 | GAA_G_3 | CAGT_G_1 |
| | GGTA_1 | GA_CT_1 | CA_TGG_2 |
| | GGTT_1 | G_AGC_1 | CG_GGG_1 |
| | | GCA_G_3 | CG_GGG_2 |
| | | GCT_G_3 | GCG_TG_1 |
| | | GC_TC_1 | GCG_TG_2 |
| | | G_GCC_1 | GC_TGG_1 |
| | | G_GTC_1 | GC_TGG_2 |
| | | GTA_G_1 | GGGA_A_1 |
| | | GT_AT_1 | GGG_CG_1 |
| | | GT_CA_1 | GGGT_A_1 |
| | | GT_CC_1 | GG_GTA_2 |
| | | GT_CT_1 | GG_GTT_2 |
| | | GTG_C_2 | GGTGGT_1 |
| | | GTGGC_1 | GGTGGT_2 |
| | | GT_GA_2 | GGTG_T_1 |
| | | G_TTA_1 | GGT_TC_1 |
| | | G_TTG_1 | GTG_AT_1 |
| | | TAGTG_1 | GTGG_C_2 |
| | | TC_TG_1 | GTGGGC_3 |
| | | TC_TG_2 | GTG_TA_1 |
| | | TG_AA_3 | GT_GTA_1 |
| | | TG_CA_1 | GT_GTT_1 |
| | | TG_CG_1 | TAG_GG_2 |
| | | TG_CT_1 | TGG_AA_2 |
| | | TGTGG_2 | TGG_AC_2 |
| | | TG_TA_3 | TGG_AC_3 |
| | | T_GAA_1 | TGG_AG_2 |
| | | T_GAC_3 | TGG_AT_2 |
| | | T_GAG_3 | TGGGTT_1 |
| | | T_GAT_3 | TGGGTT_3 |
| | | T_GCT_1 | TGGT_A_1 |
| | | T_GTA_1 | TGG_TC_3 |
| | | T_GTC_1 | TG_GAC_1 |
| | | T_GTC_3 | TG_GAG_1 |
| | | TT_AG_1 | TG_GCC_1 |
| | | TTC_G_3 | TG_GCG_3 |
| | | TT_TG_1 | T_GGAA_3 |
| | | | T_GGTA_3 |
| | | | T_TGGG_2 |
| | | | T_TGGG_3 |
